# Early Increases in Hippocampal Synchrony Predict Chronic Epileptiform Burden in a Temporal Lobe Epilepsy Model

**DOI:** 10.64898/2026.11.04.763700

**Authors:** Nadia Gasmi, Hale Tobin, Delnia Ahmadpour, Ghazaal Tahmasebi, Amirhossein Koohfar, Sam Kalnitsky, Morgan Rogers-Carter, Jacob Icardi, Sarah Roehrs, Sierra Bhatia, Hannah Shindler, Jordon Manworren, Priyanka Agochiya, Shriya Singh, Sarah Stanko, Catherine Richards, Hoonkyo Suh, Andreas Alexopoulos, Demitre Serletis, Imad Najm, Hod Dana, Dimitrios Davalos

## Abstract

Temporal lobe epilepsy (TLE), the most common focal epilepsy, is often drug-resistant and arises predominantly in the hippocampus. Disease severity is conventionally assessed by seizure frequency and duration, yet epileptiform activity outside discrete seizures may also track disease progression. Whether this activity reflects hippocampal network reorganization as part of epileptogenesis, and whether such reorganization predicts chronic severity, is unknown. Here, we combined longitudinal two-photon calcium imaging of hippocampal CA1 neurons with repeated 24-hour EEG recordings that quantified epileptiform burden in the same pilocarpine-treated mice over 16 weeks. CA1 firing and synchrony rose acutely, subsided during a latent period, and increased again chronically, with synchronized firing organized into brief waves that propagated in a consistent direction. Epileptiform burden followed a similar temporal trajectory and increasingly contained electrographic seizures. CA1 synchrony correlated with epileptiform burden and stratified mice into groups with distinct neuronal and electrographic outcomes. CA1 synchrony measured within three weeks of pilocarpine induction predicted chronic epileptiform burden, identifying early hippocampal synchrony as a candidate imaging biomarker and pointing to an early window for intervention in TLE.

## Introduction

Epilepsy affects approximately 52 million people worldwide^1^. Despite the availability of numerous antiseizure medications, approximately 15% of adults and 25% of children with epilepsy develop drug-resistant disease, and more than one third of patients seen at specialist clinics are drug-resistant^2^. Temporal lobe epilepsy (TLE), the most common focal epilepsy, profoundly impairs quality of life through cognitive deficits and other comorbidities^3, 4, 5^.

Epileptogenesis, the process whereby a normal brain becomes predisposed to spontaneous seizures^6^, offers a window for preventive therapeutic intervention. It unfolds in phases: an acute phase (hours to days) after the initial injury, characterized by status epilepticus (SE) and neuronal damage; a latent period (months to years in patients, but days to weeks in rodent models) during which the brain appears seizure-free while its networks reorganize; and a chronic phase marked by spontaneous recurrent seizures (SRS)^7, 8, 9, 10^. Selective loss of GABAergic interneurons^11, 12, 13^, reduced inhibition and the formation of recurrent excitatory circuits through synaptic reorganization and axonal sprouting^7, 8, 14^ create hyperexcitable networks capable of the hypersynchronous activity that characterizes seizures^7, 15, 16^. Yet it is currently not possible to predict early in this process who will develop severe epilepsy, partly because no reliable biomarker identifies, soon after the initial injury, who will progress to chronic, severe seizures^17^.

How disease severity should be measured is itself an open question. Epilepsy is commonly characterized by seizure frequency, duration and severity, yet seizure counts fluctuate substantially over time independent of treatment^18^, patient diaries are often incompletely kept^19^, and seizure frequency becomes an unreliable index of seizure propensity when seizures are infrequent^20^. Seizure-based outcomes can also depend on how a seizure is defined: for example, in the intrahippocampal kainate mouse model, whether carbamazepine appeared effective depended on both mouse strain and the EEG criteria used to define a seizure^21^. Importantly, pathological activity that falls outside the definition of discrete seizures carries substantial information about disease state. In patients, interictal epileptiform activity follows circadian and multidien rhythms, and seizures tend to cluster on their rising phase^22^. The burden of epileptiform activity, the proportion of recording time occupied by epileptiform patterns irrespective of seizure event numbers, also correlates with outcome: across 1,967 acutely ill patients, the probability of poor neurological outcome rose with peak epileptiform burden^23^. Clinical terminology likewise defines electrographic SE by the time occupied by seizure activity (≥10 continuous minutes, or ≥20% of any 60-min period) rather than by an event count^24^. In animal models, interictal activity correlates with disease stage: in pilocarpine-treated rats, a shift between two interictal discharge types marks the transition from the latent to the chronic phase^25^; after kainate-induced SE, the frequency of interictal spikes in the preceding days correlates with how quickly epilepsy develops^26^; and in a mouse model of childhood epilepsy, interictal features predict where seizures arise and how severe the seizure burden becomes^27^. Whether these broader measures of epileptiform activity reflect changes in neuronal network activity at cellular resolution, and whether they can be predicted from cellular-level measurements made early after the initial injury, remains unknown.

TLE is closely linked to hippocampal dysfunction: in mesial TLE, its most common form, approximately 80% of seizures arise in the hippocampus^28^. Yet how hippocampal neurons behave during epileptogenesis remains poorly understood. Seizure development in animal models is usually monitored by behavioral scoring, such as the Racine scale^29^, or by EEG^30, 31^, readouts that lack cellular resolution and cannot resolve what electrographic measures correspond to at the level of individual neurons. To bridge this gap, we combined longitudinal EEG with two-photon laser scanning microscopy (TPLSM) of hippocampal neurons expressing a genetically encoded calcium indicator (GECI), which resolves single-cell activity while capturing population dynamics^32, 33, 34, 35^, in the pilocarpine mouse model. This model recapitulates key features of human TLE, including GABAergic interneuron loss^11, 12^ and progression from SE through a latent period to chronic epilepsy with SRS and hippocampal sclerosis^36, 37^, and is well suited to longitudinal studies of disease progression^38^.

By imaging CA1 excitatory neurons and recording EEG in the same mice across the full epileptogenic trajectory, we found that pilocarpine induction drove rapid increases in CA1 synchrony within three days, carried by population events that propagated through CA1 as directionally consistent waves. EEG data, analyzed to quantify the burden of prolonged epileptiform discharge episodes rather than to detect seizures, revealed pathology that peaked after induction, declined during the latent period and remained elevated through the chronic phase. CA1 synchrony correlated with this burden in individual mice. Synchrony changes within the first three weeks after SE predicted chronic epileptiform burden 7–16 weeks after SE, identifying early hippocampal synchrony as a candidate biomarker and potential therapeutic target for TLE.

## Results

### A refined pilocarpine model for longitudinal hippocampal imaging and EEG recordings in the same mice

Transgenic mice expressing the red GECI jRGECO1a primarily in excitatory neurons^39^ were implanted with cortical and hippocampal EEG electrodes and a chronic cranial window over the contralateral CA1 (**Fig. 1a-i**). After a ∼5-week recovery that allowed glial responses to subside (**Supp. Fig. 1**) and windows to clear, mice were acclimated to a treadmill and randomly assigned to pilocarpine or vehicle. Our pilocarpine protocol combined mortality-reducing pretreatments adapted from Botterill et al.^40^ with sex-specific pilocarpine dosing and severity-triggered diazepam post-treatments (**Fig. 1a-ii**; Methods); this protocol, including a second diazepam dose when seizures persisted, achieved 100% survival in both sexes (**Fig. 1b**), a marked improvement compared to the substantially higher mortality reported previously^36, 41, 42^. These refinements maintained high induction success, as every treated mouse reached the SE threshold within 180 minutes. Notably, sex-specific dosing did not alter the induction phenotype: males and females had similar cumulative seizure scores and latencies to Racine stage 4–5 (**Fig. 1c-d**). The numbers of mice recorded by TPLSM, EEG, or both, and included in each downstream analysis are summarized in **Fig. 1e** and **Supp. Table 1**.

**Figure 1.**
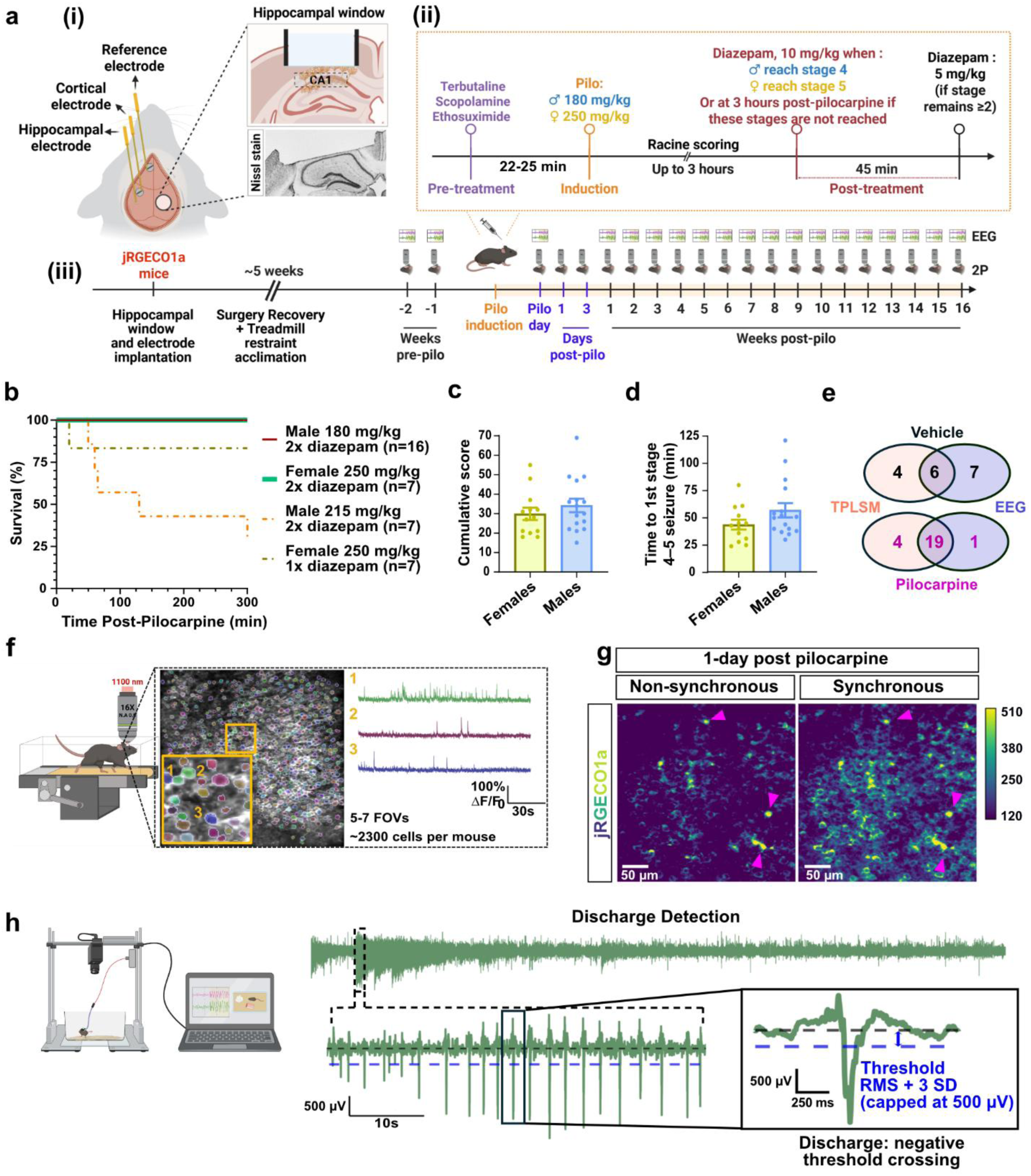
Longitudinal monitoring of epileptogenesis in the mouse brain. **(a)** Schematic of the experimental design. Thy1-jRGECO1a mice were implanted with a hippocampal cranial window for TPLSM imaging and hippocampal, cortical, and reference electrodes for EEG recordings (i). A refined SE-induction protocol was developed that included mortality-reducing pretreatments while preserving SE, pilocarpine administration at sex-specific doses (ii; 250 mg/kg for females, 180 mg/kg for males), followed by diazepam administration as indicated to limit SE duration and mortality upon reaching Racine stage 4 (males) or stage 5 (females). The timeline (iii) illustrates surgery, recovery and treadmill acclimation, baseline recordings, drug induction, and post-pilocarpine recording sessions. **(b)** Kaplan–Meier survival curves for mice receiving the different pilocarpine and diazepam protocols. "2× diazepam" denotes protocols in which a second dose was administered when seizures of stage 2 or higher persisted after the first dose; "1× diazepam" denotes protocols without this provision. **(c-d)** Following pilocarpine-induced seizures, mice were monitored and scored using the Racine scale. The cumulative Racine score (c) and the time to reach a Racine score of 4 or 5 (d; time to first seizure) were quantified (n=29 mice; 16 males, 13 females). Racine scoring was completed prior to diazepam administration and includes all 29 mice that received the final pilocarpine doses and survived (one female given a single diazepam dose died), irrespective of diazepam regimen. **(e)** Venn diagrams showing the overlap between mice with usable TPLSM recordings and mice with usable EEG recordings, separately for Vehicle and Pilocarpine groups. For Vehicle mice: 4 had TPLSM data only, 7 had EEG data only, and 6 had both (paired TPLSM + EEG). For Pilocarpine mice: 4 had TPLSM data only, 1 had EEG data only, and 19 had both (paired TPLSM + EEG). **(f)** Imaging and recording setup for awake, head-fixed mice on a voluntary treadmill (left). TPLSM images of CA1 pyramidal cells (middle) were acquired from 5–7 FOVs across CA1. For each FOV, somata were segmented using Cellpose2 to extract fluorescence traces corresponding to single-cell activity, illustrated for three example neurons (right). **(g)** Example jRGECO1a fluorescence images from the same FOV illustrating non-synchronous and synchronous firing of CA1 neurons. Arrowheads indicate characteristic neuronal cells that were active during both recordings and served as fiducial marks for relocation purposes. **(h)** Example hippocampal EEG trace from a mouse 9 days after pilocarpine administration and detection of individual epileptiform discharges as negative threshold crossings (threshold: RMS + 3 × SD of the recording, capped at 500 µV for hippocampal and 400 µV for cortical electrodes). How discharges are aggregated into epileptiform discharge episodes (EDEs) is shown in Fig. 4b.

TPLSM recorded spontaneous activity in awake, head-fixed mice free to walk on a treadmill (**Fig. 1f**) at baseline and at defined daily and weekly intervals after pilocarpine induction (**Fig. 1a-iii**). Each session sampled 5–7 fields of view (FOVs) across CA1 encompassing ∼2,300 neurons per mouse. After semi-automated segmentation, calcium transients were extracted for each segmented neuronal soma (**Fig. 1f**) and action potential (AP) timing and number were estimated^43^. Following pilocarpine induction, synchronized activity appeared as flashes across the FOV, readily distinguishable from the largely asynchronous baseline activity (**Fig. 1g**, **Supp. Video 1**).

Video-EEG (vEEG) recordings from the same mice were collected at the same weekly intervals, starting with two baseline sessions (**Fig. 1a-iii**). After pilocarpine, characteristic epileptiform discharges (rapid voltage decreases with slower return to baseline) were detected in hippocampal and cortical channels by amplitude thresholding relative to each recording’s own baseline (**Fig. 1h**) and aggregated into a measure of epileptiform burden (see below and Methods). Together with the improved survival, this provides a reliable platform for following hippocampal network activity and epileptiform burden in the same mice across the epileptogenic arc.

### CA1 synchrony follows the acute, latent and chronic phases of epileptogenesis

The same CA1 FOVs were re-imaged after pilocarpine, on the induction day when the animal’s condition allowed, on days 1 and 3, and weekly thereafter (**Fig. 2a-b**; Methods). Average CA1 firing rate rose sharply in the days after pilocarpine (acute period, pilocarpine day to Week 3), declined during a latent period (Weeks 4–6) and rose again during the chronic period (Weeks 7–16), exceeding vehicle levels in the acute and chronic but not the latent phase (**Fig. 2c-d**). These periods were defined from the CA1 data, so the acute period used here extends beyond the hours-to-days window conventionally used after SE^7, 8, 9^. CA1 neurons in pilocarpine-treated mice also began to fire together, as illustrated by coincident spikes (red) in three example cells (**Fig. 2b, c**, left). Across the full FOV, this coincident firing formed brief, population-wide events that appeared as bright flashes (**Supp. Video 1**) and as vertical bands in raster plots (**Fig. 2c**, right), consistent with the pathological network synchrony described in epilepsy^16^. We termed these events Periods of Synchronized Activity (PSAs), detected when the number of APs within a five-frame window reached at least 25% of the number of neurons in the FOV (Methods, **Supp. Fig. 2**). PSAs recurred multiple times per recording, were prominent in the acute and chronic phases, were largely absent at baseline and during the latent period, and varied between chronic-phase sessions (**Fig. 2c**, 7 vs. 13 weeks).

**Figure 2.**
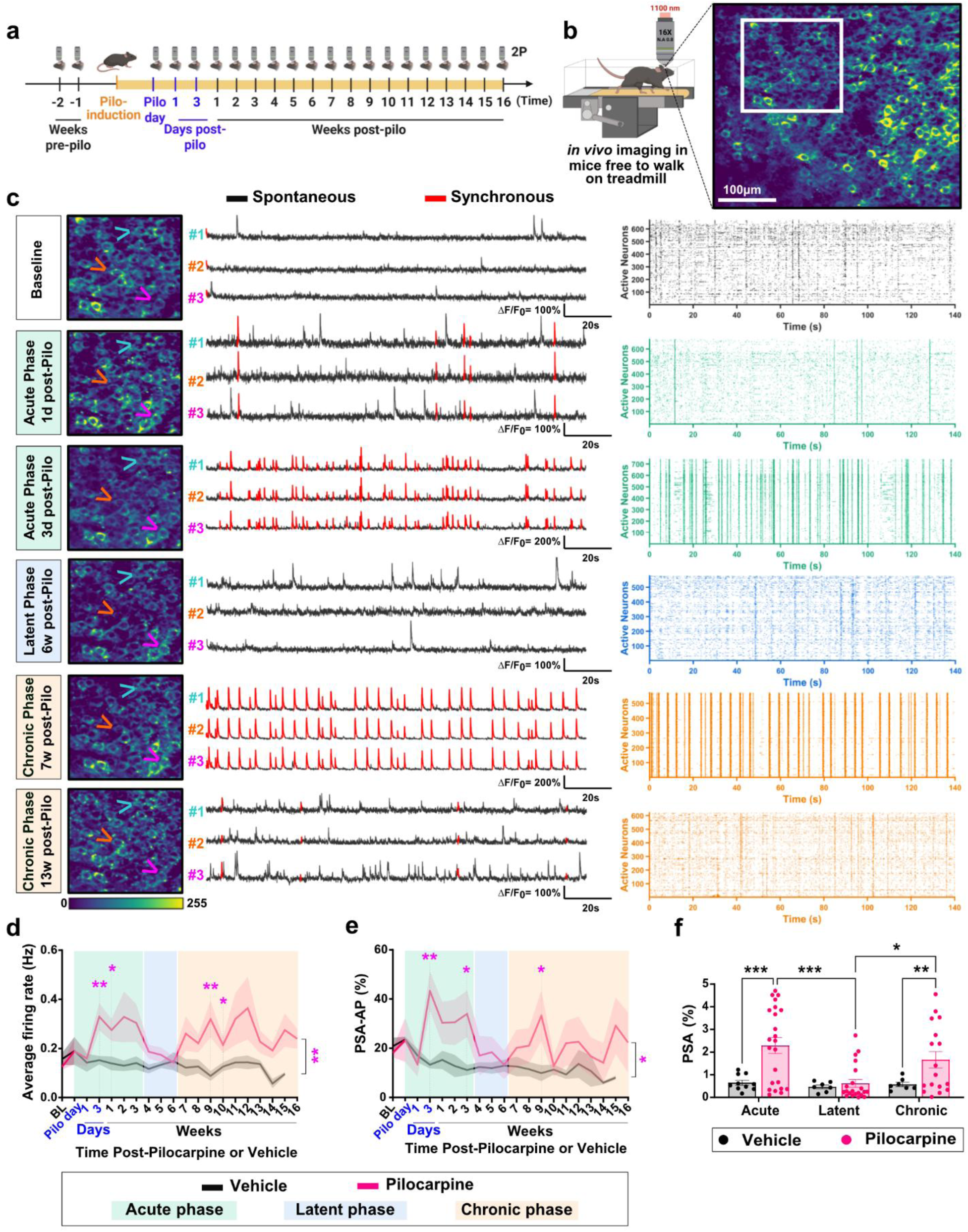
Longitudinal TPLSM imaging reveals rapid changes in neuronal synchronization following pilocarpine induction with long-term effects. **(a)** Experimental timeline of TPLSM imaging sessions. **(b)** Schematics of the experimental setup with an example TPLSM image of jRGECO1a-expressing CA1 pyramidal neurons. **(c)** Representative images from the FOV region shown in **b** across baseline, acute, latent, and chronic phases, together with activity traces from the same marked cells, illustrate same-cell monitoring and the emergence of pathological synchrony at the single-cell level. The timing of synchronized firing in the example cells is highlighted in red on the black activity traces. Raster plots (right) of all active neurons within the same respective FOVs further demonstrate the emergence and progression of synchronous events across all neurons (Periods of Synchronized Activity, PSAs), which appear as vertical bands. The 7- and 13-week chronic-phase sessions illustrate the session-to-session variability in PSA occurrence within the same FOV; group data are shown in **d–f**. **(d–e)** Longitudinal measurements of average neuronal firing rate (**d**) and of the proportion of AP firing occurring during PSA events, calculated relative to all APs recorded over each respective recording session in the CA1 network (PSA-AP; **e**). Both show a rapid increase within 3 days after pilocarpine induction, remaining elevated during the acute period, subsiding during the latent period, and rising again in the chronic period. **(f)** Summary data for PSA frequency (PSA%) during the acute, latent, and chronic phases show a significant increase in the pilocarpine group in the acute and chronic phases, but not in the latent phase. Data in **(d–e)** are presented as mean (solid lines) ± SEM (shaded areas). Shaded background regions denote disease phases: acute (day of pilocarpine to week 3, green), latent (weeks 4–6, blue), and chronic (weeks 7–16, orange). n=23 (13 males, 10 females) and 10 mice (5 males, 5 females) were included in the pilocarpine and vehicle groups at the study onset, respectively. See **Supp. Table 2** for mouse numbers in each subsequent recording week. Statistical analyses were performed using mixed-effects models with restricted maximum likelihood and Tukey’s HSD *post hoc* correction for multiple comparisons. *,p<0.05; **,p<0.01; ***,p<0.001. A complete list of significant comparisons is provided in **Supp. Table 2**.

We quantified synchrony as the percentage of recorded frames within PSAs (PSA%) and the fraction of all APs fired during PSAs (PSA-AP; **Supp. Fig. 2**). Like firing rate, both exceeded vehicle levels in the acute and chronic phases but not the latent phase (**Fig. 2e-f**, **Supp. Fig. 3a-b**), and neither synchrony nor firing rate differed by sex (**Supp. Fig. 3c-e**). Longitudinal recording from the same CA1 neurons thus captured a cellular-level hippocampal signature of the acute–latent–chronic progression previously described with EEG in the pilocarpine model^36, 38^, a course that resembles human TLE.

To examine individual animal progression trajectories, we classified mice as more- or less-affected (n = 10 and 7, respectively) according to whether PSA% exceeded three times its baseline value in at least two post-induction sessions and PSA-AP exceeded 80% in at least one (Methods; **Supp. Fig. 4a-b**). The more-affected group showed immediate and sustained increases in synchrony and firing in the acute and chronic periods, whereas the less-affected group did not differ from vehicle at any timepoint apart from a transient reduction in PSA-AP on the induction day (**Supp. Fig. 4c–j**), consistent with the substantial animal-to-animal variability of the pilocarpine model^42^. The trajectories diverged within the first week, well before the chronic phase, suggesting that the magnitude of acute CA1 disruption may foreshadow long-term severity. We therefore further examined the spatial and temporal properties of individual PSAs.

### Synchronized CA1 activity waves propagate along the posterior-to-anterior axis

PSAs were the predominant CA1 network pathology we identified, so we characterized them in detail. PSAs lasted approximately 100 ms (**Fig. 3a-b**). In a representative recording (**Fig. 3c-f**, **Supp. Video 2**), activity in most PSAs propagated as a wave from the posterior to the anterior side of the imaged FOV (**Fig. 3c**), and cellular firing rates during PSAs decreased along the same axis (**Fig. 3d**). Each neuron’s first-AP latency within PSAs was therefore negatively correlated with its firing rate, where more active neurons fired earlier. Higher firing rate was also associated with less temporal jitter (**Fig. 3e**). This recruitment order was not an artifact of measuring rate and latency over the same events: computed from non-overlapping halves of the PSAs in each recording, the correlation was only modestly weaker (mean Spearman ρ = −0.517 ± 0.014 vs. −0.621 ± 0.014 for non-overlapping vs. shared events, respectively; mean ± SEM, n = 189 recordings; **Supp. Fig. 5a-b**). The propagation followed the direction of decreasing firing rate during PSAs, but not outside PSAs (**Fig. 3f**).

**Figure 3.**
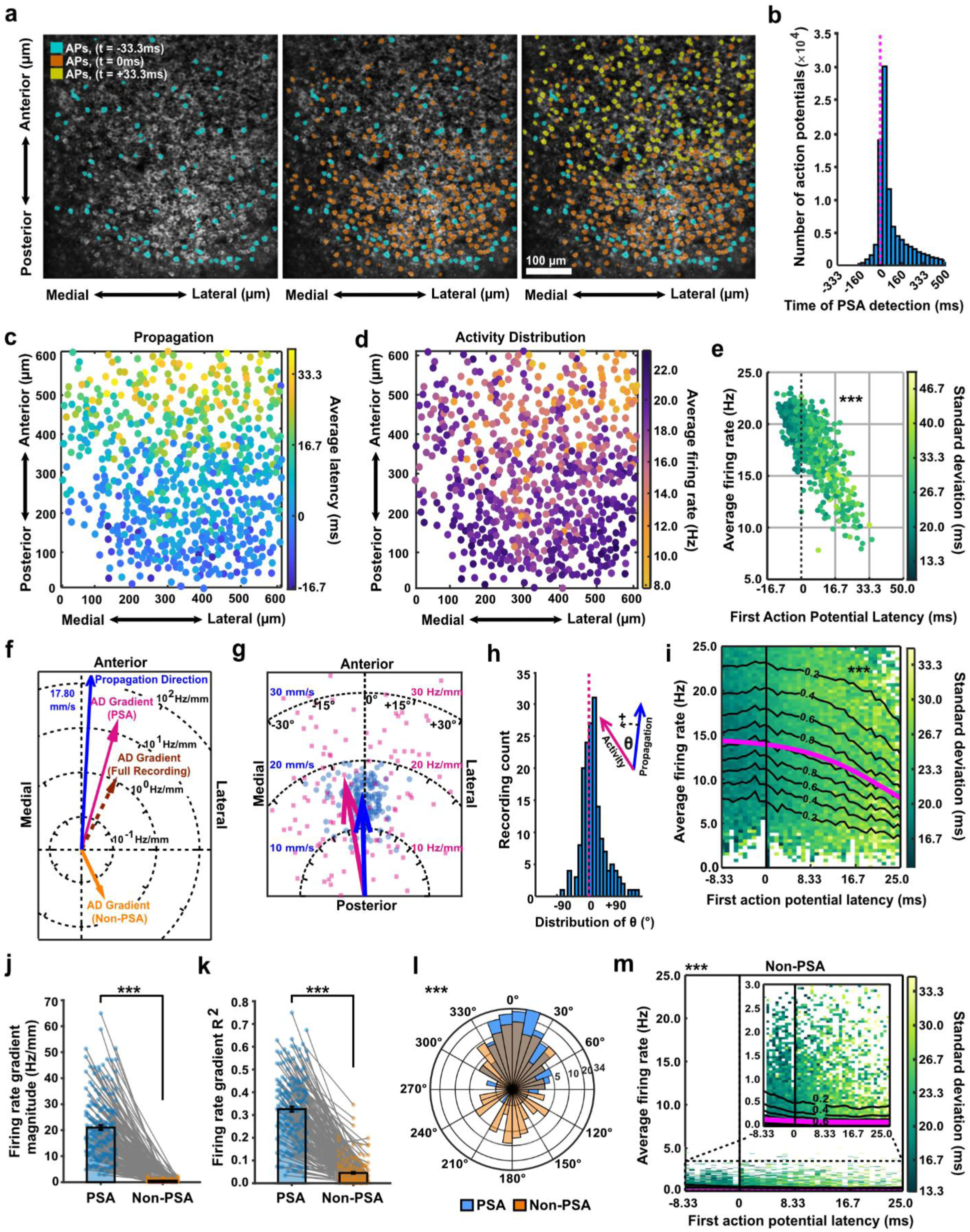
PSA events present wave-like propagation properties with activity-dependent topography and latency. **(a)** Example images from three consecutive frames of the same FOV taken before (left), at (middle), and after (right) crossing the PSA threshold. Neurons that fired their first AP in the left, middle, and right frames are labeled cyan, orange, and yellow, respectively, illustrating the wave-like propagation of a PSA event. **(b)** The distribution of all recorded APs during all analyzed PSA events centered around the frame of PSA onset shows the short duration of the PSA event (n=100,343 APs), each bin represents 33.3ms (1 frame). The vertical magenta line represents the location of the onset (zero) frame. **(c)** Map of the same FOV as in **a** showing the locations of 650 somata, with each neuron color-coded by the mean latency from PSA onset to the time of its first AP firing (mean across n=25 PSA events in a single recording; p<0.001, permutation test comparing the observed gradient magnitude against shuffled data; n=2000 permutations). **(d)** Map of the same FOV as in **a**, showing each neuron color-coded by its average firing rate across all PSA frames in the same recording shown in **c**. To characterize the spatial structure of firing-rate distributions, firing rates of all recorded neurons were fit with a plane. The direction of the gradient of this plane pointed from anterior to posterior, indicating that firing rates decreased along the posterior-to-anterior axis (p<0.001, permutation test comparing the observed gradient magnitude to shuffled data; n=2000 permutations). Firing rates were averaged across all PSA frames within a single recording. **(e)** Scatter plot of all neurons in the same FOV as in **a-d** (n=650) showing the relationship between each neuron’s average firing rate and its mean latency to fire its first AP within the detected PSA events. Neurons with higher firing rates tend to fire earlier in PSAs (Spearman’s correlation coefficient ρ=-0.7089, p=2.46 × 10^-100^). Dots are color-coded by the standard deviation of each cell’s latency, highlighting lower variability in neurons with higher firing rates. **(f)** Summary of activity distribution gradient directions and magnitudes (AD; for simplicity, the direction of decreased activity is shown) are shown during PSA, Non-PSA frames, and the entire recording of the same FOV in **a, c-e,** and with respect to the PSA propagation direction within the same FOV. PSA events when isolated produce the closest direction to propagation and present the largest gradient magnitude (logarithmic scale). **(g)** Summary of the directions and magnitudes of all fitted firing rate and propagation direction data (magenta squares and blue dots, respectively) across all recordings from all mice (n= 174 recordings from 14 mice). Arrows indicate the average across all respective data points. **(h)** Distribution of the angle between the PSA propagation and firing rate gradient from the same recordings for all analyzed recordings. The mean difference is +10.71° ± 3.14° (mean ± SEM; data from n=174 recordings). The definition of positive angle is shown with respect to the directions in **g**. **(i)** Relationship between each neuron’s mean first-AP latency within a PSA event and its average firing rate across all PSA events (n=56,791 neurons). Contour lines indicate the fraction of total neurons within the upper and lower bounds; the magenta line shows a sigmoidal function fit. Neurons with higher average firing rates tended to fire earlier within PSA events (Spearman’s ρ=−0.2740, p<0.0001). Dots are color-coded by the standard deviation of each neuron’s latency, highlighting larger variability in neurons with lower firing rates that fire late in the PSA events. **(j-k)** Average firing-rate data during PSA and non-PSA frames were each fit by a plane. Compared to PSA frames, non-PSA frames showed a significant decrease in both the gradient magnitude of the fitted plane (**j**) and the goodness of fit (R²; **k**), indicating that the spatially structured differences in firing rate across neurons largely disappear outside of PSA events (p<0.0001 for both comparisons; n=174 recordings; paired two-tailed Student’s t-test). **(l)** The distributions of firing-rate gradient angles during PSA and non-PSA frames from the same recordings were significantly different, showing a predominantly anterior-pointing direction during PSA frames and a broader, more uniform distribution during non-PSA frames (p=0.0006; n=174 recordings; two-sample Kolmogorov–Smirnov test; angles as in **h**; logarithmic scale). **(m)** Comparison of the relationship between each neuron’s mean first-AP latency within a PSA event and its average firing rate during non-PSA frames from the same recordings and neurons, using the same representation as in **i**. During non-PSA frames, the relationship between firing rate and latency changed substantially (Spearman’s ρ=−0.17, p<0.0001). *,p<0.05; **,p<0.01, and ***,p<0.001.

Across the full dataset, these features were reproducible: propagation was significant in 185 of 189 analyzed recordings, the spatial firing-rate gradient in 174 and the firing-rate–latency correlation in 186 (**Fig. 3g–i**, **Supp. Fig. 5c**). The 14 mice with analyzable PSA events (out of 23 imaged) included all 10 more-affected mice but only 2 of 7 less-affected and 2 of 6 unclassified mice. Critically, these features were specific to synchronized events. In matched non-PSA windows from the same neurons, the spatial structure of firing rates was largely lost and no longer aligned with the propagation direction (**Fig. 3j–m**, **Supp. Fig. 5d–e**); the correlation between each neuron’s firing rate outside PSAs and its first-AP latency within PSAs fell to ρ = −0.166 ± 0.016, about three-fold weaker than within PSAs (split-half ρ = −0.517) (**Supp. Fig. 5b, f**); and spatial gradients fitted to one half of the windows failed to predict the other (split-half R² = 0.049 vs. 0.465 for PSAs; n = 172 recordings; **Supp. Fig. 5g–j**). Therefore, we conclude that PSAs are structured network events whose directional organization is specific to the synchronized state, rather than a fixed property of the recorded field.

### Data-driven quantification of epileptiform burden in repeated, continuous 24-hour EEG recordings

Because seizure quantification can depend on the seizure definition adopted^21^ and uses only a small fraction of each ∼24-h recording, we quantified EEG pathology as epileptiform burden across the entire recording. Discharges were detected as in **Fig. 1h** and converted into a continuous discharge density by using a 5-min sliding window. This signal was low and stable throughout vehicle recordings, whereas in pilocarpine-treated mice it showed large excursions lasting tens of minutes (**Fig. 4a**). We defined an epileptiform discharge episode (EDE) as an epoch of >5 min in which discharge density exceeded 8/min, spectral power exceeded recording-specific thresholds in five frequency bands (δ, θ, α, σ and β), and activity occurred concurrently in the hippocampal and cortical electrodes (**Fig. 4b**, Methods). EDEs thus require sustained epileptiform activity in both electrodes, but not ictal frequency or evolution. Review of simultaneous video for a subset of recordings showed that EDEs coincided with behavioral seizures, including generalized convulsions and behavioral arrest with myoclonic twitching (**Supp. Video 3**). Of 932 EDEs recorded after pilocarpine induction, 68 (7.3%) contained a segment meeting conventional electrographic seizure criteria (≥3 Hz for ≥10 s)^44^, as did none of the 14 EDEs detected in control recordings. On blinded review by an expert epileptologist, 48 of these 68 EDEs (71%) contained an unambiguous electrographic seizure, 4 were borderline and 16 were obscured by recording irregularities, such as movement-related transients (Methods). Confirmed seizures lasted 11–79 s (median 29 s; **Fig. 4a, inset ii**) and occurred in 11 of 19 pilocarpine-treated mice. Their share of EDEs rose with disease stage, from 4/264 acute-phase EDEs (1.5%; excluding the induction day; 4/18 mice) and 6/182 latent-phase EDEs (3.3%; 2/17 mice) to 35/399 chronic-phase EDEs (8.8%; 9/18 mice; denominators are mice with EDEs in each phase; chronic vs. earlier phases, p = 0.042, Wilcoxon signed-rank test on per-mouse proportions; **Fig. 4c**). EDEs therefore span the ictal–interictal continuum, with ictal content emerging as epilepsy becomes established.

**Figure 4.**
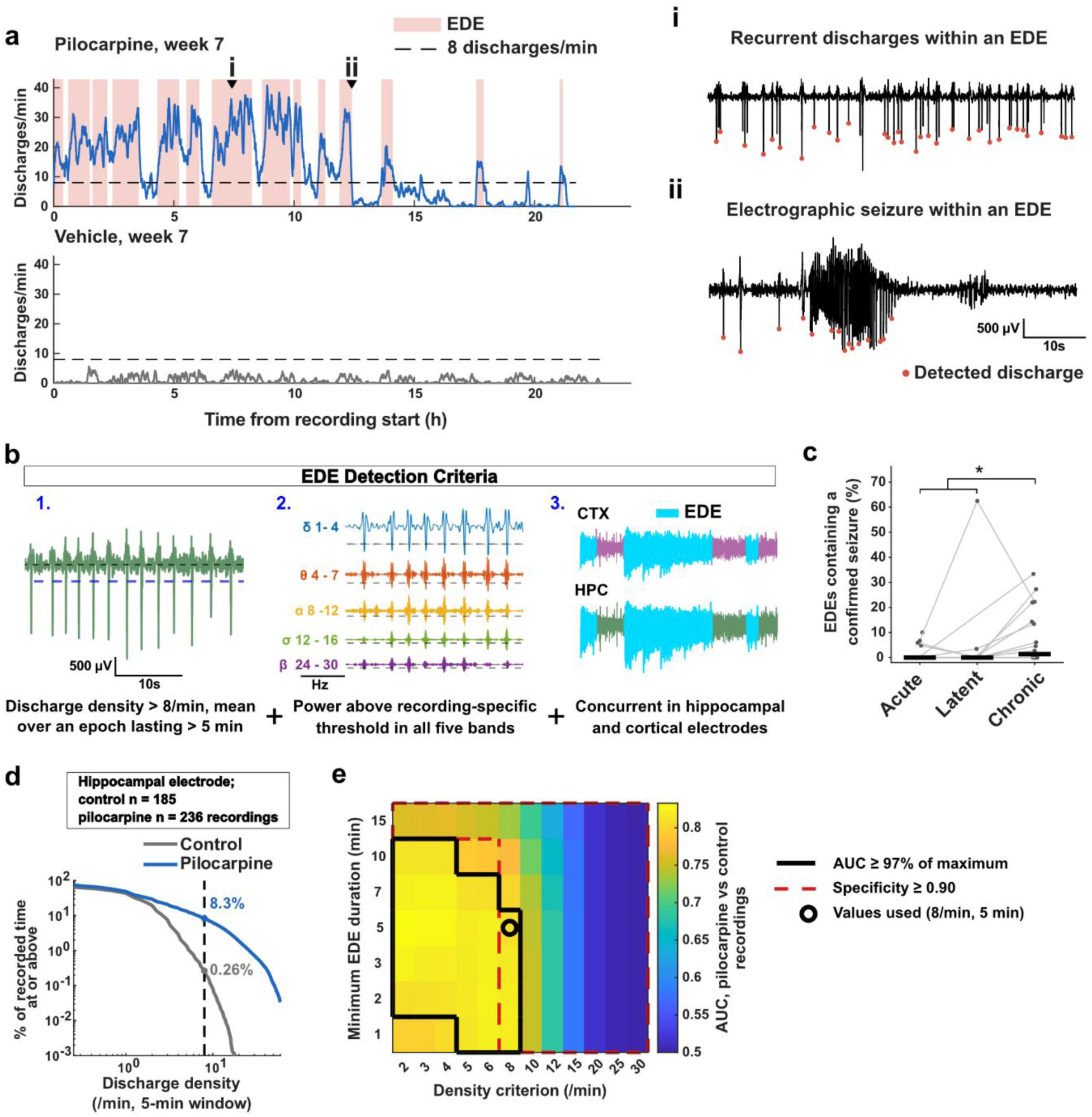
Data-driven quantification of epileptiform burden in repeated, continuous 24-hour EEG recordings. **(a)** Discharge density (hippocampal electrode, 5-min sliding window) across one ∼22-h recording from a pilocarpine-treated mouse (top) and a vehicle-treated mouse (bottom), both at week 7. Shaded epochs indicate EDEs. The dashed line marks the 8 discharges/min threshold, which applies to the mean density over an epoch, not to the instantaneous level. Triangles mark the 60-s windows shown in the insets: (i) recurrent discharges within an EDE; (ii) an electrographic seizure (∼15 s of rhythmic high-amplitude spiking) within an EDE. Red dots, detected discharges. Note that discharges during the seizure are not individually resolved (Methods). **(b)** EDE detection criteria. Discharges detected as in Fig. 1h are converted to a density signal. An epoch is an EDE if: its mean discharge density exceeds 8/min over >5 min; peak power exceeds recording-specific thresholds in all five frequency bands (δ: 1–4, θ: 4–7, α: 8–12, σ: 12–16, and β: 24–30 Hz); it overlaps in time with an epoch meeting criteria 1–2 in the other electrode. **(c)** Percentage of EDEs per mouse containing a confirmed electrographic seizure (a segment of ≥3 Hz for ≥10 s^44^ confirmed on blinded expert review), in the acute (excluding the induction day), latent and chronic phases. Each point is one mouse; lines connect data from the same mouse across phases; bars are medians (0%, 0% and 1.4%). Mice with no EDEs in a phase are not plotted (n = 18, 17 and 18 mice). *,p = 0.042, Wilcoxon signed-rank test, chronic vs. the mean of the acute and latent phases (bracket). **(d)** Percentage of recorded time at or above each discharge density, calculated for all control and pilocarpine-treated post-injection recordings (n = 185 and 236 hippocampal electrode recordings, respectively). The dashed line marks the selected threshold of 8 discharges/min, exceeded during 0.26% of control recording time and 8.3% of pilocarpine-treated recording time. **(e)** Area under the ROC curve for discriminating pilocarpine-treated from control recordings by EDE rate, across 84 combinations of density and duration criteria. Black outline: AUC within 3% of the maximum (0.833). Red dashed outline: specificity (proportion of control recordings without any EDE) ≥0.90. Circle: values used (8/min, 5 min; AUC 0.809, specificity 0.946, sensitivity 0.653).

The EDE density and duration criteria were set based on our own dataset (Methods), so we tested whether the conclusions depended on the values selected. Control recordings (n = 185: 22 vehicle baseline, 34 pilocarpine baseline, and 129 vehicle post-treatment) stayed below 8 discharges/min for 99.74% of recorded time, whereas pilocarpine-treated recordings (n = 236) spent 8.30% of their time at or above it, 32-fold more (**Fig. 4d**). We then recomputed EDE rate for every recording across 84 combinations of discharge density (2–30/min) and minimum-duration (1–15 min) criteria (**Fig. 4e**; Methods). Discrimination between pilocarpine-treated and control recordings by EDE rate, measured as the area under the receiver operating characteristic curve (AUC; 0.5, chance; 1, perfect separation), varied little across a broad region (AUC 0.80–0.83 across the 29 combinations within 3% of the maximum) that included the values used (AUC 0.809). A density of 8/min was the lowest at which specificity (the proportion of control recordings without any EDE) was ≥0.90 at every duration (0.946 at 5 min), whereas the maximal AUC, at 2/min, came at the cost of detecting EDEs in more than half of control recordings. The synchrony–burden correlation (acute PSA-AP vs. chronic EDE rate) was significant at every duration from 1 to 10 min for densities of 8–12/min (ρ = 0.49–0.68) and weaker at lower densities (ρ = 0.29–0.50; **Supp. Fig. 6a**). CA1 synchrony therefore correlates better with sustained high-density episodes than with lower-density ones. Because EDEs were counted as overlapping hippocampal–cortical event pairs, one long event in one electrode that overlapped two events in the other counted as two EDEs; merging such pairs into single episodes reduced the total from 946 to 808 EDEs without changing discrimination (AUC 0.809), prediction (AUC 0.931) or the synchrony–burden correlation (ρ = 0.655 vs. 0.620; Methods).

### Epileptiform burden persists through the latent period and correlates with CA1 synchrony

Weekly vEEG sessions, typically 23–24 h long (**Fig. 5a**, Methods), were analyzed with the acute, latent and chronic periods defined from the CA1 TPLSM data. EDEs emerged after pilocarpine, with epileptiform activity concurrent in hippocampal and cortical channels (**Fig. 5b**). Discharge rates in both electrodes rose sharply after induction, partially decayed within 1–2 weeks and remained above vehicle levels for the rest of the study (**Fig. 5c**, **Supp. Fig. 7a**); regional EDE duration, measured in each electrode without the concurrency requirement, followed the same pattern (**Supp. Fig. 7b**). We quantified epileptiform burden as the number of EDEs per hour (EDE rate) and their cumulative duration per recording. EDEs occurred in 51/68 acute, 30/50 latent and 73/118 chronic recordings from pilocarpine-treated mice (means of 0.27, 0.17 and 0.16 EDEs/h, respectively), but in only 10 of 185 control recordings (5.4%: 6/129 vehicle post-treatment, 2/22 vehicle baseline, 2/34 pilocarpine baseline), at a ∼50-fold lower rate (0.004 vs. 0.19 EDEs/h). Because approximately one-third of recordings from pilocarpine-treated mice (82 of 236) contained no EDEs, a threshold of one EDE per recording separated pilocarpine-treated from control recordings with an AUC of 0.809 (**Fig. 4e**). When each mouse’s EDE rate, averaged across post-induction recordings, was used to separate pilocarpine-treated from vehicle-treated mice, the groups showed near-complete separation (AUC 0.96; 20 vs. 13 mice, respectively; Methods). Sporadic epileptiform events are expected in control rodents^45, 46^; their rarity in control recordings indicates that EDEs reflect the post-pilocarpine condition rather than normal rodent EEG data or over-sensitivity of the analysis pipeline^47^. Epileptiform burden peaked in the first recording after induction, declined across the acute period, stayed lower but above vehicle levels during the latent period, and remained elevated, with some session-to-session variability, through the chronic period (**Fig. 5d-e, Supp. Fig. 7c–d**).

**Figure 5.**
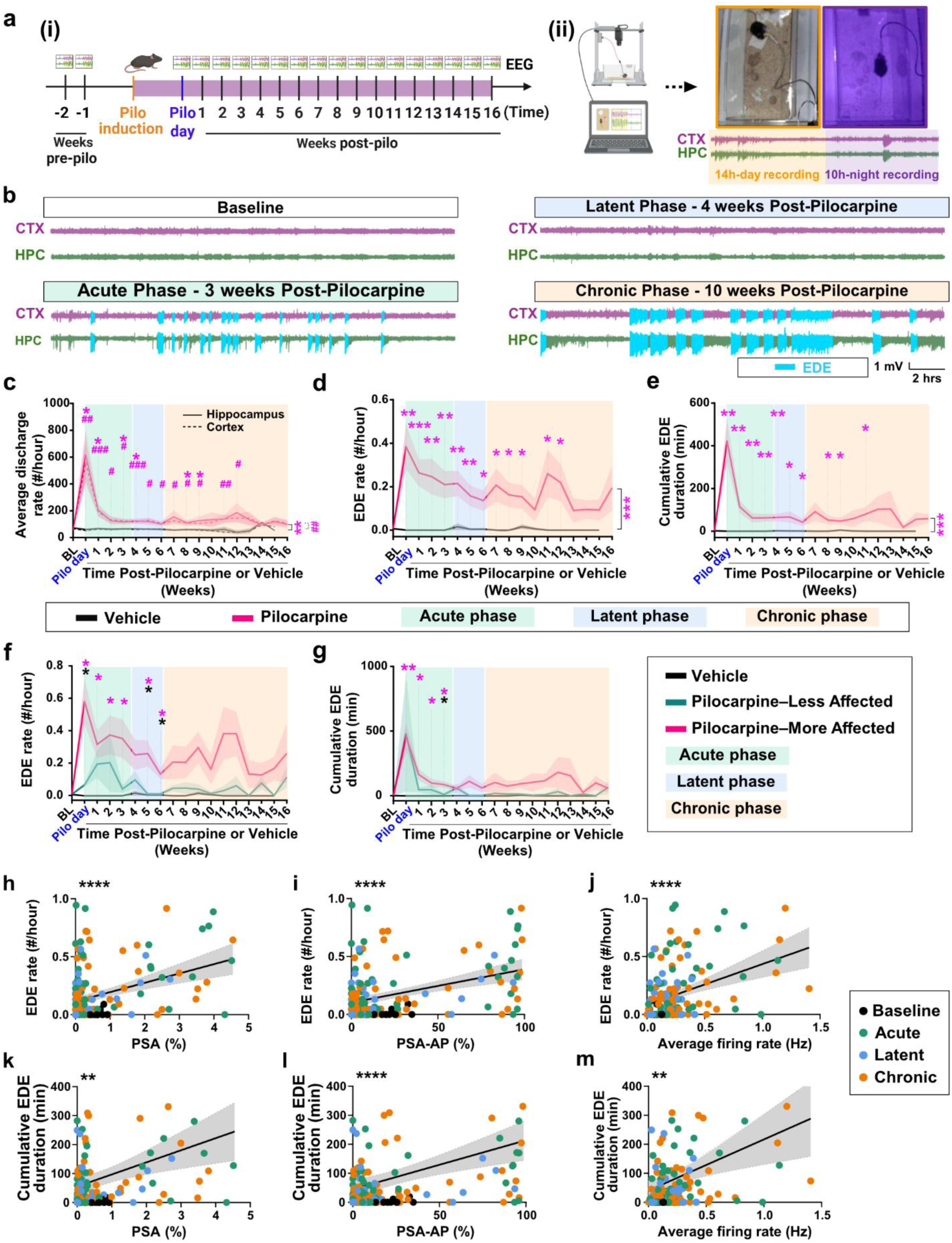
EEG monitoring reveals the effect of pilocarpine induction on epileptiform burden and its severity. **(a)** Experimental timeline of video-EEG recordings (i) and example images of daytime and nighttime monitoring (ii). **(b)** Example cortical and hippocampal (CTX and HPC, respectively) EEG traces from a pilocarpine-treated mouse during baseline, acute, latent, and chronic phases. Detected EDE times are overlaid in cyan on the corresponding traces. **(c)** Longitudinal changes in average discharge rate recorded over 24 h in hippocampal (solid line) and cortical (dashed line) electrodes. Pilocarpine-treated mice showed a marked acute increase after pilocarpine induction, followed by a prolonged decay and a subsequent chronic increase. **(d)** EDE rate over time in pilocarpine-treated vs. vehicle-treated mice. EDE rate rose sharply after induction and remained significantly elevated relative to controls throughout the recording period, with a gradual decline into the latent phase and intermittent increases and decreases during the chronic phase. **(e)** Cumulative EDE duration per 24 h recording, showing an acute increase similar to **(d)**, followed by a rapid decrease into the latent phase that remained significantly above control levels. During the chronic phase, EDE duration remained above control levels, with significant differences at several timepoints. **(f–g)** EDE rate **(f)** and cumulative EDE duration **(g)** in vehicle-treated mice and in the TPLSM-defined less-and more-affected groups (vehicle, n = 13; less-affected, n = 6; more-affected, n = 10; one less-affected mouse contributed imaging but not EEG data). Magenta, vehicle vs. more-affected; green, vehicle vs. less-affected; black, more-vs. less-affected. **(h-j)** Positive correlations between EEG-derived EDE rates and TPLSM-derived synchrony and average firing rate of CA1 neurons. **(k-m)** Positive correlations between EEG-derived cumulative EDE duration and TPLSM-derived synchrony and average firing rate of CA1 neurons. Data in **(c–e)** are presented as mean (solid lines) ± SEM (shaded areas). Shaded background regions denote disease phases: acute (day of pilocarpine to week 3, green), latent (weeks 4–6, blue), and chronic (weeks 7–16, orange). n=20 (11 males, 9 females) and 13 mice (7 males, 6 females) were included in the pilocarpine and vehicle groups at the study onset, respectively. Of these, 19 pilocarpine-treated mice contributed paired TPLSM and EEG recordings and are included in panels **(h–m**). See **Supp. Table 3** for mouse numbers in each subsequent recording week. Statistical analyses were performed using mixed-effects models with restricted maximum likelihood and Tukey’s HSD *post hoc* correction for multiple comparisons. Statistically significant changes in hippocampal and cortical recordings are indicated by asterisks (*) and hash (#) symbols, respectively. * or #,p<0.05; ** or ##,p<0.01; *** or ###,p<0.001; ****,p<0.0001. A complete list of significant comparisons is provided in **Supp. Table 3**. Each dot in panels (**h–m**) represents data from a single recording, color-coded by the disease phase at which the recordings were acquired (n=238 recordings from 19 pilocarpine-treated mice: 10 males, 9 females). Solid lines show the linear fit and shaded regions the 95% confidence interval. Additional information is reported in **Supp. Table 3**.

EEG measures segregated in parallel with the imaging-defined more- and less-affected groups (**Fig. 5f-g**). The more-affected group showed an acute rise in EDE rate, a latent-period decline and persistently elevated chronic rates that exceeded those of the less-affected group, which had a delayed acute peak and lower chronic rates (**Fig. 5f-g, Supp. Figs. 8–9**). Acute increases in cumulative EDE duration and discharge rates reached significance only in the more-affected group, owing to greater variability in the less-affected group (**Supp. Fig. 9**). EEG measures showed no consistent sex differences (**Supp. Fig. 10**).

Across recordings from the 19 mice with both EEG and TPLSM modalities, CA1 synchrony and firing correlated positively with EDE rate and cumulative EDE duration (**Fig. 5h-m, Supp. Fig. 11**), so cellular-resolution CA1 measures provide a quantitative readout associated with EEG-detected burden. We therefore asked whether early CA1 synchrony could predict the eventual severity of epilepsy in individual animals.

### Acute hippocampal synchrony and EDE rate predict chronic epileptiform burden

We analyzed all mice with acute (pilocarpine day–Week 3) and chronic (Weeks 7–16) measurements. Acute and chronic EDE rates were positively correlated (n = 19 mice; **Fig. 6a**), and PSA-AP levels were concordant between phases, with no mouse moving from a low acute to a high chronic state (n = 17 mice; **Fig. 6b**), indicating that early pathology persists into the chronic phase. To test individual-level prediction, we used logistic regression in the 17 mice with sufficient acute- and chronic-phase recordings (Methods), with mice classified as severe (n = 8) or non-severe (n = 9) from chronic EEG data alone (maximal chronic EDE rate >0.25/h; Methods), independently of the imaging-based grouping used above. Severe mice had more chronic-phase EDEs containing confirmed seizures than non-severe animals (29 vs. 6; present in 6/8 vs. 3/9 mice, respectively; p = 0.029, Mann–Whitney U test on per-mouse rates), but not a significantly higher proportion of them (29/311 vs. 6/52; p = 0.45, Mann–Whitney U test on per-mouse proportions): severe mice had more seizure-containing EDEs because they had more EDEs overall. Acute PSA-AP correlated with the chronic-phase rate of all EDEs (ρ = 0.62, p = 0.008; **Supp. Fig. 6a**) but not with the chronic-phase rate of the subset containing confirmed seizures (ρ = 0.13, p = 0.63; n = 17).

**Figure 6.**
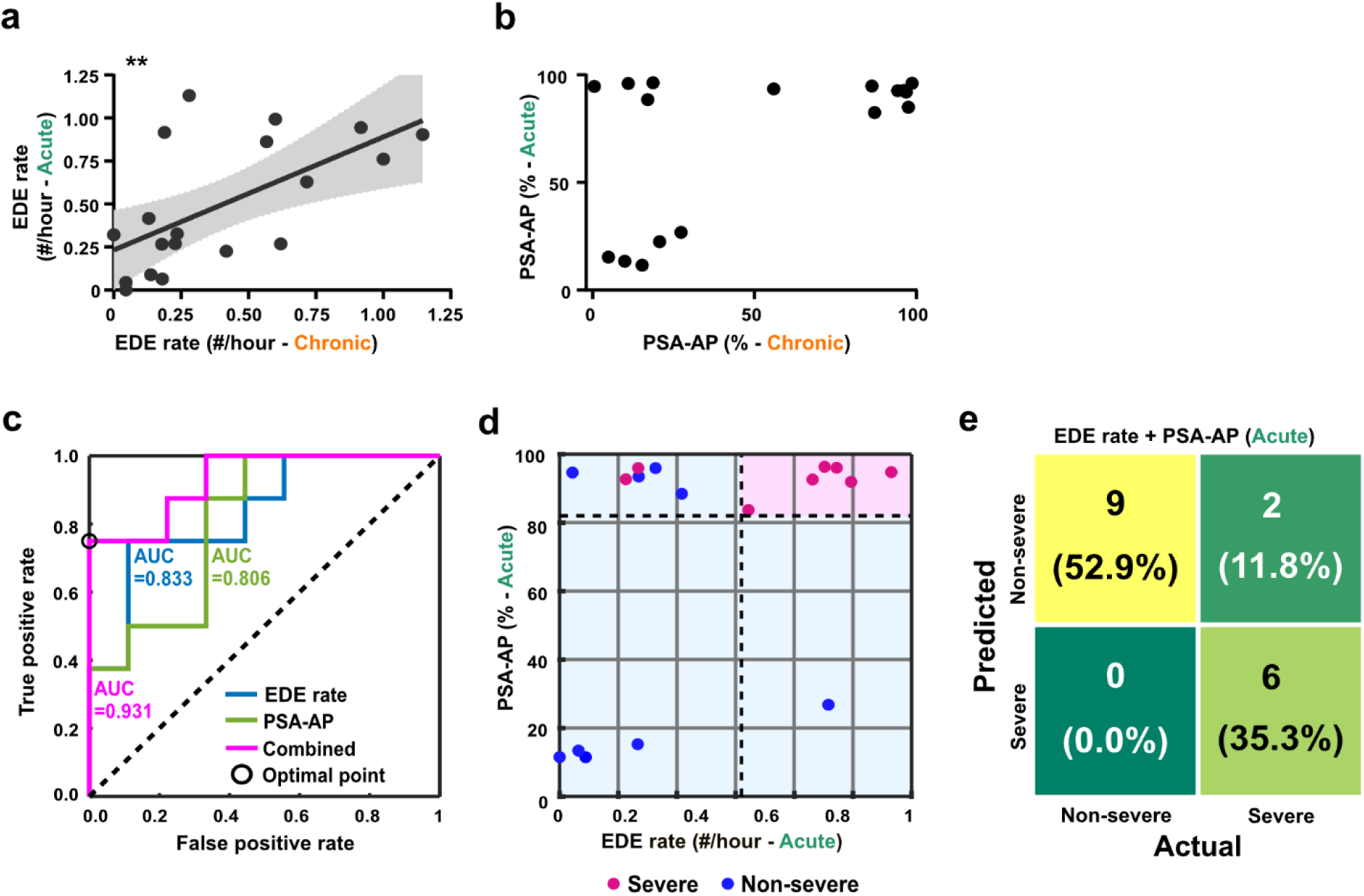
Acute changes in synchrony and EDE rate predict chronic epileptiform burden. **(a)** For pilocarpine-treated mice, EDE rates measured during the acute and chronic phases in the same animals show a positive correlation (p=0.0055; Pearson’s correlation). Each dot represents data from a single mouse, corresponding to the maximal value observed during the acute (pilocarpine day to week 3) and chronic (weeks 7–16) disease phases. A total of n=19 pilocarpine-treated mice were included (10 males, 9 females). Solid lines indicate linear regression fits, and shaded gray areas represent the 95% confidence intervals. **, p<0.01. **(b)** Same-mouse maximal PSA-AP values during the acute and chronic phases. PSA-AP values were bimodally distributed in both phases, separating into low (<50%) and high (>50%) groups. Taking 50% as a descriptive cut-off, 5 mice were low in both phases, 8 were high in both, 4 were high acutely but low chronically, and none were low acutely but high chronically. **(c)** Receiver operating characteristic (ROC) curves illustrating the performance of three logistic regression classifiers based on: (1) PSA-AP alone, (2) EDE rate alone, and (3) a combined model using both metrics. Each classifier uses acute-phase data to predict chronic severity (Methods). The dashed line marks the expected performance of a random classifier. **(d)** Visualization of the combined classifier from **c** using ROC-derived optimal decision thresholds set at the maximal Youden index for each parameter, as shown in **c**. The upper-right quadrant corresponds to predictions of severe chronic epileptiform burden. Data points from all participating mice (n=17) are plotted, with their true classifications color-coded. **(e)** Confusion matrix for the classifier in **d**, showing 6 true positive, 9 true negative, and 0 false positive, 2 false negative for the total of 17 mice in the dataset.

Maximal acute-phase PSA-AP predicted the same-mouse chronic severity (AUC = 0.806, p = 0.015; sensitivity 1.0, specificity 0.556, accuracy 0.765), as did maximal acute-phase EDE rate (AUC = 0.833, p = 0.0105; sensitivity 0.75, specificity 0.889, accuracy 0.824). A combined model including both predictors performed best (AUC = 0.931, p = 0.0079; sensitivity 0.75, specificity 1.0, accuracy 0.882; **Fig. 6c-e**). The combined model’s AUC remained 0.92–0.94 across the 29 criterion combinations of the discrimination plateau (**Fig. 4e, Supp. Fig. 6b**), and 0.79–0.93 across densities of 8–12/min and durations of 1–10 min when the severity outcome, itself defined from chronic EDE rate, was re-derived at each combination (**Supp. Fig. 6c**). The EDE-rate model drew much of its predictive power from days 0–1, whereas the PSA-AP model performed best on later acute-phase recordings (**Supp. Fig. 12a–b**). The combined model was well calibrated (Hosmer–Lemeshow p = 0.919), retained an AUC of 0.904 after bootstrap correction for optimism (the amount by which performance on the animals used for fitting may overstate performance on new animals), and remained the best model under leave-one-out cross-validation (AUC 0.833 vs. 0.750 for EDE rate and 0.611 for PSA-AP; **Supp. Fig. 12c–e**; Methods). Therefore, acute CA1 synchrony, measured within three weeks of induction, predicts chronic epileptiform burden, and combining it with EDE rate improves prediction over either modality alone.

## Discussion

We used a refined pilocarpine induction protocol that improved survival, allowing us to track the same CA1 neurons from before the epileptogenic insult through SE, the latent period and chronic SRS, alongside weekly hippocampal and cortical EEG recordings. We found that the trajectory of hippocampal network synchrony in the first weeks after SE predicted chronic epileptiform burden. Four findings support this conclusion: CA1 synchrony follows a three-phase trajectory that mirrors the clinical course of TLE; the synchronized events that define it, PSAs, propagate in a consistent direction and recruit neurons in a reproducible order; CA1 synchrony correlates with EEG-derived epileptiform burden in individual animals; and acute CA1 synchrony, combined with acute EDE rate, predicts chronic burden in individual mice, providing proof of concept for imaging biomarkers of epileptogenesis, a critical need in the field^17^.

Longitudinal single-animal studies require that animals survive induction, a persistent obstacle in the pilocarpine model, where mortality is substantial under standard regimens^36, 41, 42^ and induction success and survival vary with sex, strain and dose, so that many studies restrict analyses to a single sex^40, 44, 48^. Females are often reported to resist pilocarpine-induced SE and to die when it does occur^40, 42, 44, 48, 49, 50, 51^, whereas in our hands males were the more dose-sensitive sex. On the final protocol (sex-specific pilocarpine doses, with diazepam given according to Racine stage and SE duration), all mice reached SE and survived, induction severity was similar in males and females, and longitudinal imaging and EEG measures showed no consistent sex differences. The protocol therefore supports balanced male–female cohorts followed across the entire epileptogenic window, providing a reliable platform for testing candidate interventions.

The three-phase CA1 trajectory, elevated relative to vehicle in the acute and chronic phases but not in the latent phase, broadly aligns with the canonical model of epileptogenesis^6, 7, 9^. The EEG analysis adds a qualification: few latent-period EDEs contained seizures, consistent with the classically seizure-free latent period^36^, yet epileptiform burden remained above vehicle levels throughout. This supports the view that epileptogenesis is a continuous process in which network reorganization proceeds without interruption^6, 7, 9^, and implies that a window for antiepileptogenic intervention, if it exists, likely extends beyond the acute phase^17^. Conversely, because the latent period can be short in the mouse pilocarpine model^52^, the few seizure-containing EDEs recorded within our operationally defined acute period after the induction day may reflect an early onset of spontaneous seizures in individual mice. Unlike epileptiform burden, CA1 synchrony returned to vehicle levels during the latent period; whether this reflects transient stabilization of the CA1 network or ongoing changes that somatic calcium imaging of CA1 excitatory neurons does not capture remains to be determined.

PSAs resemble the interictal population-wide synchronizations described by Muldoon et al. in chronically epileptic pilocarpine-treated mice^53^ in their brevity and extent, but differ in composition. Those events recruited only 0.6% of stratum pyramidale cells, versus 33% of stratum oriens cells, and were followed by suppressed pyramidal-layer firing, indicating preferential recruitment of GABAergic interneurons^53^, whereas PSAs consist of synchronized firing of many excitatory neurons. This may reflect distinct event classes, how events were identified (EEG-triggered vs. optically defined, the latter selecting for high recruitment), or indicator expression (virally expressed GCaMP5G or GCaMP6m under the human synapsin promoter, which labels both excitatory and inhibitory neurons, vs. transgenic Thy1-jRGECO1a expression, which is predominantly restricted to excitatory neurons). In another study, Sparks et al. imaged dentate granule cells during interictal discharges after intrahippocampal kainate, through a window implanted three days after the insult that required partial aspiration of dorsal CA1^54^. They found that interictal discharges recruit distinct granule-cell ensembles dominated by adult-born neurons. In these studies and in miniscope recordings from CA1 of pilocarpine-treated mice, which showed place-cell instability and interneuron desynchronization emerging ∼6 weeks after SE, imaging began after the insult and covered only the chronic phase^53, 54, 55^. Our window implantation removed no hippocampal tissue and was performed ∼6 weeks before pilocarpine induction, so the same neurons could be imaged at baseline and from the day of induction onward, showing that PSAs emerge acutely and persist into the chronic phase.

Two PSA properties stand out. First, PSAs propagated as posterior-to-anterior waves, linking microcircuit-scale interictal synchrony to the traveling waves described in human cortex, where interictal discharges travel along the same paths as ictal discharges, mostly in one predominant direction^56^. Because posterior dorsal CA1 lies toward the temporal pole, this direction is compatible with recruitment from the ventral hippocampal formation, the principal seizure-onset zone in pilocarpine-treated rats^57^. Second, recruitment within a PSA was graded by firing rate: the most active neurons fired first and most consistently. Both the direction of travel and the recruitment order were reproducible across independent halves of PSA events but absent or much weaker outside PSAs, so they are properties of the synchronized state rather than of a fixed set of intrinsically hyperactive cells. This structure was detected predominantly in more-affected mice.

We specifically quantified overall epileptiform burden rather than evaluating only seizure metrics. EDEs are prolonged epochs of dense epileptiform discharge, concurrent in hippocampal and cortical electrodes, that include electroclinical seizures but do not require ictal rhythm or evolution, so they span the ictal–interictal continuum, with ictal content increasing as epilepsy became established. Measuring this mixture is informative: activity short of seizure criteria need not be benign, as regional hypermetabolism on FDG-PET identified critically ill patients whose ictal–interictal continuum patterns represented SE^58^, and continuous EEG measures index seizure propensity in mice even when seizures are rare^20^. Because seizure phenotypes and drug responses vary with mouse strain and are sensitive to the seizure definition adopted^21, 59^, and baseline EEG varies across strains^45, 46^, our threshold values are unlikely to transfer directly to other studies, but the procedure used to derive and test them could be adopted in other experimental epilepsy models.

CA1 synchrony tracked epileptiform burden in individual animals, and the imaging-based grouping, made without any input from EEG recordings, segregated EEG-derived burden. Synchrony within a ∼600 µm x 600 µm field of excitatory neurons therefore carries quantitative information associated with network-wide activity in the contralateral hemisphere, consistent with the central role of the hippocampus in this model^36^ and in human mesial TLE^28^. The electrographic counterpart of PSAs can only be inferred. Their ∼100-ms duration and sub-Hz recurrence match interictal epileptiform discharges^24^ rather than seizures, yet their strong recruitment of excitatory neurons contrasts with the sparse pyramidal-cell recruitment during EEG-detected interictal spikes^53^. PSAs may therefore correspond to a subset of interictal events, or to local population bursts too confined to register on a field electrode^60^. Either way, their association with chronic burden parallels the link between preceding interictal spike frequency and the rate of epileptogenesis after kainate-induced SE^26^. The two modalities were also complementary in prediction: the EDE-rate model drew most of its information from the first days after induction, whereas the PSA-AP model relied on later acute-phase sessions.

Prognostication after SE relies on macroscale clinical and EEG scores that estimate related but distinct outcomes, such as the risk of epilepsy after de novo SE^61^, and individual-level prediction remains difficult^17^. Interictal activity predicts seizure burden in a genetic epilepsy model but was measured at mesoscale resolution within a single disease state^27^, and early EEG measures have anticipated later epilepsy in several rodent models, including after traumatic brain injury^62, 63^. Our results extend this work in three ways: the predictor is a cellular-resolution measure of hippocampal synchrony rather than a field-potential feature; imaging and EEG from the same animals carried complementary information, so the combined model outperformed either alone; and synchrony imaged in one hemisphere forecast burden recorded from the other, indicating that the local cellular signal reflects a global disease state. The first three weeks after SE are therefore a candidate interval for early stratification and intervention.

The divergence between more- and less-affected mice within the first week, well before chronic seizures, constrains possible mechanisms. Structural remodeling after SE, such as aberrant integration of adult-born granule cells and mossy fiber sprouting, takes weeks to develop^14, 64, 65^ and is unlikely to explain synchrony changes that emerge within days. These may instead reflect an early shift in excitation–inhibition balance, consistent with reduced GABAergic inhibition early after pilocarpine-induced SE^11, 12, 13^, with slower structural reorganization later stabilizing the chronic state. Whether manipulating dentate neurogenesis^44, 49^, or applying anti-inflammatory or neuroprotective treatments, alters the PSA trajectory remains to be tested.

Several limitations of the study should be noted. The pilocarpine model reproduces core features of human TLE but not its etiological diversity, and replication across models and species is needed^36, 38^. TPLSM and EEG were acquired in separate sessions from opposite hemispheres, so testing whether individual PSAs coincide with specific electrographic events, such as ipsilateral interictal spikes, will require concurrent recording. Sampling was intermittent: weekly ∼24-h EEG sessions and ∼200-s TPLSM recordings per FOV capture a small fraction of each week, and because spontaneous seizures cluster, with seizure-free intervals of days to weeks^66, 67^, individual sessions can fall within or between clusters. EDE detection merges activity separated by less than 5 min and cannot resolve discharge rates above ∼2 Hz, and 200-Hz sampling precluded analysis of high-frequency oscillations, a candidate EEG biomarker of epileptogenesis^68^; ictal-frequency segments were therefore identified separately and confirmed on blinded expert review, and how many EDEs carry a behavioral correlate will require systematic review of time-aligned video^45, 69^. Because jRGECO1a is expressed predominantly in excitatory neurons^39^, the contribution of GABAergic interneurons to PSAs, likely substantial given their role in interictal dynamics^53^ and their vulnerability during epileptogenesis^11, 12, 13^, remains uncharacterized. The window preparation leaves the hippocampus intact but removes the overlying cortex and corpus callosum; although this was done ∼6 weeks before induction and identically in vehicle-treated mice, some influence on epileptogenesis cannot be excluded. Diazepam was given once mice reached the behavioral SE criterion, which shortens SE, and shorter SE may lead to fewer SRS^70, 71^. EDEs containing confirmed seizures were correspondingly infrequent, making seizure counts a low-yield endpoint in this cohort^20^: acute CA1 synchrony correlated with overall epileptiform burden but not with the rate of seizure-containing EDEs. Testing whether synchrony predicts seizure frequency will require longer continuous monitoring, larger cohorts or longer SE, the last at the cost of higher mortality^36, 41, 42^. Finally, the predictive analysis comprised 17 mice; despite penalized regression and bootstrap correction, prospective testing in larger, independent cohorts is required before PSA-based metrics can be considered biomarkers^17^.

These limitations notwithstanding, our findings demonstrate that the degree to which the CA1 network is disrupted during the first three weeks after an epileptogenic insult predicts chronic epileptiform burden. This dysfunction is readable at cellular resolution, detectable before chronic seizures emerge, and adds to conventional EEG monitoring when the two are combined. Identifying who will develop severe epilepsy, and intervening before chronic seizures take hold, remains a defining unmet need in epilepsy care; our data suggest that the relevant signal is already present in the hippocampus within weeks of the initial injury.

## Methods

### Animals and surgery

All surgical and experimental procedures were conducted in accordance with protocols approved by Cleveland Clinic Research Institutional Animal Care and Use and Institutional Biosafety Committees. Mice of the Thy1-jRGECO1a line 8.31^39^ were bred on a C57BL/6J background and crossed with C57BL/6 mice from Charles River Laboratories. Heterozygous jRGECO1a-positive offspring (n=54 mice, 33 males and 21 females), aged 12–20 weeks at the time of surgery, were used. Mouse numbers, group assignments, and inclusion criteria for each analysis are summarized in **Supp. Table 1**. Mice were housed in a temperature-controlled environment with a 14 h light/10 h dark cycle and *ad libitum* access to food and water. All mice underwent stereotaxic surgery for hippocampal window implantation, as previously described^72, 73^, together with EEG electrode implantation. Mice were anesthetized with isoflurane (2.5–3% induction, 1.5% maintenance) and placed on a 37°C heating pad. Local anesthesia (0.5% bupivacaine, s.c.) was administered at the craniotomy site. After scalp shaving and cleansing with Betadine (Purdue Products), lubricating gel was applied to protect the eyes from dehydration (Patterson Veterinary). A circular craniotomy was drilled above the right hippocampus (3.2 mm diameter; center at -2 mm AP, +2 mm ML relative to Bregma; Omnidrill35, World Precision Instruments) to expose the brain. The cortex and corpus callosum were carefully removed by gentle suction, avoiding damage to the alveus overlying the dorsal hippocampus. A glass coverslip (3 mm diameter, #1 glass, Warner Instruments) glued to a stainless-steel cannula (internal diameter 2.65 mm, outer diameter 3.2 mm, length 1.8 mm; New England Small Tube Corp.) was placed over the exposed hippocampus and secured with dental cement (Contemporary Ortho-Jet, Lang Dental). A custom stainless-steel head post was attached to the skull using the same cement for head fixation during imaging.

For EEG electrode implantation, three small craniotomies were drilled using the same system at stereotaxic coordinates over the somatosensory cortex (−0.5 mm AP, −3.5 mm ML), the left hippocampus (−2.1 mm AP, −4 mm ML), and the olfactory bulb, which served as the reference site. Subdural screw electrodes (0.20″ length, stainless steel jeweler’s screws, #E363/20/1.6/SPC, Protech International) were placed over the somatosensory cortex and olfactory bulb within their respective craniotomies. Electrodes were positioned primarily within the skull and subdural space, although the tips could contact the brain surface or superficial cortex. For hippocampal recordings, an intra-hippocampal electrode (0.18″ length, stainless steel jeweler’s screw, #E363/2/SPC, Protech International) was implanted through the hippocampal craniotomy at a 30° angle relative to the AP axis. All electrodes were prefabricated with gold pins to enable rapid connection to the preamplifier during recording sessions. The electrode over the somatosensory cortex is referred to as the cortical electrode, and the electrode over the hippocampus is referred to as the hippocampal electrode.

Postoperatively, animals received extended-release buprenorphine for analgesia, meloxicam for anti-inflammatory treatment, and TMS antibiotic in the drinking water for three weeks to prevent infection. Mice were transferred to a clean cage after surgery and maintained on a 37°C heating pad until fully ambulatory. All mice were allowed a minimal recovery period of 33 days to permit glial responses to subside, hippocampal windows to become optically clear, and neuronal activity to stabilize.

### Perfusion and tissue processing for histology

Mice were initially anesthetized with ketamine and xylazine (100–200 and 15–30 mg/kg BW, respectively). Following confirmation of deep anesthesia, an abdominal incision was made, and transcardial perfusion was conducted using 10 mL of cold saline, followed by 20 mL of cold 4% paraformaldehyde in 0.1M Dulbecco’s phosphate buffer saline (DPBS; pH 7.4) via a peristaltic pump. The brains were removed from the skull and stored overnight at 4°C in 4% paraformaldehyde in 0.1 M DPBS. Subsequently, brains were dehydrated in sucrose for 24 hours (or until they sank), embedded in OCT, and preserved at -80°C. Coronal sections (30μm thick) were then prepared using a cryostat (Leica Biosystems CM 1950) and stored in 24-well tissue culture plates with cryoprotectant solution (25% glycerol, 30% ethylene glycol, 45% 0.1 M PB, pH 6.7) at 4°C.

Tissue sections were incubated overnight at 4°C with primary antibodies, including rabbit polyclonal anti-Iba-1 (1:200; Biocare Medical #CP2090B), mouse anti-NeuN (MAB377, Millipore, 1:500), and rat anti-GFAP (1:500; ThermoFisher #13-0300). Following primary antibody incubation, sections were incubated with secondary antibodies for 1 hour at room temperature in the dark, rotating at 15 rpm. Secondary antibodies included FITC-conjugated donkey anti-rabbit (1:1000; Jackson Immunoresearch #711-095-152), Cy3-conjugated anti-mouse (715-165-150, Jackson ImmunoResearch, 1:1000), and Cy5-conjugated donkey anti-rat (1:500; Jackson Immunoresearch #712-175-150). Sections were mounted onto slides, counterstained with DAPI, and mounted with DAPI medium (Aqueous Fluoroshield, Abcam). Images were acquired using a 4× 0.2 NA or a 20× 0.75 NA objective lens using a BZ-X710 epifluorescence microscope (Keyence) and the BZ-X700 Viewer software.

### Pilocarpine-induced SE

SE was induced in cohorts of 2–4 mice using a modified pilocarpine protocol adapted from Botterill et al. and others^40, 44, 49^ to model TLE. Mice first received a subcutaneous injection of the muscarinic antagonist scopolamine methyl nitrate (1 mg/kg, ChemCruz #SC-264301) to minimize peripheral cholinergic effects, the β2-adrenergic agonist terbutaline hemisulfate (1 mg/kg, Sigma #T2528) to support respiration, and ethosuximide (150 mg/kg, Sigma #E7138) to reduce brainstem seizure-related mortality^40, 74^. Approximately 22-25 minutes later, pilocarpine hydrochloride (Sigma #P6503) was administered subcutaneously at 180 mg/kg for males and 250 mg/kg for females to induce SE (refined protocol that was used for all mice that went into the study). A small cohort of male mice received 215 mg/kg to compare survival as shown in **Fig. 1b**.

Seizure severity was scored using a modified Racine scale^29^, with scores recorded every 5–10 minutes after pilocarpine injection. SE induction was defined behaviorally as reaching stage 4 (males) or stage 5 (females); by design, this threshold was also the trigger for administering diazepam (10 mg/kg, i.p.) to limit mortality^40, 75^. Mice that had not reached this stage within 3 hours of pilocarpine were to receive diazepam at that timepoint, but in practice every mouse reached the threshold beforehand. Cumulative seizure severity was calculated as the total score prior to diazepam administration. If mice continued to exhibit seizures of stage 2 or higher 45 minutes after diazepam or 5 hours after pilocarpine, a second diazepam dose (5 mg/kg, i.p.) was given. Fourteen of the 23 mice in the two final-dose 2x diazepam arms (180 mg/kg males and 250 mg/kg females; **Fig. 1b**) received a second dose: 13 met this criterion and received it 50 min to 5 h after the first dose, and one received it 8 min after the first dose because of breathing difficulty.

At the end of the SE-induction protocol, mice received 1 mL sterile ACSF subcutaneously for hydration. When the post-SE clinical condition permitted, mice were immediately placed under the TPLSM microscope for CA1 imaging, followed by an EEG recording session; if the mouse condition was poor, the first TPLSM recording was postponed to the following day.

### TPLSM imaging

Activity monitoring of CA1 pyramidal neurons in the hippocampus began at least 33 days post-craniotomy to ensure adequate recovery. TPLSM was performed using a custom-built two-photon microscope (MIMMS 2.0, Janelia Research Campus design, https://www.janelia.org/open-science/mimms-22-2024) equipped with resonant–galvo–galvo scanners, a 16× water-immersion objective (N16XLWD-PF, Nikon, NA = 0.8), and an excitation wavelength of 1100 nm (InSight X3, Spectra-Physics). Emitted fluorescence was collected through dichroic beamsplitters and emission filters (705 nm and 561 nm dichroic filters, 625/90 nm BrightLine emission filter; all Semrock) and detected with a GaAsP photomultiplier tube (PMT2100, Thorlabs). Movies of each FOV consisted of 200 s recordings at 30 frames per second with 512 × 512 pixels, covering approximately 600 × 600 µm².

Prior to each recording, awake mice were head-fixed on a rodent belt treadmill (HHMI Janelia Research Campus design, DOI: https://doi.org/10.25378/janelia.24691311). Animals were habituated to the treadmill for five days before the first imaging session to reduce stress. Each recording session captured spontaneous activity from 5–7 FOVs within CA1, typically 100-250 μm under the glass window. Excitation power was adjusted between 50–200 mW using a Pockels cell (Conoptics) or fast laser power modulator (OM6ENH, Thorlabs) to maintain consistent signal-to-noise ratios across animals. Data acquisition was synchronized and controlled with ScanImage software (MBF Bioscience).

### TPLSM data analysis

For data analysis, beginning one week after pilocarpine induction, recordings from all mice were grouped into weekly bins defined as ±2 days around the nominal recording day; accordingly, week 1 comprised days 6-10 post-induction (centered on day 8), week 2 comprised days 13-17 (centered on day 15), and so on for subsequent weeks.

Imaging data were analyzed using custom MATLAB (MathWorks) scripts. Motion artifacts and slow drifts over the 200 s recording period were corrected with the TurboReg^76^ plug-in for ImageJ or MATLAB’s imregister function. Frames with residual gross motion, such as vertical brain displacement, were excluded. Regions of interest (ROIs) corresponding to individual somata were segmented with Cellpose2^77^ and manually inspected and corrected. For each ROI, fluorescence intensity (F) was computed as the mean pixel value within each ROI, and baseline fluorescence (F_0_) was defined as the median fluorescence across the recording. No neuropil correction was applied because CA1 pyramidal somata are densely packed with little intervening neuropil; consistent with a negligible neuropil contribution, the recruitment order within PSAs was reproducible across events and absent outside PSAs (**Supp. Fig. 5**). Relative fluorescence changes (ΔF/F_0_) were then calculated, and AP firing was estimated using the MLSpike model^43^. MLSpike model parameters for jRGECO1a followed published values^33^ (AP amplitude 15%, decay time 175 ms, Hill coefficient 1.9, saturation 0.001, drift correction 0.03). Model outputs were visually inspected for accuracy. Average firing rate was quantified as the total AP count divided by the recording duration.

### PSA data analysis

#### Detection and metrics

The onset (t=0) of a PSA event was defined as the first frame in which the number of detected APs across all recorded neurons was at least 25% of the number of neurons in the FOV. To reduce the dependence of PSA detection on the single-frame accuracy of the MLSpike model, we summed APs over the two frames preceding and following the MLSpike detection frame and required independent PSA events to be separated by at least five frames. Analysis of PSA event properties was performed for time windows spanning 1 frame prior to PSA onset to 3 frames after detection (5 frames in total: −1, 0, +1, +2, +3). This analysis window is distinct from the symmetric ±2-frame window used above for event detection. Next, we counted the number of frames falling within PSA time windows and, to account for differences in the number of recorded frames across FOVs and recording sessions (e.g., due to removal of motion-contaminated frames), we normalized this count by the total number of frames to obtain the percentage of PSA-positive frames (PSA%), which we use as a measure of the proportion of the recording occupied by synchronized activity. For each FOV, we also calculated the proportion of APs occurring during PSA events by dividing the total number of APs within all PSAs (the number of APs that were detected in the frames falling within any PSA time window, as defined above) by the total number of APs in the recording (PSA-AP). To quantify disease severity in each recorded mouse, we used the maximal value for mean firing rate, PSA% and PSA-AP across all recorded FOVs as the representative value for subsequent analyses (see **Fig. 2** and **Supp. Fig. 3-4**).

#### PSA spatiotemporal analyses

For PSA spatiotemporal analyses, we included individual recording sessions containing at least 10 distinct PSAs, each separated by at least 10 frames, a stricter separation than the 5-frame minimum used for event detection. To detect temporal propagation and spatial distribution of firing rates during PSA events (**see Fig. 3c-d**), we identified timing of the first AP fired by each cell within each PSA time window; APs outside this window were excluded. PSA events in which >80% of neurons fired in the same frame were excluded as well, due to insufficient temporal resolution for analysis. For the included PSAs, spatiotemporal planes were fit by regressing each cell’s latency from the t=0 frame to its first AP against that cell’s ROI-center coordinates: Latency(*x*, *y*) = *β*_1_*x* + *β*_2_*y* + *β*_3_, where (x,y) are the pixels in the recorded FOV. Propagation speed was computed as the inverse of the gradient magnitude, scaled by pixel size and frame rate. Propagation direction was defined as the angle of the fitted gradient vector and expressed in degrees. Candidate propagation events were further filtered based on fit quality and physical constraints: FOVs with fewer than 20 active cells, gradient magnitude < 0.001frames/pixel, *R*^2^ < 0.01, or speeds > 2.61 cm/s (maximum speed to traverse the FOV diagonal within a single frame acquisition time) were excluded. To assess the significance of the population-level propagation, we used a permutation test, where we compared the observed gradient to a null distribution generated from 2,000 permutations of the measured cell latencies; recordings were considered to exhibit significant propagation if the magnitude of the observed gradient exceeded the 95th percentile of the shuffled distribution (p<0.05).

For spatial distribution of firing rate gradient analyses, two-dimensional spatial planes were fit by relating each cell’s mean firing rate across either the entire recording, the PSA events, or the entire recording with the PSA events removed, to their ROI center coordinates: Firing Rate*(x,y) = β₁x + β₂y + β₃*. The opposite direction to fitted gradient vector (β₁, β₂) defined the steepest-descent direction of decreasing activity across the FOV, expressed in degrees, and its magnitude provided a measure of the spatial firing rate gradient, and expressed in Hz/mm (**Fig. 3j**). The goodness-of-fit of the spatial model was assessed by R², and significance of the population-level gradient was evaluated using an F-test of the multiple regression model. Only recordings that exhibited a significant spatial firing rate gradient based on the F-test (p<0.01) and permutation test (p<0.001; 2,000 label-shuffling iterations) were included in subsequent analyses.

To evaluate the relationship between neuronal mean firing rates and timing relative to onset of PSA events, we used the same first-AP latency data we calculated above. For each neuron, mean and standard deviation of first-AP latencies across either the entire recording, the PSA events, or the entire recording with the PSA events removed, were calculated. Monotonic relationships between firing rate and average latency were tested using Spearman’s rank correlation. To visualize variability, we generated scatter plots of firing rate vs. average latency, color-coded by the standard deviation of each cell’s latency.

#### Split-half cross-validation and matched non-PSA controls

To test whether PSA propagation reflected reproducible spatial structure rather than within-event contamination, PSA events in each recording were randomly divided into two non-overlapping halves. A spatial latency gradient was fitted to one half (as described above), and the resulting coefficients were applied to the neuron coordinates of the second (held-out) half to generate predicted latencies; split-half R² was computed between predicted and observed latencies in the held-out half and averaged across 200 random splits per recording. R² is reported as the squared Spearman rank correlation between predicted and observed held-out latencies and therefore quantifies the reproducibility of recruitment order on a 0–1 scale rather than absolute predictive error. The same procedure was applied to matched non-PSA control windows, constructed by drawing an equal number of frames with random onset times. An equivalent split-half approach was used for the firing-rate–latency relationship, with firing rate computed from one random half of the PSA events and first-AP latency from the other half (100 random splits per recording). Spatial analyses used the entire dataset of 174 recordings with a significant spatial firing-rate gradient (F-test, p<0.01); latency analyses used the entire dataset of 189 recordings containing at least 10 distinct PSA events; the propagation split-half analysis used 172 of these 174 recordings, excluding two in which non-PSA windows contained too few active neurons to yield 200 valid permutations; the firing-rate–latency split-half analysis used all 189 recordings.

#### Severity grouping

Mice were classified into less- and more-affected groups using two thresholds applied to two different TPLSM-derived metrics. The first threshold applied to PSA% and was individualized for each mouse: we calculated the mean PSA% across all recorded FOVs of that mouse’s two baseline recording sessions, and set the threshold at three times this baseline value. The second threshold applied to PSA-AP and was fixed at 80% for all mice (**Supp. Fig. 4a**). A mouse was classified as more-affected if, across all post-pilocarpine recordings, its PSA% exceeded the first threshold in at least two recording sessions and its PSA-AP exceeded the second threshold in at least one recording session. All other mice were classified as less-affected. Thresholds were set from the data: PSA% was normalized to each mouse’s own baseline to remove differences in baseline synchrony between animals, and had to be exceeded in at least two sessions to distinguish recurrent from single-session elevations; the PSA-AP cut-off of 80% lies in the gap between the low and high modes of its bimodal distribution (**Fig. 6b**). Six mice were not classified in either severity group because they lacked sufficient chronic phase recording data.

### Video-EEG recording and analysis

#### Acquisition

EEG recording sessions were conducted weekly for approximately 23-24 h (in most cases, barring animals with shorter recordings due to technical issues), with simultaneous video monitoring. Two baseline sessions were obtained before pilocarpine induction, after which weekly recordings were collected throughout the 16-week study period. Similar to the TPLSM analysis, post-induction recordings were grouped into weekly bins defined as ±2 days around the nominal recording day. During recordings, mice were placed in one of four custom-built cells positioned inside their home cages with the cage lids removed. The three EEG electrodes were connected to a custom-made connector (Protech International, Inc.) and then to an EEG recording system (Nihon Kohden, Neurofax Amplifier, #JE-921A) that was modified to accommodate the lower channel count required for mouse recordings. Signals were routed through a pendulous electroslip ring, which allowed free movement while minimizing motion artifacts, and were sampled at 200 Hz. Mice had *ad libitum* access to food and water throughout each recording session. On pilocarpine induction day, when mouse condition allowed, TPLSM imaging occurred after the end of the pilocarpine protocol, followed by EEG recording that began the same evening; thereafter, TPLSM and EEG sessions were acquired on separate days within the same weekly bin.

#### Discharge detection and EDE definition

Clinical criteria for electrographic seizures require epileptiform discharges averaging >2.5 Hz for ≥10 s or a pattern with definite evolution^24^. Electrographic seizures in rodents are conventionally scored by visual inspection as rhythmic discharges of ≥3 Hz lasting ≥3–10 s^40, 44, 78^. Notably, what should count as a seizure in rodent recordings remains debated^79, 80^, and quantitative outcomes in rodent models can depend on the seizure definition adopted^21^. Visual scoring is, moreover, not tractable for ∼24-h recordings from every mouse across 16 weeks. We therefore defined an automated burden measure, implemented in custom MATLAB scripts (steps 1–6 below), that first identified individual epileptiform discharges by amplitude-thresholding (adapted from Iyengar et al.^74^ and Botterill et al.^40^), then integrated discharge density and multi-band spectral power to identify prolonged epochs of epileptiform activity occurring concurrently in hippocampal and cortical electrodes, which we term epileptiform discharge episodes (EDEs). Two features of this definition should be noted. First, discharges are counted at most once per ∼0.5 s (step 2), so the detector cannot resolve discharge rates above ∼2 Hz and cannot establish whether ictal-frequency segments occur within an EDE. Second, EDE boundaries derive from a 5-min integration window (step 4), so EDE duration measures the period of elevated discharge density rather than the duration of continuous ictal activity. Following preclinical common data element terminology^69^, EDEs are therefore not scored as electrographic or electroclinical seizures.

Raw EEG signals were first processed with a digital multi-bandstop filter to remove 60 Hz line noise, its harmonics, high-frequency noise, and low-frequency drift (zero-phase FIR filter; passband ∼1–75 Hz, with stopbands at 16–24, 36–44 and 56–64 Hz). The first and last 30 min of each recording were excluded to avoid periods when experimenter proximity might affect the mice. EEG analysis proceeded as follows:

1. For each electrode, we computed a threshold T equal to the signal root mean square (RMS) + 3 times its standard deviation over the entire recording and detected all samples crossing below −T (i.e., negative deflections whose amplitude exceeded T, see example in **Fig. 1h**). To prevent inflation of the threshold value in recordings with dense epileptiform activity, we imposed a numerical upper bound on its value (500 µV for hippocampal and 400 µV for cortical electrodes).
2. For each crossing, we identified the local minimum within a 1 s window centered on the crossing and defined an individual discharge (**Fig. 1h**) as the 500 ms trace centered around this minimum. Deflections occurring within ∼0.5 s of a larger deflection were not counted as separate discharges. The mean discharge rate was calculated as the total number of discharges divided by recording duration. The maximal discharge rate per hour was obtained by sliding a 1 h window across the recording and selecting the window with the highest discharge count.
3. We computed short-time Fourier transforms of the EEG signals using MATLAB’s spectrogram function with 1 s Hamming windows and 50% overlap. Power was extracted in the Delta (1–4 Hz), Theta (4–7 Hz), Alpha (8–12 Hz), Sigma (12–16 Hz), and Beta (24–30 Hz) bands. Power was summed across the frequencies of each band in each spectrogram time bin, and for each band, a threshold was defined as mean + 3 × standard deviation over the entire recording.
4. To detect candidate EDEs, we defined a binary time series marking the occurrence of each detected discharge peak and convolved it with a 5-min rectangular window, yielding the number of discharges within ±2.5 min of each time point. A threshold for this convolved signal was set to RMS + 2 × standard deviation over the entire recording, with an upper bound of 30 discharges.
5. Candidate events were defined as time intervals where the convolved discharge signal from step 4 crossed and remained above the threshold; supra-threshold segments separated by less than 5 min were merged into a single candidate event.
6. A candidate event was classified as an EDE if it satisfied all of the following:

a. duration of the candidate event, from the first to the last supra-threshold time point, > 5 min;
b. in each of the five frequency bands from step 3, the maximum band power across the spectrogram windows within the event exceeded that band’s threshold (the maximum need not occur in the same window for different bands);
c. number of step-2 discharges within the event divided by its duration > 8 discharges/min;
d. temporal overlap with an event meeting criteria a–c in the other electrode.

#### Threshold sweep

The discharge-density and duration criteria in step 6 were derived from our own dataset rather than adopted from published seizure or other epileptiform activity definitions. To test whether the conclusions depended on the values selected, we recomputed EDE rate for every recording at each of 84 combinations of the density criterion (step 6c: 2, 3, 4, 5, 6, 8, 10, 12, 15, 20, 25, 30/min) and the minimum-duration criterion (step 6a: 1, 2, 3, 5, 7, 10, 15 min). Every other step of the pipeline was held unchanged: discharge detection, the 5-min convolution (step 4), merging of intervals separated by <5 min (step 5), the spectral criterion, and the two-electrode concurrency requirement. The 5-min integration window itself was not varied. Recomputed counts at 8/min and 5 min matched the original pipeline in all 421 recordings. At each combination we computed: the area under the ROC curve for discriminating pilocarpine-treated post-injection recordings from control recordings by EDE rate (n = 236 pilocarpine-treated post-injection and 185 control recordings); the specificity and sensitivity of detecting any EDE (**Fig. 4e**). The plateau was defined as combinations with AUC ≥97% of the grid maximum. In the 17 mice used for the predictive analysis, we also recomputed at each combination: the Spearman correlation between maximal acute-phase PSA-AP and maximal chronic-phase EDE rate; the apparent AUC of the combined Firth model (maximal acute PSA-AP and maximal acute EDE rate) predicting chronic severity. The outcome was either held at its definition at 8/min and 5 min, or rederived at each combination by classifying the 8 mice with the highest maximal chronic EDE rate as severe (**Supp. Fig. 6**).

Combinations at which fewer than three mice had acute or chronic EDEs were not evaluated. Because each mouse contributed multiple recordings, discrimination was also computed per mouse from each mouse’s mean post-injection EDE rate (20 pilocarpine-treated and 13 vehicle-treated mice).

#### EDE counting

Each temporally overlapping pair of hippocampal and cortical events was counted as one EDE, with EDE duration defined as their temporal overlap. EDE rate was the number of EDEs per hour of recording, and cumulative EDE duration the sum of EDE durations per recording. Because a long event in one electrode can overlap two separate events in the other, this pair-based count can count one episode twice; all analyses were therefore repeated after merging overlapping pairs that shared an event into single episodes. Events meeting criteria 6a–c in a single electrode, without the concurrency requirement, are reported separately as regional EDEs (**Supp. Fig. 7b**). Because EDE boundaries derive from the 5-min integrated discharge signal (step 4), EDE durations include part of the integration window and exceed the duration of the underlying discharge activity.

#### Seizure-criterion segments and expert review

Because the primary detector counts at most one discharge per ∼0.5 s, ictal-frequency activity within EDEs was quantified using a separate analysis scheme.

- Within each single-electrode event contributing to an EDE, discharges were re-counted as individual excursions below the step-1 threshold, with hysteresis: after an excursion below −T, the next excursion was counted only once the signal had recovered above −T/2, and excursions separated by <50 ms were merged.
- Discharge rate was computed in sliding 5-s and 10-s windows, and each event was tested against three published criteria for electrographic seizures: >3 Hz for >5 s^40^, ≥3 Hz for ≥10 s^44^, and >2.5 Hz averaged over 10 s^24^.
- An EDE was scored as containing a seizure-criterion segment if its hippocampal event, its cortical event or both met the criterion; each EDE was counted once (breakdown by electrode in **Supp. Table 11**). The ≥3 Hz for ≥10 s criterion^44^ was used for all analyses because it flagged no control EDE and was the most specific on expert review (see below); results for all three criteria are given in **Supp. Table 11**.
- Segment duration was measured from the first to the last discharge of the longest continuous segment at ≥3 Hz.

All EDEs flagged by any of the three criteria were reviewed by an expert epileptologist blinded to group, phase and analysis output, from images showing the cortical and hippocampal traces on a common time axis and voltage scale (60-s overview and 12-s zoom). Flagged events within 45 s of each other in the same recording were shown once, giving 77 flagged windows, and 25 windows from EDEs meeting no criterion were mixed in as internal controls. Each window was scored for seizure (a clearly evolving ictal pattern, or a segment of ≥3 Hz discharges lasting ≥10 s) and, separately, for artefact; borderline windows were scored as not seizures. An EDE was considered confirmed if a window containing one of its flagged events was scored as a seizure. Of the 52 windows flagged by the ≥3 Hz for ≥10 s criterion, 37 (71%) were scored as seizures, compared with 8 of 25 windows flagged only by the other two criteria (0 of 3 from vehicle-treated mice) and 1 of 25 control windows. Of the 68 EDEs meeting the ≥3 Hz for ≥10 s criterion, 48 were confirmed as seizures; of the 20 not confirmed, 4 were scored as borderline, and 16 were obscured by recording irregularities. Such irregularities, typically non-cerebral transients from movement, chewing or grooming, are common in freely moving rodents^45, 81^ and can inflate discharge counts; automated detections therefore require visual verification^82^. These 16 EDEs (1.7% of post-induction EDEs, from 8 of 236 recordings) were not counted as seizure-containing, and none fell in the session that set a mouse’s maximal acute- or chronic-phase EDE rate, so they do not affect the severity classification or the predictors. **Fig. 4c** and the seizure analyses in Results use only the 48 confirmed EDEs; counts flagged by each automated criterion before review are given in **Supp. Table 11**.

### Cross-modal analyses

For same-mouse correlations (**Fig. 5h–m, Supp. Fig. 11a–f**), each TPLSM session was paired with the EEG session recorded from the same mouse in the same weekly bin (acquired on a separate day), giving 238 pairs from 19 mice.

### Predictive Modeling

For predictive analyses, we included the 17 pilocarpine-treated mice with at least two TPLSM and two EEG sessions in the acute phase (days 0–26) and at least two EEG sessions in the chronic phase (**Supp. Table 1**). These mice were divided into severe and non-severe chronic epilepsy groups based on chronic-phase epileptiform burden (Weeks 7–16 post-pilocarpine), defined from EEG data alone so that the outcome was independent of the metrics being tested as predictors. Mice whose maximal chronic-phase EDE rate exceeded 0.25/h were classified as severe (lowest severe, 0.28/h; highest non-severe, 0.235/h); all others were classified as non-severe.

Because PSA% and PSA-AP varied between recordings within the same animal, the maximum value of each metric across the acute period (pilocarpine day to Week 3) was used to represent that animal’s pathology severity. Maximal acute-phase values were likewise used for the remaining candidate predictors: average neuronal firing rate, EDE rate, and cumulative EDE duration. All variables were treated as continuous and summarized as medians with interquartile ranges, and group differences were assessed with the Wilcoxon rank-sum test.

Candidate predictors were screened for complete or quasi-complete separation prior to modeling. Given the relatively small sample size and the potential for separation, all regression models used Firth’s penalized logistic regression (logistf package in R), which yields bias-reduced maximum likelihood estimates. Univariate Firth models were fit for each acute-phase predictor (Day 0–Week 3), and odds ratios, 95% confidence intervals, and receiver operating characteristic (ROC) curves with their AUC were calculated. PSA-AP and EDE rate were the only predictors to reach significance (**Supp. Table 12**); these two were carried forward into three models: 1) a PSA-AP univariate model, 2) an EDE-rate univariate model, and 3) a combined multivariate model including both predictors.

Model performance was evaluated using AUC, sensitivity, specificity, and accuracy at the threshold maximizing Youden’s index; these thresholds were derived from the full dataset, so the corresponding classification metrics are apparent rather than cross-validated. Three complementary checks were then applied. Calibration of the combined model was assessed with the Hosmer–Lemeshow goodness-of-fit test, grouping animals into five bins of predicted risk and plotting the observed proportion of severe mice against the mean predicted probability within each bin. Optimism was estimated by bootstrap resampling (1,000 resamples; Harrell’s method): each model was refit on a resample of mice drawn with replacement and evaluated on both that resample and the original dataset; the mean difference between the two AUCs defined the optimism, which was subtracted from the apparent AUC to give a bias-corrected estimate. Separately, out-of-bag performance was estimated by fitting each model on a bootstrap resample and computing its AUC on the mice not drawn into that resample; resamples whose out-of-bag set lacked either outcome class were redrawn until 1,000 valid AUCs were obtained (**Supp. Fig. 12d**). Generalization was assessed by leave-one-out cross-validation, in which each mouse was held out in turn, the model was refit on the remaining mice, and the held-out predictions were pooled to construct a cross-validated ROC curve. To assess when predictive information emerged, the three models were refit using maximal values from days 0–1, 2–9 and 10–26 after induction and compared with the full acute period (days 0–26); 95% confidence intervals for these AUCs were computed with DeLong’s method (**Supp. Fig. 12a–b**). ROC curves and AUC values were computed with the pROC package.

### Statistical analyses

All statistical analyses were performed using GraphPad Prism (version 10), Matlab (R2023a and R2025a), and R (version 4.4.2). Correlations were assessed by linear regression, Pearson’s, or Spearman correlations, and p-values and 95% confidence intervals were reported. Repeated-measures data were analyzed using linear mixed-effects models (group, time and their interaction as fixed effects; mouse as a random effect; REML estimation), which accommodate missing sessions, followed by Tukey’s HSD post-hoc tests. Per-mouse comparisons used the Wilcoxon signed-rank test (paired, across phases) or the Mann–Whitney U test (between groups). For phase-level summaries, the maximal value of each metric was taken for each mouse within the acute (induction day to Week 3), latent (Weeks 4–6) and chronic (Weeks 7–16) periods; because weekly bins span ±2 days, these correspond to days 0–26, 27–45 and 46–116 after induction. For weeks where vehicle data was lacking (less than 3 mice per recording date), we performed no statistical comparisons across groups (week 16 in **Fig. 2d-e**). For datasets involving comparison of a single factor, one-way ANOVA was used, followed by Tukey’s HSD post hoc correction. Statistical significance was defined as *p<*0.05. Data are reported as mean ± SEM, and *n* denotes the number of animals per group. The specific statistical test that was used for each comparison is indicated in the corresponding figure or table legends. No statistical method was used to predetermine sample size. Mice were randomly assigned to pilocarpine or vehicle.

## Data Availability

The data supporting the findings of this study will be made available by the corresponding authors upon reasonable request.

## Author Contributions

N.G. and A.K. performed the surgeries with training from M.R.C., S. Stanko, and design advice from H.D. and D.D.; N.G., G.T., A.K., H.T. performed optical and EEG recording experiments; H.T., D.A., G.T., S.K., M.R.C., J.I., S.R., S.B., H. Shindler, J.M., P.A., S. Singh, H.D., and D.D. performed data analyses. J.I., C.R., and S. Stanko performed colony maintenance and genotyping; N.G. S.R. and S. Stanko did histological analyses; A.A. blindly evaluated seizure detection accuracy; H. Suh., A.A., D.S., and I.N. contributed to the conceptual development of the project and provided feedback for EEG analyses; D.D. and H.D., wrote the manuscript with contributions and figure preparation from N.G., H.T. and D.A., and comments from all authors; H.D. and D.D. conceived the project, designed experiments and analyses, and supervised the study.

