## Supplementary Figures and Tables for "Early Increases in Hippocampal Synchrony Predict Chronic Epileptiform Burden in a Temporal Lobe Epilepsy Model"

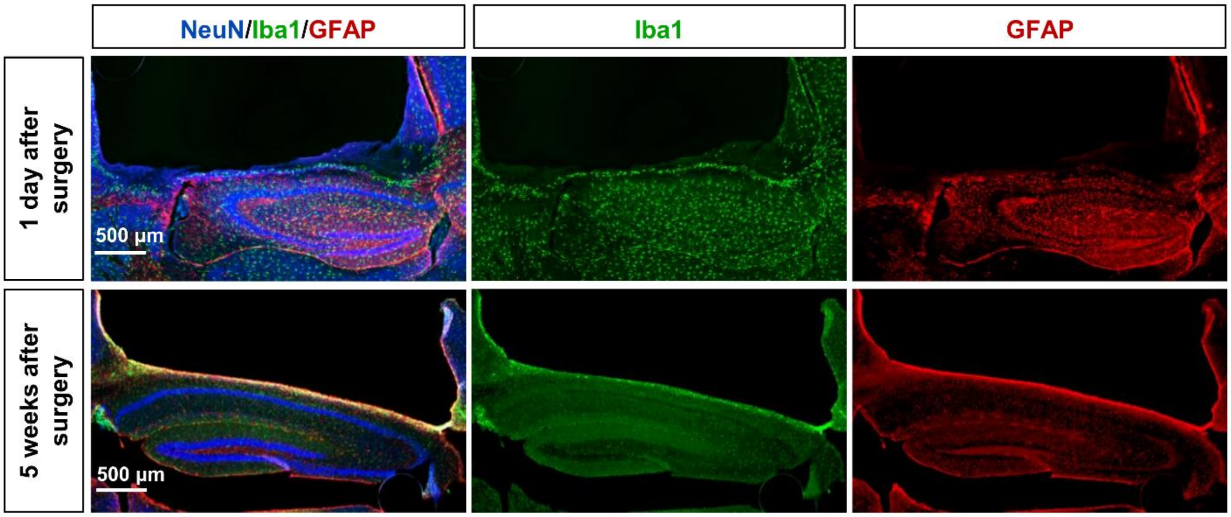


**Supplementary Figure 1. Histological evaluation of glial responses and hippocampal window placement above the hippocampus.**

Epifluorescence microscopy images of coronal brain sections after craniotomy and window placement above the hippocampus, stained with NeuN (neuronal nuclei, blue), IBA1 (microglia, green) and GFAP (astrocytes, red). The extensive immune (Iba1) and astroglial (GFAP) responses stimulated acutely by the surgical procedure (day 1) subside, revealing a quiescent glial environment throughout the hippocampus 5 weeks later.

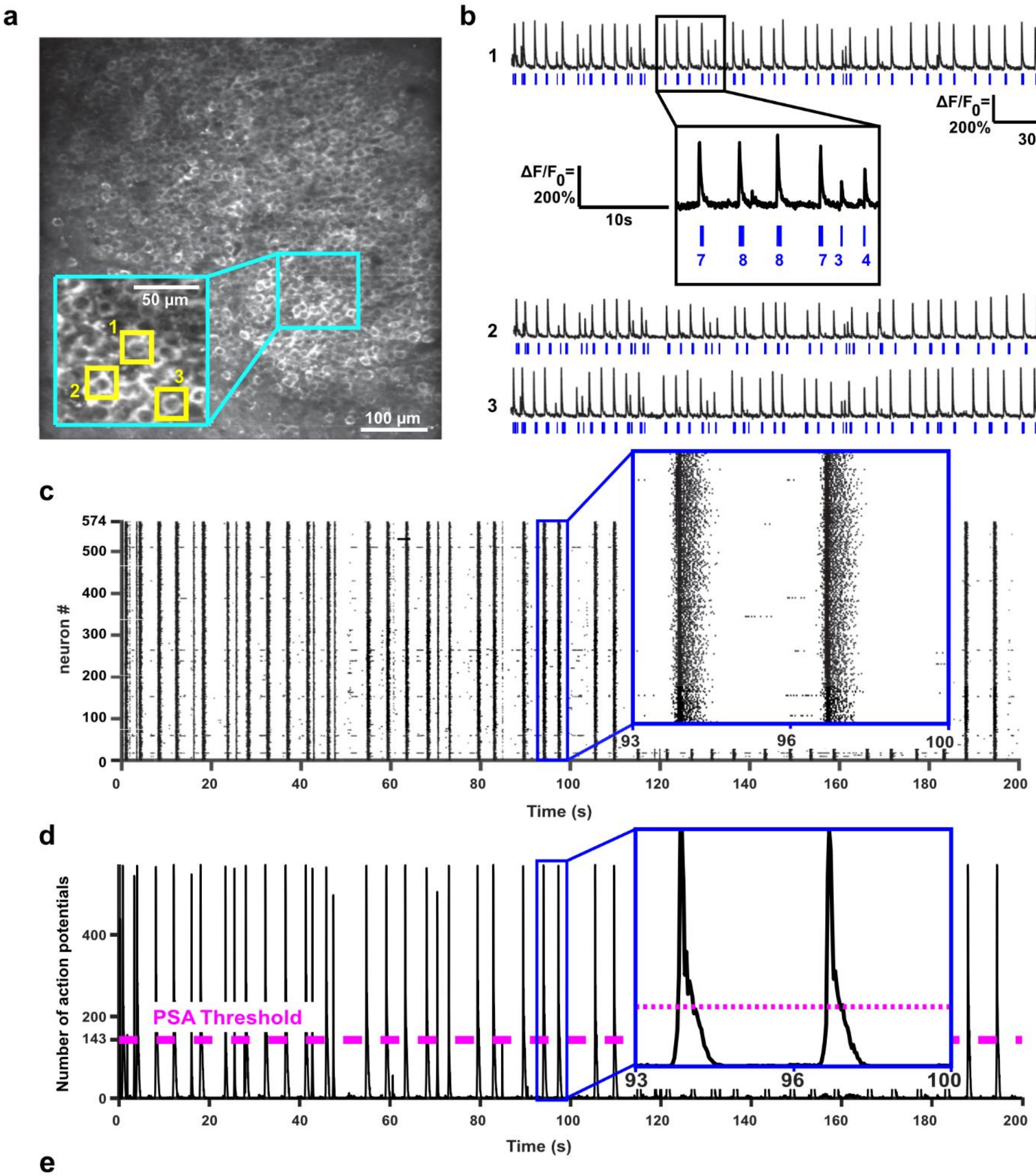


**e**

$$\text{PSA-AP:} \quad \frac{\text{PSA APs: 197,140}}{\text{Total APs: 205,208}} * 100 = 96.07\%$$

$$\text{PSA \%:} \quad \frac{\text{PSA Frames: 209}}{\text{Total Frames: 6000}} * 100 = 3.483\%$$

**Supplementary Figure 2 (relates to Fig. 2). Example of how PSA-AP and PSA% values are calculated.**

**(a)** Example FOV of jRGECO1a-expressing CA1 neurons. The inset (cyan) shows a magnified region with three example cells highlighted.

**(b)** $\Delta F/F_0$ traces for the three neurons shown in **a** (black traces), with APs extracted by the MLSpike algorithm (blue ticks). The inset shows a magnified portion of the trace with numbers indicating the number of APs detected in each burst.

**(c)** Raster plot summarizing AP firing from all active neurons within the FOV shown in (**a**) during a single recording. The presence of vertical lines indicates synchronous firing by most neurons. The inset shows a magnified portion of the raster plot.

**(d)** Summed AP counts across all neurons for the raster plot shown in **c**. The threshold for defining a PSA event is set to 143 APs (rounded 25% of the 574 neurons recorded within the FOV; dashed magenta line). The inset shows a magnified portion of the plot.

**(e)** Example calculations of PSA-AP and PSA% for the FOV and data presented in **a-d**.

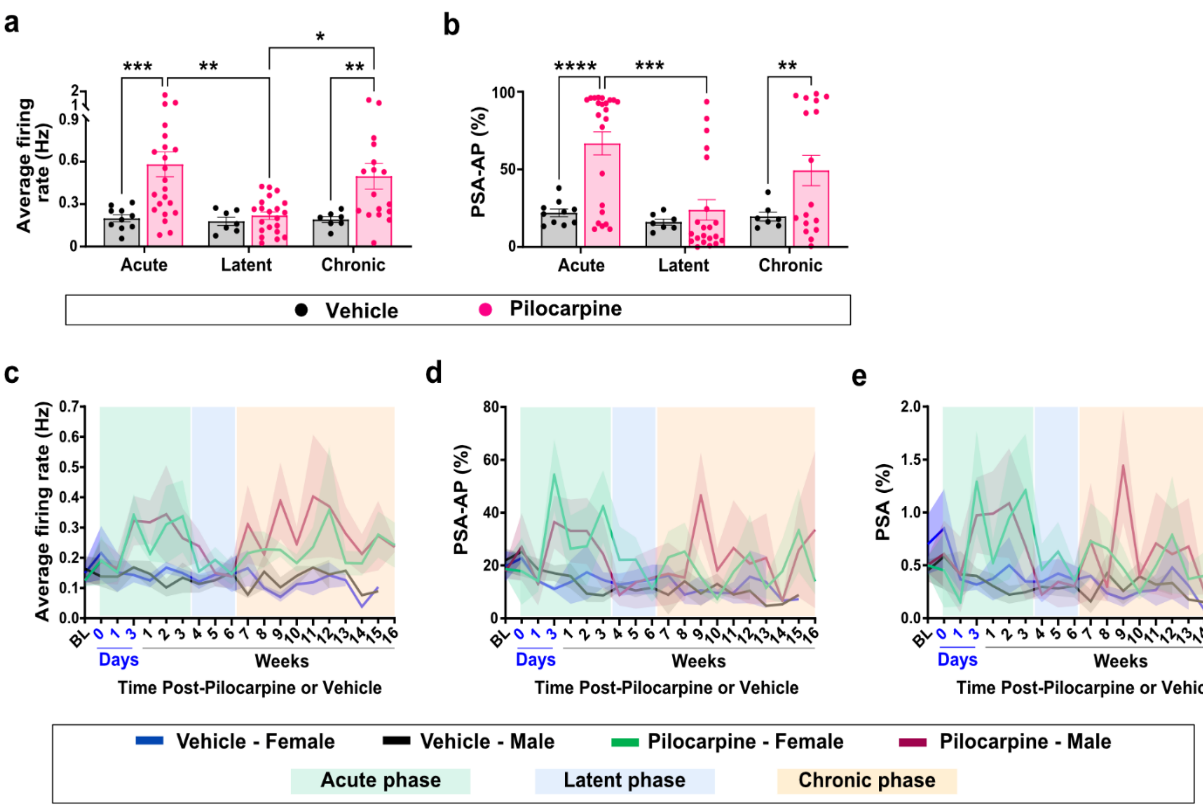


**Supplementary Figure 3 (relates to Fig. 2). Neuronal activity in pilocarpine-treated mice stratified by disease phase and sex.**

**(a–b)** Same data as in **Fig. 2d–e**, grouped by disease phase: acute (day of pilocarpine to Week 3), latent (Weeks 4–6), and chronic (Weeks 7–16). Pilocarpine-treated mice showed significant increases relative to vehicle-treated mice during the acute and chronic periods, but not during the latent period. Each dot represents the maximal value for one mouse within each phase, with each session's value taken as the maximum across recorded FOVs. Error bars indicate mean ± SEM. *,p<0.05; **,p<0.01, ***,p<0.001, and ****, p<0.0001.

**(c–e)** Average firing rate (**c**), PSA-AP (**d**), and PSA% (**e**) over time, separated into male and female groups. No significant sex differences were detected in average firing rates or synchrony metrics. Shaded background regions denote disease phases: acute (green), latent (blue), and chronic (orange). Mouse numbers and statistical analyses are the same as in **Fig. 2d-f.** A complete list of statistical comparisons and mouse numbers in each recording week are provided in **Supp. Table 4**.

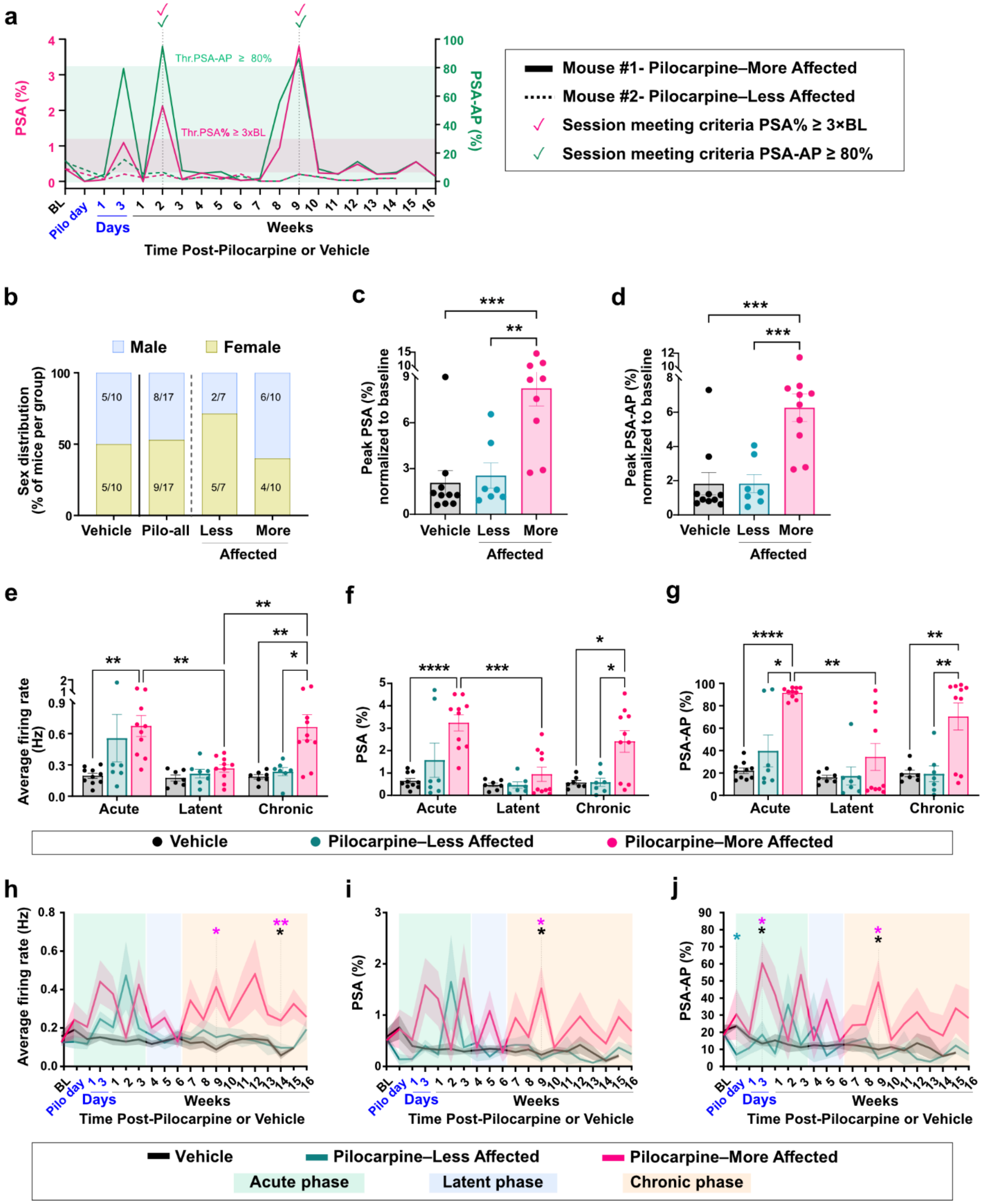


**Supplementary Figure 4 (relates to Fig. 2). Classification strategy and neuronal activity across disease phases and severity groups in pilocarpine-treated mice.**

**(a)** Example of the classification of mice into the less-affected and more-affected groups. PSA% and PSA-AP traces from two representative mice are shown (maximal values per recording session). Thresholds are as defined in Methods (pink shading, individualized PSA% threshold for mouse #1; green shading, fixed PSA-AP threshold). Mice meeting both criteria were classified as more-affected (solid lines; mouse #1); all others as less-affected (dashed lines; mouse #2).

**(b)** Distribution of male and female mice across the experimental and severity groups.

**(c-d)** The more-affected group showed significantly higher PSA% and PSA-AP values than both the less-affected and vehicle groups. The latter two groups did not differ significantly (n=10 more-affected, 7 less-affected, 10 vehicle; each dot represents the maximal value for a single mouse, normalized to baseline, error bars indicate mean ± SEM; one-way ANOVA with Tukey HSD *post-hoc* correction).

**(e-g)** Average firing rate (e), PSA% (f) and PSA-AP (g) in the vehicle, less-affected and more-affected groups, binned by disease phase. Each dot represents the maximal value for a single mouse during the respective phase. The same mice as in c–d are shown, with the same analysis as in Fig. 2d–f.

**(h-j)** Temporal evolution of average firing rate (h), PSA% (i) and PSA-AP (j) in the severity and vehicle groups across the entire recording period. The same mice as in **c–d** are shown, with the same analysis as in **Fig. 2d–f**. Magenta asterisks indicate significance of vehicle vs. more-affected groups; green asterisks indicate significance of vehicle vs. less-affected groups; black asterisks indicate significance of more-affected vs. less-affected groups.

*,$p<0.05$; **,$p<0.01$; ***,$p<0.001$; ****,$p<0.0001$. A complete list of statistical comparisons is provided in **Supp. Table 5**.

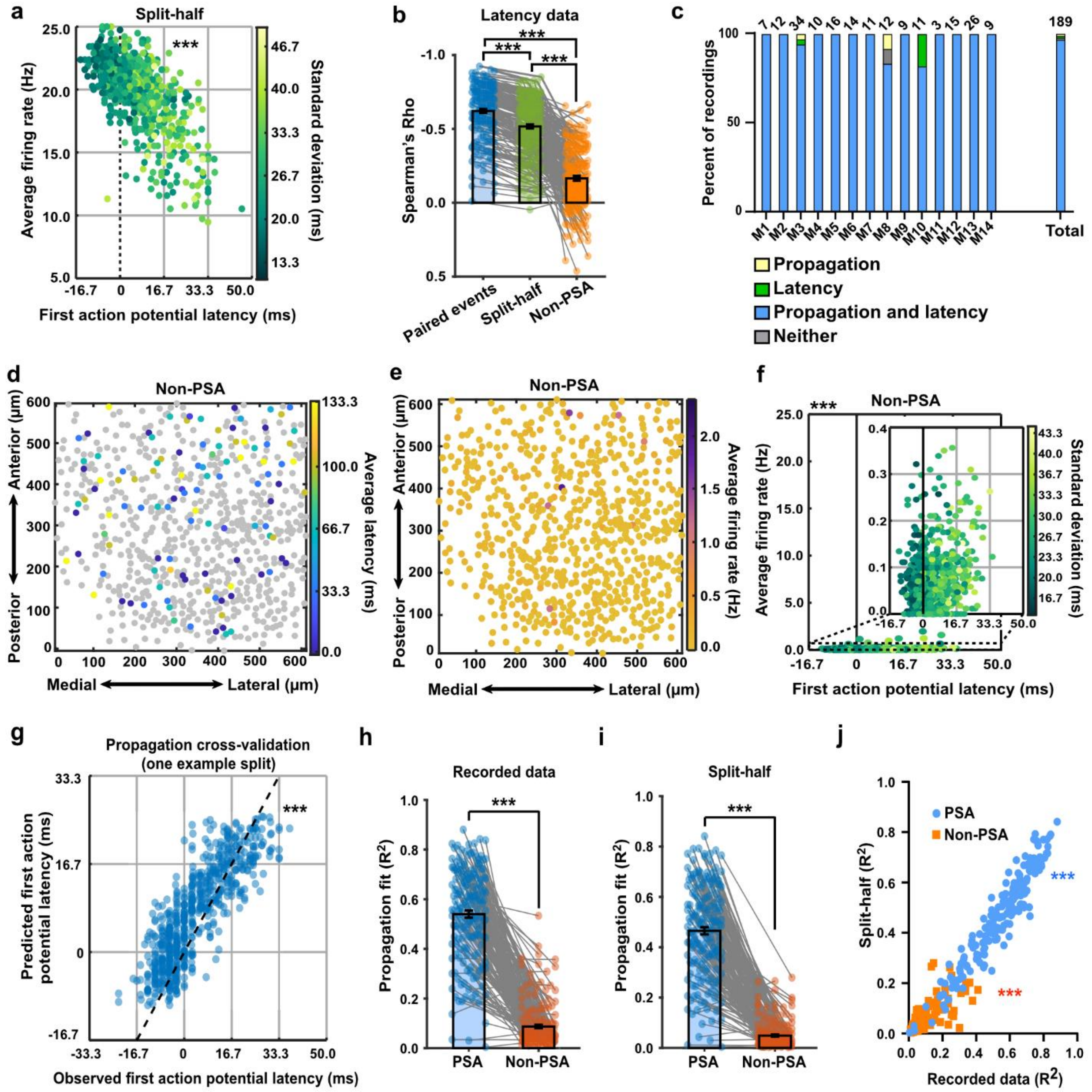


**Supplementary Figure 5 (relates to Fig. 3). Spatial firing rate gradients and latency to activity effects are specific to PSA events**

**(a)** Scatter plot of all neurons in the same FOV as in **Fig. 3e** (n=650) showing that the relationship between each neuron's mean firing rate during PSA frames and its mean latency to the first AP during detected PSA events is not driven by shared events. Firing rates were calculated from a randomly selected half of detected PSA events, whereas first-AP latencies were calculated from the remaining half. This split-half analysis was repeated across 100 random permutations. Dots are color-coded by the standard deviation of each neuron's first-AP latency, showing that the greater latency variability among neurons with lower firing rates was preserved in the split-half analysis.

**(b)** Comparison of Spearman's rho values quantifying the association between first-AP latency and mean firing rate across recordings. Correlations were calculated using firing rates measured during the same PSA events used to determine first-AP latency ("paired events," as in **Fig. 3e**), using independent halves of PSA events for the two measures ("split-half events," as in panel **a**), or using firing rates measured during non-PSA frames ("non-PSA," as in panel **f**). Dots represent individual recordings; lines connect values from the same recording across conditions. Mean Spearman's rho values were −0.621±0.0136 (mean±SEM) for paired events, −0.517±0.014 for split-half events, and −0.166±0.0161 for non-PSA firing rates. All pairwise comparisons were significant ($p<0.001$ repeated-measures one-way ANOVA with Tukey's HSD post hoc correction; error bars indicate mean±SEM; n=189 recordings).

**(c)** Summary of detected types of correlations during PSA events across all analyzed mice. Every mouse exhibited either exclusively, or a large majority of, PSA events that combined a propagating wave ("propagation") with a correlation between first-AP latency and the same-neuron average firing rate ("latency").

**(d)** Map of the same FOV as in **Fig. 3c** showing the locations of 650 somata, with each neuron color-coded by the mean latency from the onset of randomly selected non-PSA control events to the time of its first AP firing (in sample $R^2$ = 0.0302, propagation permutation test p-value = 0.1905, n=109 participating cells).

**(e)** Map of the same FOV as in **Fig. 3d** showing each neuron color-coded by its average firing rate during the non-PSA frames only. The firing rate gradient magnitude significantly exceeded the null distribution generated from shuffled data, but the magnitude and directions were substantially smaller and different than during the PSA frames, respectively (n=2000 permutations, $p<0.001$).

**(f)** Scatter plot of all neurons in the same FOV as in **Fig. 3e** (n=650), showing each neuron's average firing rate outside PSA frames plotted against its mean first-AP latency during the detected PSA events. The relationship seen during PSA frames was absent: the correlation was weak and in the opposite direction (Spearman's $\rho=+0.1773$, $p=5.45\times10^{-6}$). Dots are color-coded by the standard deviation of each cell's latency.

**(g)** Representative split-half cross-validation example for the same recording shown in **Fig. 3c**. PSA events were randomly divided into two non-overlapping halves; a spatial latency gradient was fitted to one half and the resulting coefficients applied to the neuron coordinates of the held-out half to generate predicted latencies (Spearman correlation $R^2$ = 0.6986, Spearman's $\rho$ = 0.8358, p = $6.350\times10^{-171}$).

**(h)** Comparison of the propagation goodness of fit ($R^2$) values for all PSA events in a recording compared with those obtained from an equal number of randomly selected non-PSA events from the same recordings. Analysis was performed across 200 random permutations. PSA mean $R^2$=0.54, Non-PSA $R^2$=0.087. (paired two-tailed t-test, p=$4.82\times10^{-66}$; n=174 recordings, lines connect the pairs).

**(i)** Comparison of the propagation goodness of fit ($R^2$) values for split-half analysis of all PSA events (PSA column, 200 permutations) and an equal number of random non-PSA events within the same recordings (non-PSA column, 200 permutations). The split-half analysis maintained approximately 85% of the $R^2$ value, while the fit quality decreased for non-PSA frames (PSA mean half-split Spearman correlation $R^2$=0.465, Non-PSA $R^2$=0.049; paired two-tailed t-test, p=$7.36\times10^{-64}$; n=172 recordings).

**(j)** Relationship between paired events regression $R^2$ and split-half cross-validation $R^2$ (panels **h** and **i**, respectively) for the same recordings, shown separately for PSA events and matched non-PSA windows (n=172 recordings). For both PSA and non-PSA events, recordings with better paired event fits also yielded better $R^2$ values in split-half analysis (PSA events: Spearman's $\rho$=0.9508, non-PSA events: $\rho$=0.8253; p<0.0001 for both fits).

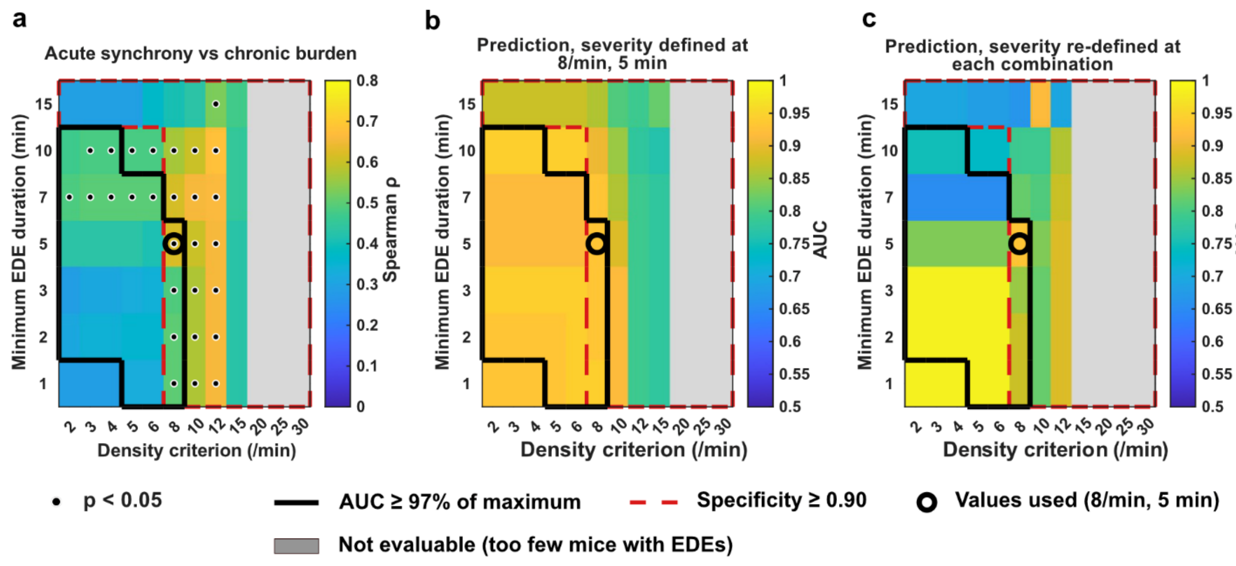


**Supplementary Figure 6 (relates to Figs. 4 and 6). The synchrony–burden correlation and severity prediction hold across EDE detection criteria.**

Each cell represents a combination of density and duration criteria used for the analysis; outlines and circles are as in **Fig. 4e**. Data are from n = 17 pilocarpine-treated mice included in the predictive analysis (Methods). Grey cells were not evaluated because fewer than three mice exhibited acute or chronic EDEs meeting the corresponding criteria.

**(a)** Colors represent the Spearman correlation coefficient between maximal acute-phase PSA-AP and maximal chronic-phase EDE rate; dots indicate cells with significant correlations ($p < 0.05$). Significant correlations were observed for all EDE duration criteria from 1 to 10 min at density criteria of 8–12 discharges/min ($\rho$ = 0.49–0.68). Correlations were weaker at lower density criteria ($\rho$ = 0.29–0.50).

**(b)** Apparent AUC of the combined Firth logistic model (maximal acute PSA-AP and EDE rate) for predicting chronic severity, recomputed at each combination of EDE density and duration criteria, with severity defined at the selected criteria (8 discharges/min, 5 min). Within the black outline, the combinations that best separated pilocarpine-treated from control recordings (**Fig. 4e**), AUC ranged from 0.92 to 0.94; the circle marks the selected criteria.

**(c)** As in **b**, with chronic severity groups re-derived at each combination by classifying the 8 mice with the highest maximal chronic EDE rate as severe. AUC values ranged between 0.79–0.93 for density criteria of 8–12/min and durations of 1–10 min. Where fewer than 8 mice had any chronic EDE, the outcome could not be re-derived (grey cells).

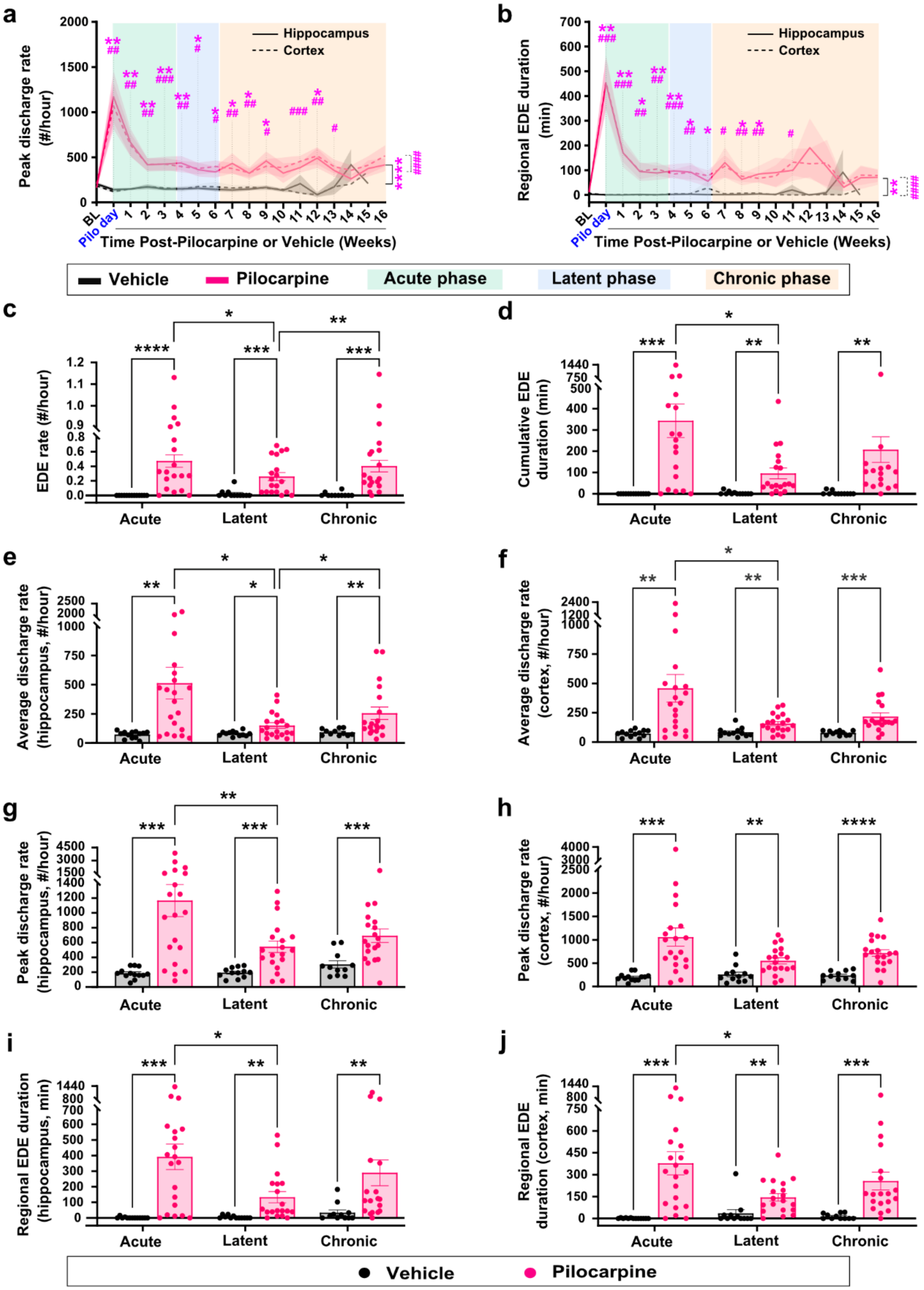


**Supplementary Figure 7 (relates to Fig. 5). Additional EEG measures of pilocarpine-induced epileptiform burden.**

**(a)** Maximal discharge rate of detected discharges (as shown in **Fig. 1h**) within a 1 h window for hippocampal and cortical electrodes. Both electrodes show a marked increase during the first week after pilocarpine induction, which largely decays by 2 weeks but remains significantly above vehicle levels. Cortical and hippocampal traces are qualitatively similar.

**(b)** Cumulative duration of regional EDEs per recording, shown separately for the hippocampal and cortical electrodes. Regional EDEs meet the duration, spectral, and discharge-rate criteria within a single region and are not required to be synchronous across regions (see **Fig. 4b** and Methods). Unlike cumulative EDE duration (**Fig. 5e**), which is the summed temporal overlap of hippocampal and cortical events, this metric is measured independently for each region and is therefore not directly comparable in magnitude. Both electrodes exhibit a temporal pattern similar to (**a**) and **Fig. 5d-e**: an acute increase during the week following pilocarpine induction, followed by a decline through the latent period that remains significantly above vehicle levels, and continued elevation above vehicle levels during the chronic period.

For data in (**a–b**), mouse numbers and statistical analyses are the same as in **Fig. 5c–e**. A complete list of statistical comparisons and mouse numbers in each recording week are provided in **Supp. Table 6**. Significance markers are as defined in **Fig. 5c–e**.

**(c)** EDE rate by disease stage, showing significantly higher rates in pilocarpine-treated than control mice across the entire study. Within pilocarpine-treated mice, maximal EDE rates during the latent period are significantly lower than during the acute and chronic periods.

**(d)** Cumulative EDE duration by disease stage, showing trends similar to (**c**).

**(e–h)** Average and peak discharge rates by disease stage for the hippocampal and cortical electrodes.

**(i–j)** Cumulative regional EDE duration by disease stage for each electrode.

For data in **c–j**, each dot represents the maximal value of the corresponding metric from a single mouse within a given disease phase. Bars indicate mean ± SEM. Mouse numbers and statistical analyses and significance markers are the same as in **Fig. 5c–e**. A complete list of statistical comparisons is provided in **Supp. Table 6**.

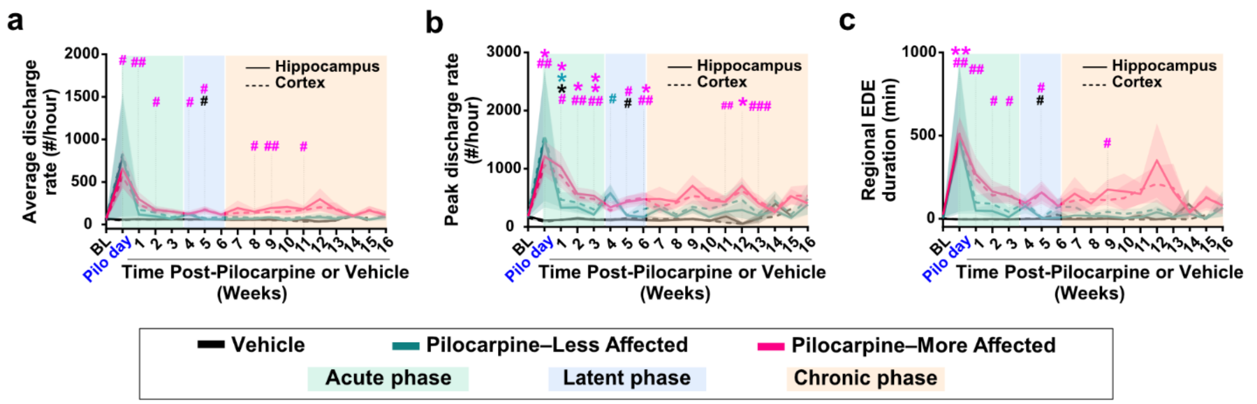


**Supplementary Figure 8 (relates to Fig. 5). EEG measures of pilocarpine-induced pathology, stratified by severity groups.**

**(a–c)** Average discharge rate (**a**), peak discharge rate (**b**) and regional EDE duration (**c**) (same data as in **Fig. 5c** and **Supp. Fig. 7a–b**), stratified by severity group. EDE rate and cumulative EDE duration by severity group are shown in **Fig. 5f–g**. Although severity groups were defined based on TPLSM data, EEG metrics show parallel differences: the more-affected group exhibits an acute increase, followed by a latent-period decrease and sustained elevation during the chronic period. The less-affected group shows consistently lower values without a chronic-period increase.

Data presented as mean ± SEM (solid lines: hippocampus; dashed lines: cortex; shaded areas indicate disease phases). Vehicle: n=13; pilocarpine: n=16 (6 less-affected, 10 more-affected). One less-affected mouse contributed imaging but not EEG data and is therefore included in the TPLSM-based analyses (17 pilocarpine mice; 10 more-affected, 7 less-affected) but not here. Statistical analyses were performed using mixed-effects models with restricted maximum likelihood and Tukey's HSD post-hoc corrections. (*) and (#) indicate significance in hippocampal and cortical data, respectively. Magenta: vehicle vs. more-affected; green: vehicle vs. less-affected; black: more-affected vs. less-affected. * or #,$p<0.05$; ** or ##,$p<0.01$; *** or ###,$p<0.001$. A complete list of significant comparisons is provided in **Supp. Table 7**.

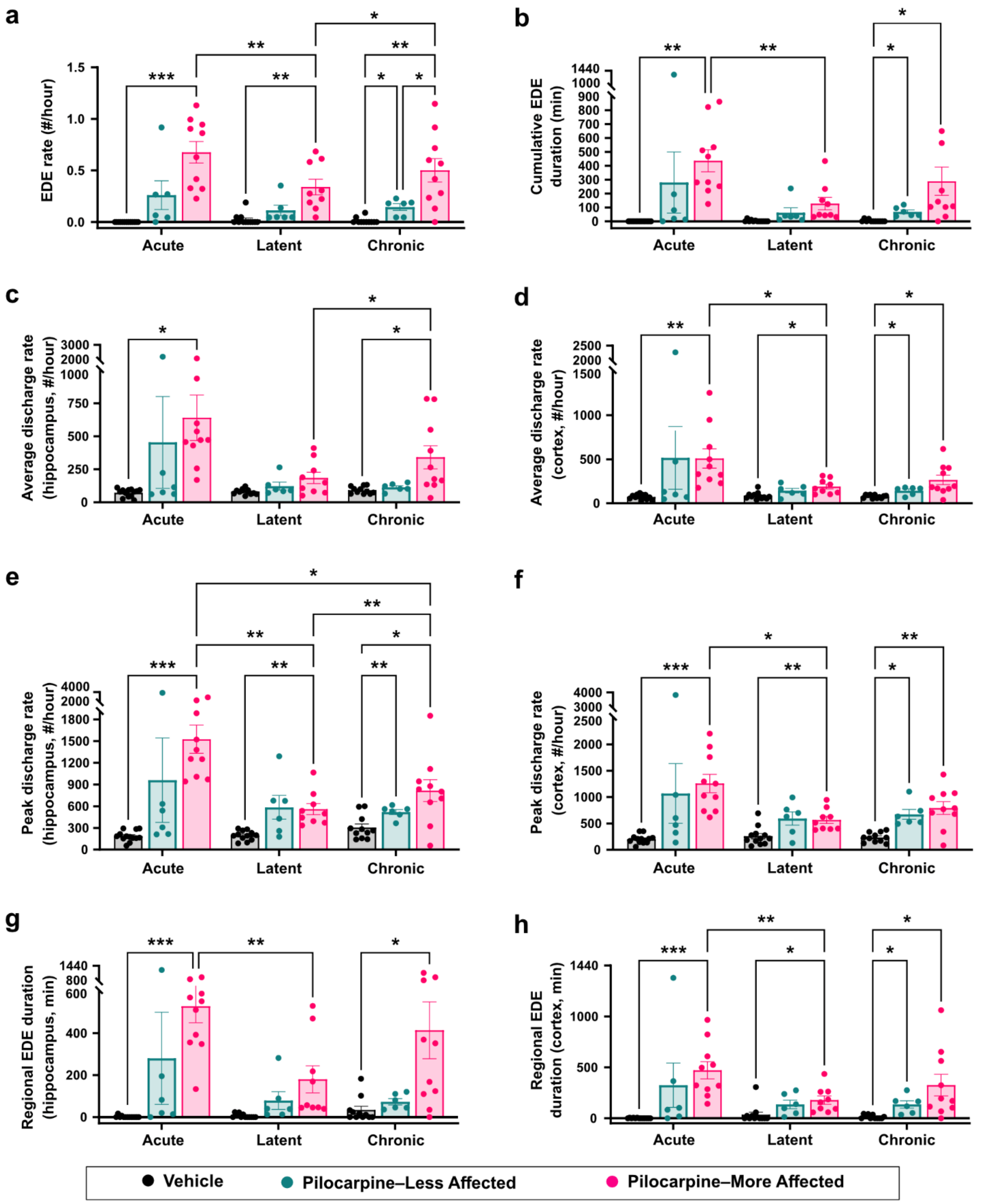


**Supplementary Figure 9 (relates to Fig. 5). EEG measures of pilocarpine-induced pathology across disease phases and severity groups.**

(**a-h**) EEG metrics (same data as in **Fig. 5f–g** and **Supp. Fig. 8a–c**) stratified by severity group (vehicle, less-affected, and more-affected). By EEG measures, the less-affected group remains distinguishable from vehicle, while significant differences between less-affected and more-affected groups emerge predominantly during the chronic period. Dots represent maximal measured values per period from individual mice; bars indicate mean±SEM. Vehicle: n=13; pilocarpine: n=16 (6 less-affected, 10 more-affected). One less-affected mouse contributed imaging but not EEG data and is therefore included in the TPLSM-based analyses (17 pilocarpine mice; 10 more-affected, 7 less-affected) but not here. Statistical analyses as in **Fig. 5c–e**; A complete list of significant comparisons is provided in **Supp. Table 8**.

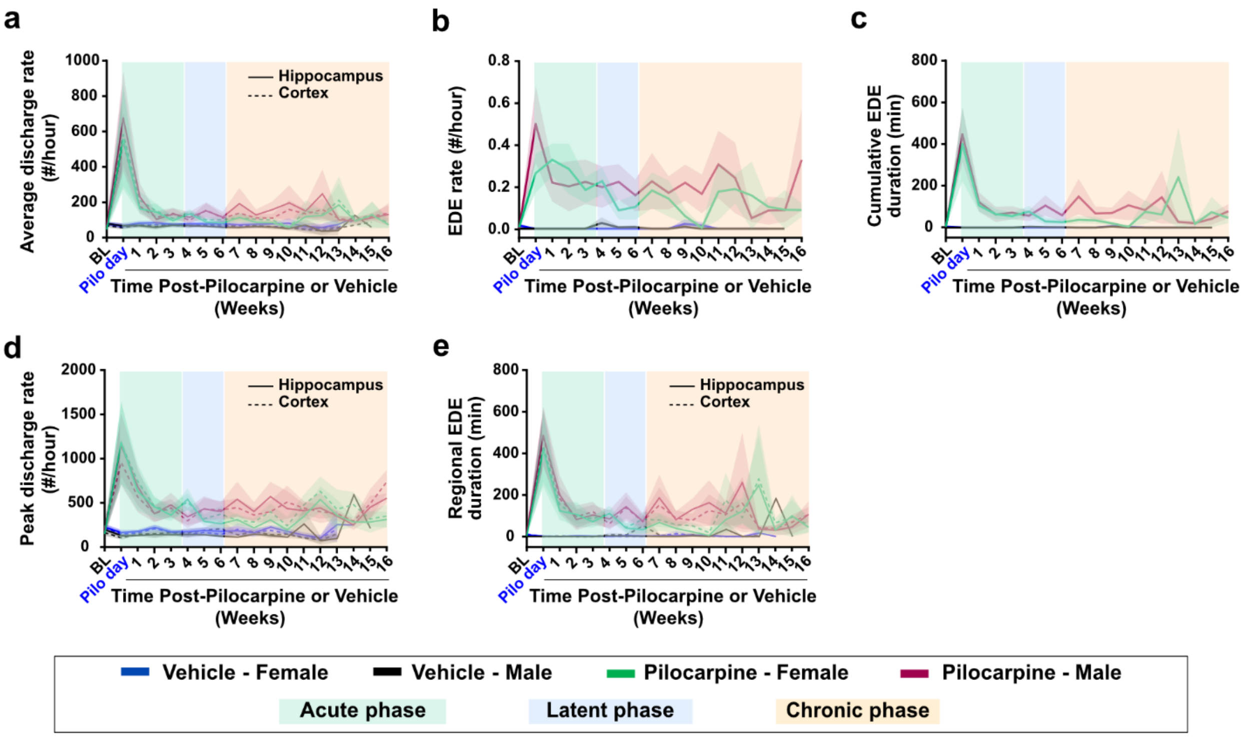


**Supplementary Figure 10 (relates to Fig. 5). EEG measures of pilocarpine-induced pathology, stratified by sex.**

**(a-e)** EEG metrics (same data as in **Fig. 5c-e** and **Supp**. **Fig. 7a-b**) stratified by sex. No consistent sex differences were detected; isolated timepoint differences are listed in **Supp. Table 9**. Vehicle: n=13 (7 male and 6 female) mice and pilocarpine: n=20 (11 male and 9 female) mice. Statistical analyses are the same as in **Fig. 5c-e** and **Supp. Fig. 7a-b.** Complete list of statistical comparisons and mouse numbers in each recording week are provided in **Supp. Table 9**.

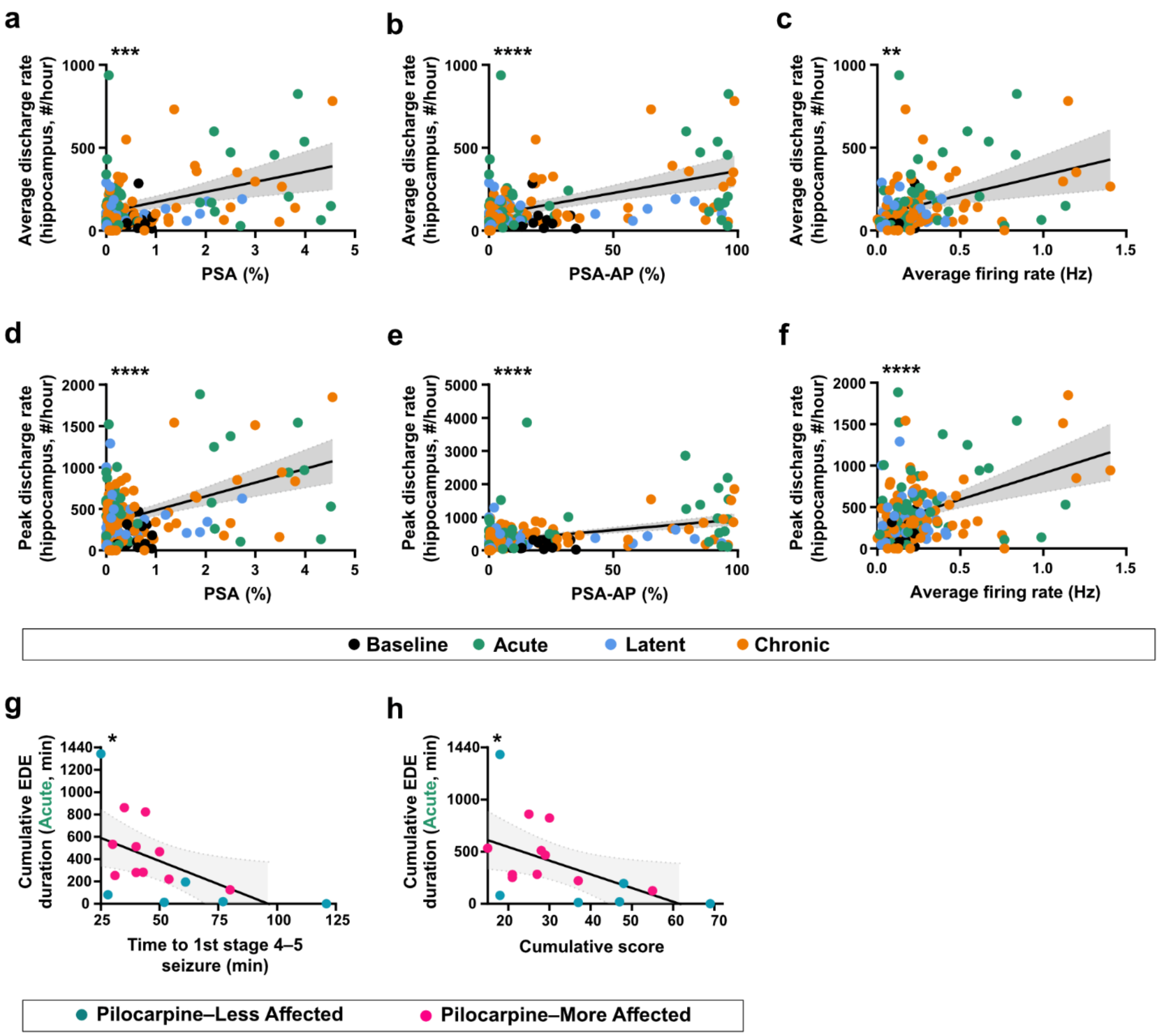


**Supplementary Figure 11 (relates to Fig. 5). Correlations between pilocarpine induction metrics, brain activity metrics, and hippocampal synchrony.**

**(a-c)** We found positive correlations between TPLSM-derived synchrony metrics or average firing rate and the average number of discharges recorded by the contralateral hippocampal EEG electrode.

**(d-f)** We found positive correlations between TPLSM-derived synchrony metrics or average firing rate and the maximal number of discharges recorded by the EEG electrode in the contralateral hippocampus.

For data in **(a–f)**, mouse numbers and statistical analyses are the same as in **Fig. 5h-m**. A complete list of statistical values is provided in **Supp. Table 10**. **, p<0.01; ***, p<0.001; ****, p<0.0001.

**(g-h)** Relationship between early behavioral seizure measures and subsequent EEG epileptiform burden: (n=16 mice; 8 male and 8 female). (**g**) Time to first seizure (Racine score 4 or 5) during the pilocarpine induction protocol is negatively correlated with the maximal cumulative EDE duration in a single EEG recording during the acute period. (**h**) Lower cumulative Racine score during the pilocarpine induction protocol (measured up to diazepam administration when applicable; see Methods) is associated with greater cumulative EDE duration during the acute period.

For data in (**g–h**), each dot represents the time to first seizure or cumulative Racine score for a single mouse paired with its maximal cumulative EDE duration during the acute period. The dots are colored according to the mouse severity group. Solid lines show the linear regression lines, shaded gray areas are the 95% confidence intervals. Statistical analyses were performed using single linear regression. $R^2$ and P-values for each fit are provided in **Supp. Table 10**. *,p<0.05.

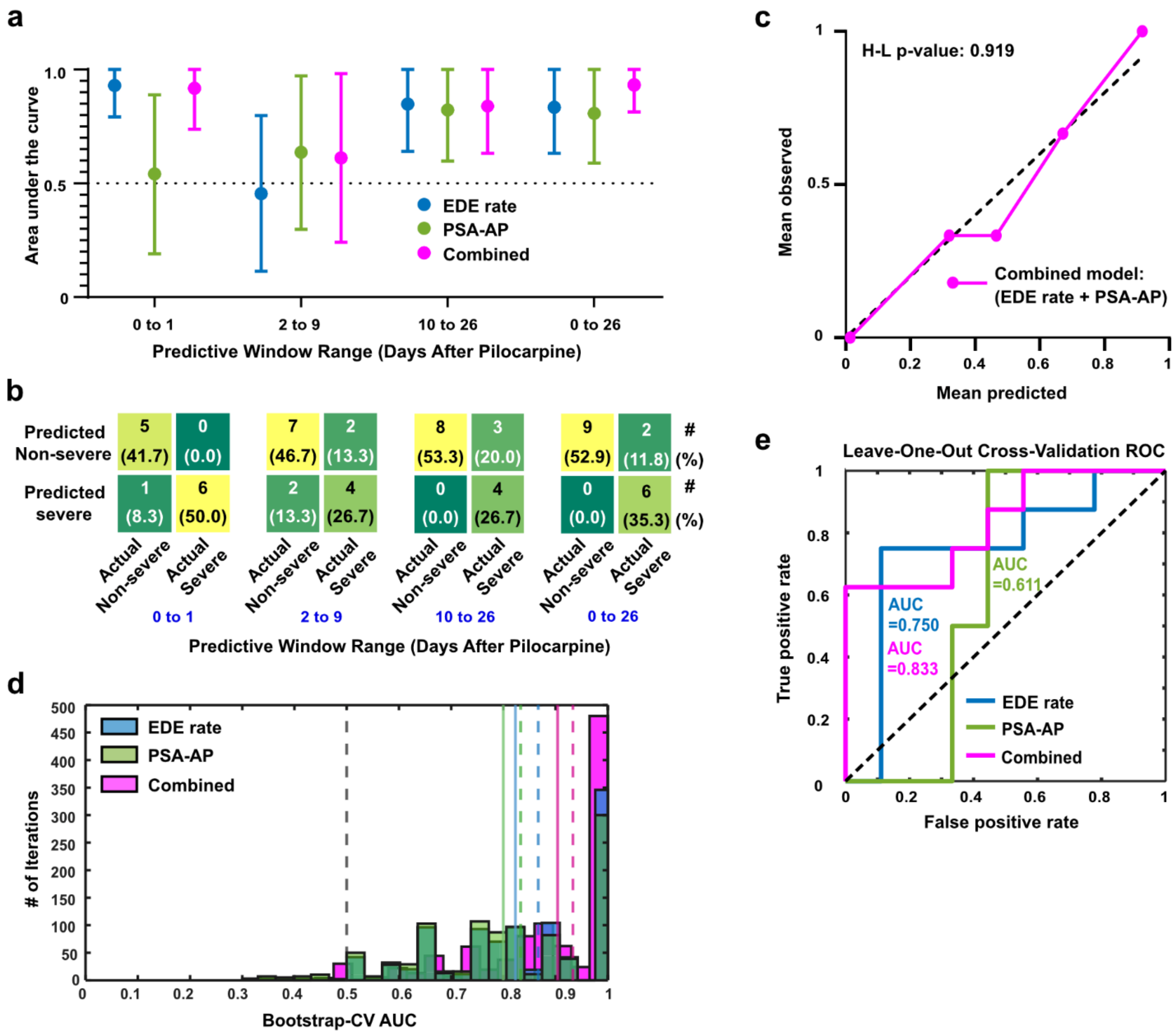


**Supplementary Figure 12 (relates to Fig. 6). Assessment of prediction accuracy.**

**(a)** Analysis of prediction quality using different acute-phase data ranges to predict the same-mouse chronic condition. Using days 0–1 (pilocarpine day and following day), PSA-AP-based prediction accuracy reduces to near-chance levels, while EDE-rate-based prediction is high. Using days 2–9 or 10–26, PSA-AP-based prediction improves in the later window, whereas EDE-rate-based prediction falls below that obtained from days 0–1 alone. Over the entire acute period, PSA-AP and EDE rate perform comparably. The combined model performs comparably to the better single-modality model in each restricted window and best over the full acute period. Dots show the calculated model AUC for ROC analysis; error bars indicate 95% confidence intervals using DeLong's method.

**(b)** Confusion matrices for each one of the analyzed cases in **a** using the combined model at the respective Youden optimal thresholds. Totals differ between windows (n=12, 15, 15 and 17 mice for the 0–1, 2–9, 10–26 and 0–26 day windows, respectively), reflecting the number of mice with usable recordings in each window.

**(c)** Hosmer-Lemeshow calibration plots for the combined multivariate model. Observed proportion of severe mice were plotted against the mean predicted probability. The dashed diagonal line demonstrates ideal calibration, (H-L p-value = 0.919 indicates adequate calibration).

**(d)** Bootstrap estimates of model performance (1,000 resamples). Histograms show out-of-bag AUCs: each model was fit on a bootstrap resample of the mice and evaluated on the mice not drawn into that resample (resamples whose out-of-bag set lacked either outcome class were redrawn). Dashed vertical lines indicate the mean out-of-bag AUC of each model; solid vertical lines indicate the bias-corrected AUC from Harrell's optimism correction (PSA-AP 0.800, EDE rate 0.827, combined 0.904); the grey dashed line indicates chance (AUC = 0.5). For the combined model, optimism was 0.026, giving a bias-corrected AUC of 0.904 from an apparent AUC of 0.931.

**(e)** Leave-one-out cross-validation shows that the EDE rate model generalizes better than the PSA-AP model. The combined model performs best (AUC 0.833).

### Supplementary Tables

**Supplementary Table 1. Summary of mouse numbers contributing to each analysis.**

| Cohort | Treatment group | n (mice) | Sex (M / F) | Criteria for inclusion | Figure reference |
|---|---|---|---|---|---|
| Total enrolled | Vehicle | 17 | 10 M / 7 F | All mice that received pilocarpine or vehicle injection | |
| | Pilocarpine | 37 | 23 M / 14 F | | Figure 1b |
| Refined Pilocarpine Group | Pilocarpine | 29 | 16 M / 13 F | Pilocarpine mice which survived the refined protocol | Figure 1c-d |
| TPLSM cohort | Vehicle | 10 | 5 M / 5 F | All surviving mice with at least one TPLSM recording session | Figure 2d-f, Supplementary Figure 3 |
| | Pilocarpine | 23 | 13 M / 10 F | | |
| PSA analysis | Pilocarpine | 14 | 8 M / 6 F | One recording session with at least 10 distinct PSA events | Figure 3, Supplementary Figure 5 |
| EEG cohort | Vehicle | 13 | 7 M / 6 F | All surviving mice with at least one EEG recording session | Figure 4, Figure 5c-e, Supplementary Figure 7, Supplementary Figure 10 |
| | Pilocarpine | 20 | 11 M / 9 F | | |
| Paired TPLSM + EEG cohort | Pilocarpine | 19 | 10 M / 9 F | Pilocarpine mice with at least one paired EEG and TPLSM recording session | Figure 5h-m, Supplementary Figure 11a-f |
| Acute-chronic correlation (EEG) | Pilocarpine | 19 | 10 M / 9 F | Pilocarpine mice with both acute and chronic phase EEG recording data | Figure 6a |
| Acute-chronic correlation (TPLSM) | Pilocarpine | 17 | 8 M / 9 F | Pilocarpine mice with both acute and chronic phase TPLSM recording data | Figure 6b |
| Predictive modeling | Pilocarpine | 17 (8 severe / 9 non-severe) | 8 M / 9 F | Pilocarpine mice with ≥2 TPLSM and ≥2 EEG sessions in the acute phase, ≥2 EEG sessions in the chronic phase | Figure 6c-e, Supplementary Figure 6, Supplementary Figure 12 |
| TPLSM cohort - more/less affected | Vehicle | 10 | 5 M / 5 F | All surviving mice with TPLSM imaging data across all three phases | Supplementary Figure 4 |
| | Pilocarpine | 17 | 8 M / 9 F | | |
| EEG cohort - more/less affected | Vehicle | 13 | 7 M / 6 F | All surviving mice with EEG and TPLSM data across all three phases | Figure 5f–g, Supplementary Figure 8, Supplementary Figure 9, Supplementary Figure 11g-h |
| | Pilocarpine | 16 | 8 M / 8 F | | |

Overview of all mice included in the study and how they were distributed across analyses. For each cohort, the table shows the treatment group (pilocarpine or vehicle), the total number of mice, the number of males and females, the criteria used to include an animal in that cohort, and the figures in which the cohort appears.

**Supplementary Table 2. Detailed statistical summary for Figure 2.**

| (i) F-statistics and corresponding p-values from mixed-effects model analyses - Figure 2d-f. | | | | | | |
|---|---|---|---|---|---|---|
| | F-test statistic | | | P-value | | |
| Panel | Time | Group | Time x Group (interaction) | Time | Group | Time x Group (interaction) |
| d | F (3.798, 68.78) = 0.5768 | F (1, 31) = 9.498 | F (3.798, 68.78) = 0.7285 | 0.6718 | 0.0043 | 0.5688 |
| e | F (4.355, 78.87) = 0.5179 | F (1, 31) = 4.353 | F (4.355, 78.87) = 0.6784 | 0.7378 | 0.0453 | 0.6212 |
| f | F (1.805, 43.32) = 4.975 | F (1, 31) = 6.762 | F (1.805, 43.32) = 3.320 | 0.0137 | 0.0141 | 0.0502 |

| (ii) Significant between-group comparisons at individual timepoints - Figure 2d-f. | | | |
|---|---|---|---|
| Panel | Timepoint/phase | Groups | Adjusted p-value |
| d | Day 3<br>Week 1<br>Week 9<br>Week 10 | Vehicle vs. Pilocarpine | 0.0056<br>0.0165<br>0.0089<br>0.0266 |
| e | Day 3<br>Week 3<br>Week 9 | Vehicle vs. Pilocarpine | 0.0014<br>0.0335<br>0.0470 |
| f | Acute<br>Chronic | Vehicle vs. Pilocarpine | 0.0001<br>0.0100 |

| (iii) Significant within-group comparisons across timepoints - Figure 2e-f. | | | |
|---|---|---|---|
| Panel | Group | Timepoint/Phase | Adjusted p-value |
| e | Vehicle | Pilo day vs. Day 3 | 0.0070 |
| f | Pilocarpine | Acute vs. Latent<br>Latent vs. Chronic | 0.0003<br>0.0478 |

| (iv) Number of mice remaining in longitudinal recordings at each subsequent time point - Figure 2d-e. | | | | | | | | | | | | | | | | | | | | |
|---|---|---|---|---|---|---|---|---|---|---|---|---|---|---|---|---|---|---|---|---|
| Timepoint | Baseline | Pilo day | Day 1 | Day 3 | Week 1 | Week 2 | Week 3 | Week 4 | Week 5 | Week 6 | Week 7 | Week 8 | Week 9 | Week 10 | Week 11 | Week 12 | Week 13 | Week 14 | Week 15 | Week 16 |
| Vehicle, n remaining | 10 | 10 | 10 | 10 | 7 | 7 | 7 | 7 | 7 | 7 | 7 | 7 | 7 | 6 | 5 | 5 | 3 | 2 | 2 | 0 |
| Pilocarpine, n remaining | 23 | 23 | 23 | 23 | 23 | 23 | 21 | 21 | 21 | 17 | 17 | 16 | 15 | 14 | 13 | 12 | 12 | 11 | 10 | 7 |

(i) Mixed-effects models were used to assess the statistical significance of the data shown in **Fig. 2d-f**. The table reports the F-statistics and corresponding p-values for the main effects of time and group, as well as their interaction. Statistically significant effects ($p < 0.05$) are highlighted in green.

(ii) Summary of the significant between-group differences between specific recording times ($p < 0.05$, Tukey HSD correction for multiple comparisons) for the data shown in **Fig. 2d–f**.

(iii) Summary of the significant within-group differences between specific recording times ($p < 0.05$, Tukey HSD correction for multiple comparisons) for the data shown in **Fig. 2e–f**.

(iv) Summary of the number of mice remaining in the study at each recording time.

### Supplementary Table 3. Detailed statistical summary for Figure 5.

**(i) F-statistics and corresponding p-values from mixed-effects model analyses - Figure5c-g.**

| | F-test statistic | | | P-value | | |
|---|---|---|---|---|---|---|
| Panel | Time | Group | Time x Group (interaction) | Time | Group | Time x Group (interaction) |
| c (Hippocampus) | F (1.540, 34.33) = 3.995 | F (1, 31) = 9.855 | F (1.540, 34.33) = 4.149 | 0.0371 | 0.0037 | 0.0334 |
| c (Cortex) | F (1.678, 37.02) = 4.525 | F (1, 31) = 16.00 | F (1.678, 37.02) = 4.892 | 0.0226 | 0.0004 | 0.0174 |
| d | F (3.638, 80.30) = 1.480 | F (1, 31) = 15.89 | F (3.638, 80.30) = 1.602 | 0.2197 | 0.0004 | 0.1867 |
| e | F (2.871, 63.36) = 4.134 | F (1, 31) = 18.52 | F (2.871, 63.36) = 4.196 | 0.0106 | 0.0002 | 0.0099 |
| f | F (2.706, 49.28) = 2.407 | F (2, 26) = 22.56 | F (5.411, 49.28) = 1.837 | 0.0843 | <0.0001 | 0.1178 |
| g | F (2.434, 44.34) = 6.276 | F (2, 26) = 16.29 | F (4.869, 44.34) = 2.384 | 0.0023 | <0.0001 | 0.0547 |

**(ii) Significant between-group comparisons at individual timepoints - Figure 5c-g.**

| Panel | Timepoint | Group | Adjusted p-value |
|---|---|---|---|
| c (Hippocampus) | Pilo day | Vehicle vs. Pilocarpine | 0.0110 |
| | Week 1 | | 0.0122 |
| | Week 3 | | 0.0349 |
| | Week 4 | | 0.0162 |
| | Week 8 | | 0.0374 |
| | Week 9 | | 0.0483 |
| c (Cortex) | Pilo day | Vehicle vs. Pilocarpine | 0.0077 |
| | Week 1 | | 0.0009 |
| | Week 2 | | 0.0243 |
| | Week 3 | | 0.0118 |
| | Week 4 | | 0.0009 |
| | Week 5 | | 0.0140 |
| | Week 6 | | 0.0495 |
| | Week 7 | | 0.0278 |
| | Week 8 | | 0.0104 |
| | Week 9 | | 0.0116 |
| | Week 11 | | 0.0051 |
| | Week 12 | | 0.0141 |
| d | Pilo day | Vehicle vs. Pilocarpine | 0.0025 |
| | Week 1 | | 0.0005 |
| | Week 2 | | 0.0079 |
| | Week 3 | | 0.0040 |
| | Week 4 | | 0.0040 |
| | Week 5 | | 0.0058 |
| | Week 6 | | 0.0151 |
| | Week 7 | | 0.0165 |
| | Week 8 | | 0.0112 |
| | Week 9 | | 0.0259 |
| | Week 11 | | 0.0352 |
| | Week 12 | | 0.0300 |
| e | Pilo day | Vehicle vs. Pilocarpine | 0.0015 |
| | Week 1 | | 0.0013 |
| | Week 2 | | 0.0093 |
| | Week 3 | | 0.0033 |
| | Week 4 | | 0.0028 |
| | Week 5 | | 0.0265 |
| | Week 6 | | 0.0300 |
| | Week 8 | | 0.0212 |
| | Week 9 | | 0.0242 |
| | Week 11 | | 0.0323 |
| f | Pilo Day | Vehicle vs. Pilocarpine-More Affected | 0.0107 |
| | Pilo Day | Pilocarpine-Less Affected vs. Pilocarpine-More Affected | 0.0190 |
| | Week 1 | Vehicle vs. Pilocarpine-More Affected | 0.0144 |
| | Week 2 | Vehicle vs. Pilocarpine-More Affected | 0.0344 |
| | Week 3 | Vehicle vs. Pilocarpine-More Affected | 0.0489 |
| | Week 5 | Vehicle vs. Pilocarpine-More Affected | 0.0306 |
| | Week5 | Pilocarpine-Less Affected vs. Pilocarpine-More Affected | 0.0338 |
| | Week 6 | Vehicle vs. Pilocarpine-More Affected | 0.0114 |
| | Week 6 | Pilocarpine-Less Affected vs. Pilocarpine-More Affected | 0.0127 |
| g | Pilo Day | Vehicle vs. Pilocarpine-More Affected | 0.0083 |
| | Week 1 | Vehicle vs. Pilocarpine-More Affected | 0.0163 |
| | Week 2 | Vehicle vs. Pilocarpine-More Affected | 0.0445 |
| | Week 3 | Vehicle vs. Pilocarpine-More Affected | 0.0228 |
| | Week 3 | Pilocarpine-Less Affected vs. Pilocarpine-More Affected | 0.0388 |

| (iii) Significant within-group comparisons across timepoints - Figure 5c-d. | | | |
|---|---|---|---|
| **Panel** | **Group** | **Timepoint** | **Adjusted p-value** |
| **c (Cortex)** | Vehicle | Week 6 vs. Week 13 | 0.0386 |
| **d** | Pilocarpine | Baseline vs. Week 1 | 0.0284 |

| (iv) Correlation analyses between TPLSM- and EEG-derived measures of hippocampal activity, synchronicity, and seizure severity - Figure 5h-m. | | | | |
|---|---|---|---|---|
| **Panel** | **Pearson $R^2$** | **Pearson p-value** | **Spearman ρ** | **Spearman p-value** |
| **h** | 0.1111 | <0.0001 | 0.1562 | 0.0337 |
| **i** | 0.1255 | <0.0001 | 0.2505 | 0.0006 |
| **j** | 0.1108 | <0.0001 | 0.3180 | <0.0001 |
| **k** | 0.0561 | 0.0012 | 0.1530 | 0.0377 |
| **l** | 0.0816 | <0.0001 | 0.2449 | 0.0008 |
| **m** | 0.0524 | 0.0017 | 0.3132 | <0.0001 |

| (v) Number of mice remaining in longitudinal recordings at each subsequent time point - Figure 5c-g. | | | | | | | | | | | | | | | | | | |
|---|---|---|---|---|---|---|---|---|---|---|---|---|---|---|---|---|---|---|
| **Timepoint** | **Baseline** | **Pilo day** | **Week 1** | **Week 2** | **Week 3** | **Week 4** | **Week 5** | **Week 6** | **Week 7** | **Week 8** | **Week 9** | **Week 10** | **Week 11** | **Week 12** | **Week 13** | **Week 14** | **Week 15** | **Week 16** |
| **(c-e) Vehicle, n remaining** | 13 | 13 | 12 | 12 | 12 | 12 | 12 | 12 | 11 | 10 | 9 | 8 | 4 | 4 | 4 | 1 | 1 | 0 |
| **(c-e) Pilocarpine, n remaining** | 20 | 20 | 20 | 20 | 20 | 20 | 20 | 19 | 19 | 18 | 17 | 14 | 13 | 13 | 13 | 11 | 10 | 7 |
| **(c-e) Vehicle, n remaining** | 13 | 13 | 12 | 12 | 12 | 12 | 12 | 12 | 11 | 10 | 9 | 8 | 4 | 4 | 4 | 2 | 1 | 0 |
| **(f-g) Pilocarpine - Less Affected n remaining** | 6 | 6 | 6 | 6 | 6 | 6 | 6 | 6 | 6 | 6 | 6 | 4 | 3 | 3 | 3 | 3 | 3 | 2 |
| **(f-g) Pilocarpine - More Affected n remaining** | 10 | 10 | 10 | 10 | 10 | 10 | 10 | 10 | 10 | 9 | 8 | 8 | 8 | 8 | 8 | 7 | 6 | 4 |

(i) Mixed-effects models were used to assess the statistical significance of the data shown in **Fig. 5c-g**. The table reports the F-statistics and corresponding p-values for the main effects of time and group, as well as their interaction. Statistically significant effects ($p < 0.05$) are highlighted in green.

(ii) Summary of the significant between-group differences between specific recording times ($p < 0.05$, Tukey HSD correction for multiple comparisons) for the data shown in **Fig. 5c–g**.

(iii) Summary of the significant within-group differences between specific recording times ($p < 0.05$, Tukey HSD correction for multiple comparisons) for the data shown in **Fig. 5c–g**.

(iv) Summary of goodness-of-fit for the data in **Fig. 5h-m,** based on Pearson and Spearman correlation coefficients and their p-values. All p-values are significant ($p < 0.05$).

(v) Summary of the number of mice remaining in the study at each recording time.

**Supplementary Table 4. Detailed statistical summary for Supplementary Figure 3.**

| (i) F-statistics and corresponding p-values from mixed-effects model analyses - Supplementary Figure 3a-e. | | | | | | |
|---|---|---|---|---|---|---|
| | F-test statistic | | | P-value | | |
| **Panel** | **Time** | **Group** | **Time x Group (interaction)** | **Time** | **Group** | **Time x Group (interaction)** |
| **a** | F (1.554, 37.28) = 3.118 | F (1, 31) = 9.695 | F (1.554, 37.28) = 2.483 | 0.0677 | 0.0040 | 0.1088 |
| **b** | F (1.900, 45.60) = 5.790 | F (1, 31) = 8.121 | F (1.900, 45.60) = 3.381 | 0.0065 | 0.0077 | 0.0450 |
| **c (Vehicle male vs. female)** | - | F (1, 8) = 0.01570 | - | 0.3833 | 0.9034 | 0.8485 |
| **c (Pilocarpine male vs. female)** | F (3.640, 44.83) = 1.405 | F (1, 21) = 0.3255 | F (3.640, 44.83) = 0.2321 | 0.2502 | 0.5744 | 0.9052 |
| **d (Vehicle male vs. female)** | - | F (1, 8) = 0.2993 | - | 0.5277 | 0.5993 | 0.8337 |
| **d (Pilocarpine male vs. female)** | F (3.879, 47.77) = 1.329 | F (1, 21) = 0.04679 | F (3.879, 47.77) = 0.4669 | 0.2734 | 0.8308 | 0.7541 |
| **e (Vehicle male vs. female)** | - | F (1, 8) = 0.6780 | - | 0.6343 | 0.4341 | 0.8098 |
| **e (Pilocarpine male vs. female)** | F (3.856, 47.49) = 1.090 | F (1, 21) = 0.003805 | F (3.856, 47.49) = 0.3817 | 0.3713 | 0.9514 | 0.8140 |

| (ii) Significant between-group comparisons across disease phases - Supplementary Figure 3a-b. | | | |
|---|---|---|---|
| **Panel** | **Phase** | **Group** | **Adjusted p-value** |
| **a** | Acute | Vehicle vs. Pilocarpine | 0.0003 |
| | Chronic | Vehicle vs. Pilocarpine | 0.0042 |
| **b** | Acute | Vehicle vs. Pilocarpine | <0.0001 |
| | Chronic | Vehicle vs. Pilocarpine | 0.0091 |

| (iii) Significant within-group comparisons across phases - Supplementary Figure 3a-b. | | | |
|---|---|---|---|
| **Panel** | **Group** | **Phase** | **Adjusted p-value** |
| **a** | Pilocarpine | Acute vs. Latent | 0.0025 |
| | Pilocarpine | Latent vs. Chronic | 0.0163 |
| **b** | Pilocarpine | Acute vs. Latent | 0.0002 |

| (iv) Number of mice remaining in longitudinal recordings at each subsequent time point - Supplementary Figure 3c-e. | | | | | | | | | | | | | | | | | | | |
|---|---|---|---|---|---|---|---|---|---|---|---|---|---|---|---|---|---|---|---|
| **Timepoint** | **Baseline** | **Pilo day** | **Day 1** | **Day 3** | **Week 1** | **Week 2** | **Week 3** | **Week 4** | **Week 5** | **Week 6** | **Week 7** | **Week 8** | **Week 9** | **Week 10** | **Week 11** | **Week 12** | **Week 13** | **Week 14** | **Week 15** | **Week 16** |
| **Vehicle - Male, n remaining** | 5 | 5 | 5 | 5 | 4 | 4 | 4 | 4 | 4 | 4 | 4 | 4 | 4 | 3 | 2 | 2 | 1 | 1 | 1 | 0 |
| **Vehicle - Female, n remaining** | 5 | 5 | 5 | 5 | 3 | 3 | 3 | 3 | 3 | 3 | 3 | 3 | 3 | 3 | 3 | 3 | 2 | 1 | 1 | 0 |
| **Pilocarpine - Male, n remaining** | 13 | 13 | 13 | 13 | 13 | 13 | 11 | 11 | 11 | 8 | 8 | 8 | 8 | 7 | 7 | 6 | 6 | 5 | 5 | 3 |
| **Pilocarpine - Female, n remaining** | 10 | 10 | 10 | 10 | 10 | 10 | 10 | 10 | 10 | 9 | 9 | 8 | 7 | 7 | 6 | 6 | 6 | 6 | 5 | 4 |

(i) Mixed-effects models were used to assess the statistical significance of the data shown in **Supplementary Fig. 3a–e**. F-statistics and corresponding p-values for the main effects of time and group, as well as their interaction. For panels c–e, sex-based comparisons were analyzed separately within the Vehicle and Pilocarpine groups; for some subgroup analyses, degrees of freedom could not be estimated due to insufficient data points, and only p-values are reported for these comparisons. Statistically significant effects ($p < 0.05$) are highlighted in green.

(ii) Summary of the significant between-group differences across disease phases ($p < 0.05$, Tukey HSD correction for multiple comparisons) for the data shown in **Supplementary Fig. 3a–b**.

(iii) Summary of the significant within-group differences across disease phases ($p < 0.05$, Tukey HSD correction for multiple comparisons) for the data shown in **Supplementary Fig. 3a–b.**

(iv) Summary of the number of mice remaining in the study at each recording time, separated by sex, for the data shown in **Supplementary Fig. 3c–e.**

### Supplementary Table 5. Statistical summary for Supplementary Figure 4.

**(i) F-statistics and corresponding p-values from ordinary one-way ANOVA - Supplementary Figure 4c-d.**

| Panel | F-statistic | P- value |
|---|---|---|
| c | F(2, 24) = 13.25 | 0.0001 |
| d | F(2, 24) = 13.47 | 0.0001 |

**(iii) Significant between-group post hoc comparisons - Supplementary Figure 4c-j.**

| Panel | Phase/Timepoint | Group | Adjusted p-value |
|---|---|---|---|
| c | Overall | Pilocarpine-Less Affected vs. Pilocarpine-More Affected | 0.0016 |
| | | Pilocarpine-More Affected vs. Vehicle | 0.0002 |
| d | Overall | Pilocarpine-Less Affected vs. Pilocarpine-More Affected | 0.0009 |
| | | Pilocarpine-More Affected vs. Vehicle | 0.0003 |
| e | Acute | Vehicle vs. Pilocarpine-More Affected | 0.0025 |
| | Chronic | Vehicle vs. Pilocarpine-More Affected | 0.0090 |
| | Chronic | Pilocarpine-More Affected vs. Pilocarpine-Less Affected | 0.0175 |
| f | Acute | Vehicle vs. Pilocarpine-More Affected | <0.0001 |
| | Chronic | Vehicle vs. Pilocarpine-More Affected | 0.0105 |
| | | Pilocarpine-More Affected vs. Pilocarpine-Less Affected | 0.0114 |
| g | Acute | Pilocarpine-Less Affected vs. Pilocarpine-More Affected | 0.0243 |
| | Acute | Pilocarpine-More Affected vs. Vehicle | <0.0001 |
| | Chronic | Pilocarpine-Less Affected vs. Pilocarpine-More Affected | 0.0069 |
| | Chronic | Pilocarpine-More Affected vs. Vehicle | 0.0056 |
| h | Week 9 | Vehicle vs. Pilocarpine-More Affected | 0.0293 |
| | Week 14 | Pilocarpine-More Affected vs. Vehicle | 0.0081 |
| | Week 14 | Pilocarpine-More Affected vs. Pilocarpine- Less Affected | 0.0234 |
| i | Week 9 | Vehicle vs. Pilocarpine-Less Affected | 0.0459 |
| | | Pilocarpine-More Affected vs. Pilocarpine- Less Affected | 0.0338 |
| j | Pilo day | Vehicle vs. Pilocarpine- Less Affected | 0.0457 |
| | Day 3 | Vehicle vs. Pilocarpine-More Affected | 0.0174 |
| | Day 3 | Pilocarpine-More Affected vs. Pilocarpine- Less Affected | 0.0457 |
| | Week 9 | Vehicle vs. Pilocarpine-More Affected | 0.0456 |
| | Week 9 | Pilocarpine-More Affected vs. Pilocarpine- Less Affected | 0.0249 |

**(ii) F-statistics and corresponding p-values from mixed-effects model analyses - Supplementary Figure 4e-j.**

| | F-test statistic | | | P-value | | |
|---|---|---|---|---|---|---|
| Panel | Time | Group | Time x Group (interaction) | Time | Group | Time x Group (interaction) |
| e | F (1.390, 29.19) = 5.820 | F (2, 24) = 10.27 | F (2.780, 29.19) = 2.483 | 0.0142 | 0.0006 | 0.0845 |
| f | F (1.675, 35.17) = 9.690 | F (2, 24) = 14.80 | F (3.350, 35.17) = 3.249 | 0.0008 | <0.0001 | 0.0290 |
| g | F (1.906, 40.02) = 9.672 | F (2, 24) = 21.07 | F (3.811, 40.02) = 3.426 | 0.0005 | <0.0001 | 0.0182 |
| h | F (4.026, 61.73) = 0.6654 | F (2, 24) = 9.668 | F (8.052, 61.73) = 1.273 | 0.6194 | 0.0008 | 0.2737 |
| i | F (3.694, 56.65) = 0.6892 | F (2, 24) = 10.28 | F (7.389, 56.65) = 1.124 | 0.5914 | 0.0006 | 0.3614 |
| j | F (3.693, 63.19) = 1.156 | F (2, 30) = 14.07 | F (7.385, 63.19) = 1.391 | 0.3377 | <0.0001 | 0.2220 |

**(iv) Significant within-group post hoc comparisons - Supplementary Figure 4e-j.**

| Panel | Group | Timepoint/Phase | Adjusted p-value |
|---|---|---|---|
| e | Pilocarpine-More Affected | Acute vs. Latent | 0.0073 |
| | | Latent vs. Chronic | 0.0097 |
| f | Pilocarpine-More Affected | Acute vs. Latent | 0.0005 |
| g | Pilocarpine-More Affected | Acute vs. Latent | 0.0029 |
| h | Pilocarpine- Less Affected | Week 7 vs. Week 15 | 0.0306 |
| | | Week 9 vs. Week 11 | 0.044 |
| | | Week 12 vs. Week 15 | 0.0212 |
| i | vehicle | Week 14 vs. Week 15 | 0.044 |
| | Pilocarpine-More Affected | Pilo day vs. Week 14 | 0.0302 |
| | Pilocarpine-Less Affected | Week 4 vs. Week 13 | 0.0211 |
| j | Vehicle | Pilo day vs. Day 3 | 0.007 |
| | Pilocarpine- Less Affected | Baseline vs. Week 5 | 0.032 |
| | Pilocarpine-More Affected | Pilo day vs. Week 15 | 0.2228 |

| (v) Number of mice remaining in longitudinal recordings at each subsequent time point - Supplementary Figure 4h-j. | | | | | | | | | | | | | | | | | | | | |
|---|---|---|---|---|---|---|---|---|---|---|---|---|---|---|---|---|---|---|---|---|
| **Timepoint** | **Baseline** | **Pilo day** | **Day 1** | **Day 3** | **Week 1** | **Week 2** | **Week 3** | **Week 4** | **Week 5** | **Week 6** | **Week 7** | **Week 8** | **Week 9** | **Week 10** | **Week 11** | **Week 12** | **Week 13** | **Week 14** | **Week 15** | **Week 16** |
| **Vehicle, n remaining** | 10 | 10 | 10 | 10 | 7 | 7 | 7 | 7 | 7 | 7 | 7 | 7 | 7 | 6 | 5 | 5 | 3 | 2 | 2 | 0 |
| **Pilocarpine - Less Affected, n remaining** | 7 | 7 | 7 | 7 | 7 | 7 | 7 | 7 | 7 | 7 | 7 | 7 | 6 | 5 | 4 | 4 | 4 | 4 | 3 | 2 |
| **Pilocarpine - More Affected, n remaining** | 10 | 10 | 10 | 10 | 10 | 10 | 10 | 10 | 10 | 10 | 10 | 9 | 9 | 9 | 9 | 8 | 8 | 7 | 7 | 5 |

(i) Ordinary one-way ANOVA was used to assess the statistical significance of the data shown in **Supplementary Fig. 4c–d**. The table reports the F-statistic and corresponding p-value for each panel. Statistically significant effects ($p < 0.05$) are highlighted in green.

(ii) Mixed-effects models were used to assess the statistical significance of the data shown in **Supplementary Fig. 4e–j.** The table reports the F-statistics and corresponding p-values for the main effects of time and group, as well as their interaction. Statistically significant effects ($p < 0.05$) are highlighted in green.

(iii) Summary of the significant between-group differences at specific phases/timepoints ($p < 0.05$, Tukey HSD correction for multiple comparisons) for the data shown in **Supplementary Fig. 4c–j.**

(iv) Summary of the significant within-group differences across specific phases/timepoints ($p < 0.05$, Tukey HSD correction for multiple comparisons) for the data shown in **Supplementary Fig. 4e–j.**

(v) Summary of the number of mice remaining in the study at each recording time, for the data shown in **Supplementary Fig. 4e–j.**

**Supplementary Table 6. Detailed statistical summary for Supplementary Figure 7.**

| (i) F-statistics and corresponding p-values from mixed-effects model analyses - Supplementary Figure 7a-j. | | | | | | |
|---|---|---|---|---|---|---|
| | F-test statistic | | | p-value | | |
| Panel | Time | Group | Time x Group (interaction) | Time | Group | Time x Group (interaction) |
| a (Hippocampus) | F (3.142, 70.03) = 2.919 | F (1, 31) = 21.17 | F (3.142, 70.03) = 3.441 | 0.0378 | <0.0001 | 0.0197 |
| a (Cortex) | F (3.819, 84.30) = 2.350 | F (1, 31) = 38.15 | F (3.819, 84.30) = 3.154 | 0.0635 | <0.0001 | 0.0197 |
| b (Hippocampus) | F (2.883, 64.25) = 2.786 | F (1, 31) = 13.83 | F (2.883, 64.25) = 2.880 | 0.0498 | 0.0008 | 0.0446 |
| b (Cortex) | F (3.543, 78.19) = 3.493 | F (1, 31) = 22.47 | F (3.543, 78.19) = 3.831 | 0.0144 | <0.0001 | 0.0090 |
| c | F (1.640, 46.74) = 2.957 | F (1, 31) = 19.36 | F (1.640, 46.74) = 4.457 | 0.0715 | 0.0001 | 0.0229 |
| d | F (1.548, 44.11) = 3.283 | F (1, 31) = 16.64 | F (1.548, 44.11) = 3.478 | 0.0586 | 0.0003 | 0.0508 |
| e | F (1.142, 32.54) = 3.490 | F (1, 31) = 8.464 | F (1.142, 32.54) = 3.953 | 0.0658 | 0.0066 | 0.0503 |
| f | F (1.096, 31.23) = 3.150 | F (1, 31) = 10.90 | F (1.096, 31.23) = 3.624 | 0.0825 | 0.0024 | 0.0628 |
| g | F (1.246, 35.52) = 3.322 | F (1, 31) = 20.99 | F (1.246, 35.52) = 4.367 | 0.0683 | <0.0001 | 0.0360 |
| h | F (1.301, 37.07) = 2.035 | F (1, 31) = 30.69 | F (1.301, 37.07) = 3.141 | 0.1583 | <0.0001 | 0.0744 |
| i | F (1.681, 47.91) = 3.372 | F (1, 31) = 13.95 | F (1.681, 47.91) = 3.419 | 0.0505 | 0.0008 | 0.0486 |
| j | F (1.647, 46.94) = 2.182 | F (1, 31) = 19.89 | F (1.647, 46.94) = 3.892 | 0.1325 | 0.0001 | 0.0347 |

| (ii) Significant between-group comparisons at individual timepoints or across disease phases - Supplementary Figure 7 a-j. | | | |
|---|---|---|---|
| **Panel** | **Timepoint/Phase** | **Group** | **Adjusted p-value** |
| **a (Hippocampus)** | Pilo day | Vehicle vs. Pilocarpine | 0.0031 |
| | Week 1 | | 0.0035 |
| | Week 2 | | 0.0057 |
| | Week 3 | | 0.0034 |
| | Week 4 | | 0.0026 |
| | Week 5 | | 0.0184 |
| | Week 6 | | 0.0100 |
| | Week 7 | | 0.0199 |
| | Week 8 | | 0.0206 |
| | Week 9 | | 0.0163 |
| | Week 12 | | 0.0047 |
| **a (Cortex)** | Pilo day | Vehicle vs. Pilocarpine | 0.0027 |
| | Week 1 | | 0.0014 |
| | Week 2 | | 0.0033 |
| | Week 3 | | 0.0010 |
| | Week 4 | | 0.0022 |
| | Week 5 | | 0.0105 |
| | Week 6 | | 0.0167 |
| | Week 7 | | 0.0057 |
| | Week 8 | | 0.0078 |
| | Week 9 | | 0.0115 |
| | Week 11 | | 0.0001 |
| | Week 12 | | 0.0014 |
| | Week 13 | | 0.0476 |
| **b (Hippocampus)** | Pilo day | Vehicle vs. Pilocarpine | 0.0011 |
| | Week 1 | | 0.0090 |
| | Week 2 | | 0.0246 |
| | Week 3 | | 0.0066 |
| | Week 4 | | 0.0029 |
| | Week 5 | | 0.0308 |
| | Week 6 | | 0.0281 |
| | Week 8 | | 0.0239 |
| | Week 9 | | 0.0273 |
| **b (Cortex)** | Pilo day | Vehicle vs. Pilocarpine | 0.0009 |
| | Week 1 | | 0.0003 |
| | Week 2 | | 0.0085 |
| | Week 3 | | 0.0013 |
| | Week 4 | | 0.0004 |
| | Week 5 | | 0.0089 |
| | Week 7 | | 0.0254 |
| | Week 8 | | 0.0083 |
| | Week 9 | | 0.0065 |
| | Week 11 | | 0.0306 |
| **c** | Acute | Vehicle vs. Pilocarpine | <0.0001 |
| | Latent | | 0.0005 |
| | Chronic | | 0.0001 |
| **d** | Acute | Vehicle vs. Pilocarpine | 0.0003 |
| | Latent | | 0.0020 |
| | Chronic | | 0.0033 |
| **e** | Acute | Vehicle vs. Pilocarpine | 0.0042 |
| | Latent | | 0.0116 |
| | Chronic | | 0.0069 |
| **f** | Acute | Vehicle vs. Pilocarpine | 0.0040 |
| | Latent | | 0.0019 |
| | Chronic | | 0.0004 |
| **g** | Acute | Vehicle vs. Pilocarpine | 0.0002 |
| | Latent | | 0.0003 |
| | Chronic | | 0.0009 |
| **h** | Acute | Vehicle vs. Pilocarpine | 0.0003 |
| | Latent | | 0.0014 |
| | Chronic | | <0.0001 |
| **i** | Acute | Vehicle vs. Pilocarpine | 0.0001 |
| | Latent | | 0.0022 |
| | Chronic | | 0.0071 |
| **j** | Acute | Vehicle vs. Pilocarpine | 0.0001 |
| | Latent | | 0.0051 |
| | Chronic | | 0.0009 |

| (iii) Significant within-group post hoc comparisons across disease phases - Supplementary Figure 7c-j. | | | |
|---|---|---|---|
| **Panel** | **Group** | **Phase** | **Adjusted p-value** |
| **c** | Pilocarpine | Acute vs. Latent | 0.0202 |
| | | Latent vs. Chronic | 0.0100 |
| **d** | Pilocarpine | Acute vs. Latent | 0.0154 |
| **e** | Pilocarpine | Acute vs. Latent | 0.0364 |
| | | Latent vs. Chronic | 0.0288 |
| **f** | Pilocarpine | Acute vs. Latent | 0.0405 |
| **g** | Pilocarpine | Acute vs. Latent | 0.0160 |
| **i** | Pilocarpine | Acute vs. Latent | 0.0114 |
| **j** | Pilocarpine | Acute vs. Latent | 0.0160 |

| (iv) Number of mice remaining in longitudinal recordings at each subsequent time point - Supplementary Figure 7a-b. | | | | | | | | | | | | | | | | | | |
|---|---|---|---|---|---|---|---|---|---|---|---|---|---|---|---|---|---|---|
| **Timepoint** | **Baseline** | **Pilo day** | **Week 1** | **Week 2** | **Week 3** | **Week 4** | **Week 5** | **Week 6** | **Week 7** | **Week 8** | **Week 9** | **Week 10** | **Week 11** | **Week 12** | **Week 13** | **Week 14** | **Week 15** | **Week 16** |
| **Vehicle, n remaining** | 13 | 13 | 12 | 12 | 12 | 12 | 12 | 12 | 11 | 10 | 9 | 8 | 4 | 4 | 4 | 1 | 1 | 0 |
| **Pilocarpine, n remaining** | 20 | 20 | 20 | 20 | 20 | 20 | 20 | 19 | 19 | 18 | 17 | 14 | 13 | 13 | 13 | 11 | 10 | 7 |

(i) Mixed-effects models were used to assess the statistical significance of the data shown in **Supplementary Fig. 7a–j**. The table reports the F-statistics and corresponding p-values for the main effects of time and group, as well as their interaction. Statistically significant effects ($p < 0.05$) are highlighted in green.

(ii) Summary of the significant between-group differences at specific timepoints or disease phases ($p < 0.05$, Tukey HSD correction for multiple comparisons) for the data shown in **Supplementary Fig. 7a–j**.

(iii) Summary of the significant within-group differences across disease phases ($p < 0.05$, Tukey HSD correction for multiple comparisons) for the data shown in **Supplementary Fig. 7c–j**.

(iv) Summary of the number of mice remaining in the study at each recording time for the data shown in **Supplementary Fig. 7a–b**.

### Supplementary Table 7. Detailed statistical summary for Supplementary Figure 8.

**(i) F-statistics and corresponding p-values from mixed-effects model analyses - Supplementary Figure 8a-c.**

| | F-test statistic | | | p-value | | |
|---|---|---|---|---|---|---|
| **Panel** | **Time** | **Group** | **Time x Group (interaction)** | **Time** | **Group** | **Time x Group (interaction)** |
| **a (Hippocampus)** | F (1.482, 26.18) = 6.321 | F (2, 26) = 7.331 | F (2.964, 26.18) = 2.406 | 0.0102 | 0.0030 | 0.0905 |
| **a (Cortex)** | F (1.627, 29.64) = 8.787 | F (2, 26) = 11.41 | F (3.254, 29.64) = 3.163 | 0.0018 | 0.0003 | 0.0357 |
| **b (Hippocampus)** | F (2.818, 49.78) = 5.837 | F (2, 26) = 20.12 | F (5.635, 49.78) = 3.067 | 0.0021 | <0.0001 | 0.0140 |
| **b (Cortex)** | F (3.244, 59.08) = 6.268 | F (2, 26) = 28.37 | F (6.487, 59.08) = 3.044 | 0.0007 | <0.0001 | 0.0099 |
| **c (Hippocampus)** | F (2.587, 45.71) = 4.150 | F (2, 26) = 12.26 | F (5.174, 45.71) = 1.914 | 0.0146 | 0.0002 | 0.1082 |
| **c (Cortex)** | F (2.981, 54.30) = 5.267 | F (2, 26) = 17.34 | F (5.963, 54.30) = 2.315 | 0.0030 | <0.0001 | 0.0465 |

**(ii) Significant between-group post hoc comparisons at individual timepoints - Supplementary Figure 8a-c.**

| **Panel** | **Timepoint** | **Group** | **Adjusted p-value** |
|---|---|---|---|
| **a (Cortex)** | Pilo Day | Vehicle vs. Pilocarpine-More Affected | 0.0144 |
| | Week 1 | Vehicle vs. Pilocarpine-More Affected | 0.0034 |
| | Week 2 | Vehicle vs. Pilocarpine-More Affected | 0.0391 |
| | Week 4 | Vehicle vs. Pilocarpine-More Affected | 0.0260 |
| | Week 5 | Vehicle vs. Pilocarpine-More Affected | 0.0235 |
| | Week 5 | Pilocarpine-Less Affected vs. Pilocarpine-More Affected | 0.0320 |
| | Week 8 | Vehicle vs. Pilocarpine-More Affected | 0.0301 |
| | Week 9 | Vehicle vs. Pilocarpine-More Affected | 0.0048 |
| | Week 11 | Vehicle vs. Pilocarpine-More Affected | 0.0478 |
| **b (Hippocampus)** | Pilo Day | Vehicle vs. Pilocarpine-More Affected | 0.0124 |
| | Week 1 | Pilocarpine-More Affected vs. Vehicle | 0.0165 |
| | Week 1 | Pilocarpine-More Affected vs. Pilocarpine-Less Affected | 0.0483 |
| | Week 1 | Vehicle vs. Pilocarpine-Less Affected | 0.0378 |
| | Week 2 | Vehicle vs. Pilocarpine-More Affected | 0.0189 |
| | Week 3 | Vehicle vs. Pilocarpine-More Affected | 0.0144 |
| | Week 3 | Pilocarpine-Less Affected vs. Pilocarpine-More Affected | 0.0330 |
| | Week 6 | Vehicle vs. Pilocarpine-More Affected | 0.0394 |
| | Week 12 | Vehicle vs. Pilocarpine-More Affected | 0.0265 |
| **b (Cortex)** | Pilo Day | Vehicle vs. Pilocarpine-More Affected | 0.0046 |
| | Week 1 | Vehicle vs. Pilocarpine-More Affected | 0.0186 |
| | Week 2 | Vehicle vs. Pilocarpine-More Affected | 0.0097 |
| | Week 3 | Vehicle vs. Pilocarpine-More Affected | 0.0050 |
| | Week 4 | Vehicle vs. Pilocarpine-Less Affected | 0.0477 |
| | Week 5 | Pilocarpine-More Affected vs. Vehicle | 0.0111 |
| | Week 5 | Pilocarpine-More Affected vs. Pilocarpine-Less Affected | 0.0184 |
| | Week 11 | Vehicle vs. Pilocarpine-More Affected | 0.0058 |
| | Week 12 | Vehicle vs. Pilocarpine-More Affected | 0.0004 |
| **c (Hippocampus)** | Pilo Day | Vehicle vs. Pilocarpine-More Affected | 0.0063 |
| **c (Cortex)** | Pilo Day | Vehicle vs. Pilocarpine-More Affected | 0.0046 |
| | Week 1 | Vehicle vs. Pilocarpine-More Affected | 0.0053 |
| | Week 2 | Vehicle vs. Pilocarpine-More Affected | 0.0497 |
| | Week 3 | Vehicle vs. Pilocarpine-More Affected | 0.0410 |
| | Week 5 | Pilocarpine-More Affected vs. Vehicle | 0.0327 |
| | Week 5 | Pilocarpine-More Affected vs. Pilocarpine-Less Affected | 0.0334 |
| | Week 9 | Vehicle vs. Pilocarpine-More Affected | 0.0447 |

**(iii) Number of mice remaining in longitudinal recordings at each subsequent time point - Supplementary Figure 8.**

| **Timepoint** | **Baseline** | **Pilo day** | **Week 1** | **Week 2** | **Week 3** | **Week 4** | **Week 5** | **Week 6** | **Week 7** | **Week 8** | **Week 9** | **Week 10** | **Week 11** | **Week 12** | **Week 13** | **Week 14** | **Week 15** | **Week 16** |
|---|---|---|---|---|---|---|---|---|---|---|---|---|---|---|---|---|---|---|
| **Vehicle, n remaining** | 13 | 13 | 12 | 12 | 12 | 12 | 12 | 12 | 11 | 10 | 9 | 8 | 4 | 4 | 4 | 2 | 1 | 0 |
| **Pilocarpine - Less Affected n remaining** | 6 | 6 | 6 | 6 | 6 | 6 | 6 | 6 | 6 | 6 | 6 | 4 | 3 | 3 | 3 | 3 | 3 | 2 |
| **Pilocarpine - More Affected n remaining** | 10 | 10 | 10 | 10 | 10 | 10 | 10 | 10 | 10 | 9 | 8 | 8 | 8 | 8 | 8 | 7 | 6 | 4 |

(i) Mixed-effects models were used to assess the statistical significance of the data shown in **Supplementary Fig. 8a–c.** The table reports the F-statistics and corresponding p-values for the main effects of time and group, as well as their interaction. Statistically significant effects ($p < 0.05$) are highlighted in green.

(ii) Summary of the significant between-group differences at specific timepoints ($p < 0.05$, Tukey HSD correction for multiple comparisons) for the data shown in **Supplementary Fig. 8a–c**.

(iii) Summary of the number of mice remaining in the study at each recording time for the data shown in **Supplementary Fig. 8a–c**.

**Supplementary Table 8. Detailed statistical summary for Supplementary Figure 9.**

| (i) F-statistics and corresponding p-values from mixed-effects model analyses - Supplementary Figure 9a-h. | | | | | | |
|---|---|---|---|---|---|---|
| | F-test statistic | | | p-value | | |
| Panel | Time | Group | Time x Group (interaction) | Time | Group | Time x Group (interaction) |
| a | F (1.760, 42.23) = 7.674 | F (2, 26) = 21.44 | F (3.519, 42.23) = 4.596 | 0.0021 | <0.0001 | 0.0050 |
| b | F (1.505, 36.12) = 5.376 | F (2, 26) = 12.46 | F (3.010, 36.12) = 2.101 | 0.0150 | 0.0002 | 0.1170 |
| c | F (1.105, 26.52) = 6.093 | F (2, 26) = 5.875 | F (2.210, 26.52) = 2.297 | 0.0179 | 0.0078 | 0.1158 |
| d | F (1.091, 26.18) = 6.174 | F (2, 26) = 6.158 | F (2.182, 26.18) = 2.048 | 0.0176 | 0.0065 | 0.1459 |
| e | F (1.212, 29.09) = 6.153 | F (2, 26) = 18.05 | F (2.424, 29.09) = 3.482 | 0.0146 | <0.0001 | 0.0363 |
| f | F (1.232, 45.59) = 4.615 | F (2, 74) = 18.26 | F (2.464, 45.59) = 2.077 | 0.0298 | <0.0001 | 0.1270 |
| g | F (1.587, 38.09) = 5.603 | F (2, 26) = 12.27 | F (3.175, 38.09) = 2.761 | 0.0116 | 0.0002 | 0.0524 |
| h | F (1.639, 39.34) = 3.997 | F (2, 26) = 12.92 | F (3.278, 39.34) = 2.201 | 0.0335 | 0.0001 | 0.0981 |

| (ii) Significant between-group post hoc comparisons across disease phases - Supplementary Figure9a-h. | | | |
|---|---|---|---|
| Panel | Phase | Group | Adjusted p-value |
| a | Acute | Vehicle vs. Pilocarpine-More Affected | 0.0003 |
| | Latent | Vehicle vs. Pilocarpine-More Affected | 0.0081 |
| | Chronic | Vehicle vs. Pilocarpine-More Affected | 0.0051 |
| | Chronic | Vehicle vs. Pilocarpine-Less Affected | 0.0188 |
| | Chronic | Pilocarpine-More Affected vs. Pilocarpine-Less Affected | 0.0308 |
| b | Acute | Vehicle vs. Pilocarpine-More Affected | 0.0010 |
| | Chronic | Vehicle vs. Pilocarpine-More Affected | 0.0493 |
| | Chronic | Vehicle vs. Pilocarpine-Less Affected | 0.0109 |
| c | Acute | Vehicle vs. Pilocarpine-More Affected | 0.0224 |
| | Chronic | Vehicle vs. Pilocarpine-More Affected | 0.0450 |
| d | Acute | Vehicle vs. Pilocarpine-More Affected | 0.0081 |
| | Latent | Vehicle vs. Pilocarpine-More Affected | 0.0126 |
| | Chronic | Vehicle vs. Pilocarpine-More Affected | 0.0176 |
| | Chronic | Vehicle vs. Pilocarpine-Less Affected | 0.0364 |
| e | Acute | Vehicle vs. Pilocarpine-More Affected | 0.0002 |
| | Latent | Vehicle vs. Pilocarpine-More Affected | 0.0036 |
| | Chronic | Vehicle vs. Pilocarpine-More Affected | 0.0204 |
| | Chronic | Vehicle vs. Pilocarpine-Less Affected | 0.0087 |
| f | Acute | Vehicle vs. Pilocarpine-More Affected | 0.0004 |
| | Latent | Vehicle vs. Pilocarpine-More Affected | 0.0058 |
| | Chronic | Vehicle vs. Pilocarpine-More Affected | 0.0031 |
| | Chronic | Vehicle vs. Pilocarpine-Less Affected | 0.0107 |
| g | Acute | Vehicle vs. Pilocarpine-More Affected | 0.0002 |
| | Chronic | Vehicle vs. Pilocarpine-More Affected | 0.0495 |
| h | Acute | Vehicle vs. Pilocarpine-More Affected | 0.0009 |
| | Latent | Vehicle vs. Pilocarpine-More Affected | 0.0272 |
| | Chronic | Vehicle vs. Pilocarpine-More Affected | 0.0405 |
| | Chronic | Vehicle vs. Pilocarpine-Less Affected | 0.0430 |

| (iii) Significant within-group post hoc comparisons across disease phases - Supplementary Figure 9a-h. | | | |
|---|---|---|---|
| Panel | Group | Phase | Adjusted p-value |
| a | Pilocarpine-More Affected | Acute vs. Latent | 0.0047 |
| a | Pilocarpine-More Affected | Latent vs. Chronic | 0.0379 |
| b | Pilocarpine-More Affected | Acute vs. Latent | 0.0094 |
| c | Pilocarpine-More Affected | Latent vs. Chronic | 0.0317 |
| d | Pilocarpine-More Affected | Acute vs. Latent | 0.0223 |
| e | Pilocarpine-More Affected | Acute vs. Latent | 0.0058 |
| | Pilocarpine-More Affected | Acute vs. Chronic | 0.0434 |
| | Pilocarpine-More Affected | Latent vs. Chronic | 0.0051 |
| f | Pilocarpine-More Affected | Acute vs. Latent | 0.0175 |
| g | Pilocarpine-More Affected | Acute vs. Latent | 0.0030 |
| h | Pilocarpine-More Affected | Acute vs. Latent | 0.0050 |

(i) Mixed-effects models were used to assess the statistical significance of the data shown in **Supplementary Fig. 9a–h**. The table reports the F-statistics and corresponding p-values for the main effects of time and group, as well as their interaction. Statistically significant effects ($p < 0.05$) are highlighted in green.

(ii) Summary of the significant between-group differences across disease phases ($p < 0.05$, Tukey HSD correction for multiple comparisons) for the data shown in **Supplementary Fig. 9a–h**.

(iii) Summary of the significant within-group differences across disease phases ($p < 0.05$, Tukey HSD correction for multiple comparisons) for the data shown in **Supplementary Fig. 9a–h**.

**Supplementary Table 9. Detailed statistical summary for Supplementary Figure 10.**

| (i) F-statistics and corresponding p-values from mixed-effects model analyses - Supplementary Figure 10a-e. | | | | | | |
|---|---|---|---|---|---|---|
| | F-test statistic | | | p-value | | |
| Panel | Time | Group | Time x Group (interaction) | Time | Group | Time x Group (interaction) |
| a (Pilocarpine male vs. female, Hippocampus) | F (1.556, 18.49) = 5.089 | F (1, 18) = 0.1856 | F (1.556, 18.49) = 0.3359 | 0.0236 | 0.6717 | 0.6656 |
| a (Vehicle male vs. female, Hippocampus) | F (0.7955, 5.739) = 0.7510 | F (1, 11) = 0.2171 | F (0.7955, 5.739) = 0.6089 | 0.3891 | 0.6504 | 0.4289 |
| a (Pilocarpine male vs. female, Cortex) | F (1.558, 18.43) = 5.914 | F (1, 18) = 0.2324 | F (1.558, 18.43) = 0.3009 | 0.0149 | 0.6356 | 0.6897 |
| a (Vehicle male vs. female, Cortex) | F (2.858, 20.21) = 0.5702 | F (1, 11) = 0.009148 | F (2.858, 20.21) = 0.3769 | 0.6331 | 0.9255 | 0.7614 |
| b (Pilocarpine male vs. female) | F (3.334, 39.41) = 2.128 | F (1, 18) = 0.02346 | F (3.334, 39.41) = 0.7716 | 0.1061 | 0.8800 | 0.5291 |
| b (Vehicle male vs. female) | F (1.904, 13.47) = 0.6461 | F (1, 11) = 0.02610 | F (1.904, 13.47) = 0.7103 | 0.5323 | 0.8746 | 0.5025 |
| c (Pilocarpine male vs. female) | F (2.936, 34.71) = 5.487 | F (1, 18) = 0.01985 | F (2.936, 34.71) = 0.8700 | 0.0036 | 0.8895 | 0.4640 |
| c (Vehicle - Male vs. Vehicle - Female) | F (2.000, 14.14) = 1.165 | F (1, 11) = 0.01809 | F (2.000, 14.14) = 0.6472 | 0.3401 | 0.8954 | 0.5383 |
| d (Pilocarpine male vs. female, Hippocampus) | F (3.062, 36.39) = 4.030 | F (1, 18) = 0.2679 | F (3.062, 36.39) = 0.4038 | 0.0138 | 0.6110 | 0.7551 |
| d (Vehicle male vs. female, Hippocampus) | F (2.131, 15.37) = 1.280 | F (1, 11) = 1.269 | F (2.131, 15.37) = 1.088 | 0.3081 | 0.2839 | 0.3653 |
| d (Pilocarpine male vs. female, Cortex) | F (3.645, 43.09) = 3.560 | F (1, 18) = 0.06459 | F (3.645, 43.09) = 0.6638 | 0.0159 | 0.8023 | 0.6071 |
| d (Vehicle male vs. female, Cortex) | F (4.154, 29.37) = 0.5851 | F (1, 11) = 0.3954 | F (4.154, 29.37) = 0.4534 | 0.6819 | 0.5423 | 0.7757 |
| e (Pilocarpine male vs. female, Hippocampus) | F (3.205, 38.08) = 3.723 | F (1, 18) = 0.08854 | F (3.205, 38.08) = 0.8366 | 0.0174 | 0.7694 | 0.4887 |
| e (Vehicle male vs. female, Hippocampus) | F (1.478, 10.67) = 1.081 | F (1, 11) = 0.03524 | F (1.478, 10.67) = 1.617 | 0.3524 | 0.8545 | 0.2401 |
| e (Pilocarpine male vs. female, Cortex) | F (3.591, 42.46) = 4.906 | F (1, 18) = 0.1070 | F (3.591, 42.46) = 0.9389 | 0.0033 | 0.7473 | 0.4433 |
| e (Vehicle male vs. female, Cortex) | F (1.253, 9.845) = 0.5414 | F (1, 110) = 0.5944 | F (1.253, 9.845) = 0.6285 | 0.5181 | 0.4424 | 0.4817 |

| (ii) Significant between-group post hoc comparisons at individual timepoints - Supplementary Figure 10d. | | | |
|---|---|---|---|
| Panel | Timepoint | Group | Adjusted p-value |
| d (Hippocampus) | Week 9 | Vehicle male vs. female | 0.0285 |

| (iii) Number of mice remaining in longitudinal recordings at each subsequent time point - Supplementary Figure 10a-e. | | | | | | | | | | | | | | | | | | |
|---|---|---|---|---|---|---|---|---|---|---|---|---|---|---|---|---|---|---|
| Timepoint | Baseline | Pilo day | Week 1 | Week 2 | Week 3 | Week 4 | Week 5 | Week 6 | Week 7 | Week 8 | Week 9 | Week 10 | Week 11 | Week 12 | Week 13 | Week 14 | Week 15 | Week 16 |
| Vehicle - Male, n remaining | 7 | 7 | 7 | 7 | 7 | 7 | 7 | 7 | 6 | 6 | 5 | 5 | 2 | 2 | 2 | 1 | 1 | 0 |
| Vehicle - Female, n remaining | 6 | 6 | 5 | 5 | 5 | 5 | 5 | 5 | 5 | 4 | 4 | 3 | 2 | 2 | 2 | 1 | 0 | 0 |
| Pilocarpine - Male, n remaining | 11 | 11 | 11 | 11 | 11 | 11 | 11 | 10 | 10 | 9 | 9 | 8 | 8 | 8 | 8 | 6 | 5 | 3 |
| Pilocarpine - Female, n remaining | 9 | 9 | 9 | 9 | 9 | 9 | 9 | 9 | 9 | 9 | 8 | 6 | 5 | 5 | 5 | 5 | 5 | 4 |

(i) Mixed-effects models were used to assess the statistical significance of the data shown in **Supplementary Fig. 10a–e**, comparing male and female mice within each treatment group. The table reports the F-statistics and corresponding p-values for the main effects of time and group (sex), as well as their interaction. No consistent sex differences were detected; isolated timepoint differences are listed in **(ii)**. Statistically significant effects ($p < 0.05$) are highlighted in green.

(ii) Summary of the significant between-group differences at specific timepoints ($p < 0.05$, Tukey HSD correction for multiple comparisons) for the data shown in **Supplementary Fig. 10d.**

(iii) Summary of the number of mice remaining in the study at each recording time, separated by treatment group and sex, for the data shown in **Supplementary Fig. 10a–e**.

**Supplementary Table 10. Detailed statistical summary for correlations shown in Supplementary Figure 11.**

| (i) Correlation analyses between TPLSM- and EEG-derived measures of hippocampal activity, synchronicity, and seizure severity - Supplementary Figure 11a-f. | | | | |
|---|---|---|---|---|
| **Panel** | **Pearson $R^2$** | **Pearson p-value** | **Spearman $\rho$** | **Spearman p-value** |
| **a** | 0.0938 | 0.0008 | 0.1857 | 0.0097 |
| **b** | 0.0938 | <0.0001 | 0.3269 | <0.0001 |
| **c** | 0.0492 | 0.0023 | 0.3490 | <0.0001 |
| **d** | 0.1110 | <0.0001 | 0.1607 | 0.0256 |
| **e** | 0.1481 | <0.0001 | 0.2779 | <0.0001 |
| **f** | 0.0939 | <0.0001 | 0.3199 | <0.0001 |

| (ii) Correlation analyses between induction-day behavioral severity measures (time to first stage 4-5 seizure and cumulative racine score) and acute-phase EEG cumulative seizure duration - Supplementary Figure 11g-h. | | |
|---|---|---|
| **Panel** | **Pearson $R^2$** | **Pearson p-value** |
| **g** | 0.2977 | 0.0235 |
| **h** | 0.2799 | 0.0290 |

(i) Pearson correlation ($R^2$ and p-values) and Spearman rank correlation ($\rho$ and p-values) were used to assess the relationship between TPLSM- and EEG-derived measures of hippocampal activity, synchrony, and epileptiform burden for the data shown in **Supplementary Fig. 11a–f**. Statistically significant values ($p < 0.05$) are highlighted in green.

(ii) Linear regression ($R^2$ and p-values) was used to assess the relationship between pilocarpine-induction measures (time to first stage 4–5 seizure; cumulative Racine score) and maximal acute-phase cumulative EDE duration for the data shown in **Supplementary Fig. 11g–h.** Statistically significant values ($p < 0.05$) are highlighted in green.

**Supplementary Table 11. EDEs containing seizure-criterion segments under three published electrographic seizure definitions, and confirmation on blinded expert review.**

| Criterion | Botterill et al. (>3 Hz for >5 s) | Jain et al. (≥3 Hz for ≥10 s) | ACNS (>2.5 Hz averaged over 10 s) | ≥3 Hz for ≥10 s, confirmed on review |
|---|---|---|---|---|
| **All post-induction EDEs, n/932 (%)[a]** | 87 (9.3) | 68 (7.3) | 86 (9.2) | 48 (5.2) |
| **Induction day, n/87** | 7 | 5 | 5 | 3 |
| **Acute, n/264 (%); mice** | 8 (3.0); 7/18 | 4 (1.5); 4/18 | 8 (3.0); 7/18 | 4 (1.5); 4/18 |
| **Latent, n/182 (%); mice** | 6 (3.3); 2/17 | 6 (3.3); 2/17 | 6 (3.3); 2/17 | 6 (3.3); 2/17 |
| **Chronic, n/399 (%); mice** | 66 (16.5); 12/18 | 53 (13.3); 11/18 | 67 (16.8); 12/18 | 35 (8.8); 9/18 |
| **Mice with ≥1 such EDE, of 19** | 16 | 13 | 14 | 11 |
| **Control EDEs, n/14[b]** | 2 | 0 | 1 | 0 |
| **Segment duration, s, median (range); n measurable[c]** | 26 (9–79); 71 | 26 (10–79); 68 | 26 (9–79); 71 | 29 (11–79); 48 |
| **Electrodes: both / hippocampus only / cortex only** | 35 / 18 / 34 | 30 / 17 / 21 | 39 / 15 / 32 | 25 / 8 / 15 |
| **Review windows scored as seizure, n/flagged[d]** | 43/70 | 37/52 | 43/71 | — |

[a] Denominator 932 is all EDEs recorded after pilocarpine administration; the phase rows sum to it.

[b] Control EDEs are the 14 detected in vehicle-treated recordings and in pre-pilocarpine baselines.

[c] The segment is the longest stretch in which the discharge rate, computed in a 10-s sliding window, stayed at or above 3 Hz. The Botterill criterion (a 5-s window above 3 Hz) and the ACNS criterion (a 10-s average above 2.5 Hz) can be met without the 10-s rate ever reaching 3 Hz; those EDEs have no segment to measure and are excluded from the segment-duration row. The Jain criterion is defined on the same basis, so every EDE it flags has a measurable segment.

[d] An expert epileptologist blinded to group, phase and analysis output scored 77 flagged windows (flagged events within 45 s shown once) and 25 control windows from EDEs meeting no criterion; 1 of 25 control windows was scored as a seizure. An EDE was counted as confirmed if a window containing one of its flagged events was scored as a seizure.

**Supplementary Table 12. Univariate screening of acute-phase predictors of chronic severity.**

| Predictor | Severe median (IQR) | Non-severe median (IQR) | Wilcoxon p | Firth OR (95% CI) | Firth p | AUC |
|---|---|---|---|---|---|---|
| Firing rate, Hz | 0.68 (0.55–0.78) | 0.30 (0.22–0.99) | 0.361 | 1.256 (0.161–10.623) | 0.820 | 0.639 |
| PSA, % | 3.76 (2.55–3.89) | 0.63 (0.37–4.32) | 0.312 | 1.526 (0.883–2.929) | 0.134 | 0.653 |
| PSA-AP | 93.72 (92.43–95.96) | 26.80 (13.41–93.46) | 0.039 | 1.039 (1.007–1.133) | 0.015 | 0.806 |
| EDE rate, events/h | 0.88 (0.54–0.96) | 0.27 (0.06–0.33) | 0.024 | 36.615 (2.155–1940.375) | 0.010 | 0.833 |
| Cumulative EDE duration, min | 435.92 (274.95–605.81) | 80.67 (13.53–280.18) | 0.061 | 1.001 (0.999–1.004) | 0.320 | 0.778 |

n = 17 mice (8 severe, 9 non-severe). Values are maximal acute-phase (pilocarpine day to Week 3) measures, shown as median (IQR); group differences by Wilcoxon rank-sum test. Odds ratios are from univariate Firth logistic regression, per 1 Hz (firing rate), per percentage point (PSA%, PSA-AP), per EDE/h (EDE rate) or per minute (cumulative EDE duration); AUC is from the corresponding ROC curve.

### **Supplementary Video Legends**

**Supplementary Video 1. Longitudinal TPLSM recordings of the same CA1 region**

TPLSM recordings (30 fps, 3-frame running average) of the same CA1 FOV at baseline, 1 day, 3 days, 6 weeks, 7 weeks, and 13 weeks post-pilocarpine. FOV size: 600×600 $\mu m^2$, 512×512 pixels.

**Supplementary Video 2. TPLSM propagation example recording**

TPLSM recording (30 fps, raw) of CA1 region at 10 weeks post-pilocarpine induction showing PSA propagation from posterior to anterior (600×600 $\mu m^2$, 512×512 pixels).

**Supplementary Video 3. Behavioral seizures during EDEs**

Examples of behavioral seizures occurring within EDEs, with vEEG signals during daytime and nighttime recordings.
